# Structure and function of aerotolerant, multiple-turnover THI4 thiazole synthases

**DOI:** 10.1101/2021.08.03.453570

**Authors:** Jaya Joshi, Qiang Li, Jorge D. García-García, Bryan J. Leong, You Hu, Steven D. Bruner, Andrew D. Hanson

## Abstract

Plant and fungal THI4 thiazole synthases produce the thiamin thiazole moiety in aerobic conditions via a single-turnover suicide reaction that uses an active-site Cys residue as sulfur donor. Multiple-turnover (i.e. catalytic) THI4s lacking an active-site Cys (non-Cys THI4s) that use sulfide as sulfur donor have been characterized – but only from archaeal methanogens that are anaerobic, O_2_-sensitive hyperthermophiles from sulfide-rich habitats. These THI4s prefer iron as cofactor. A survey of prokaryote genomes uncovered non-Cys THI4s in aerobic mesophiles from sulfide-poor habitats, suggesting that multiple-turnover THI4 operation is possible in aerobic, mild, low-sulfide conditions. This was confirmed by testing 23 representative non-Cys THI4s for complementation of an *Escherichia coli* Δ*thiG* thiazole auxotroph in aerobic conditions. Sixteen were clearly active, and more so when intracellular sulfide level was raised by supplying Cys, demonstrating catalytic function in the presence of O_2_ at mild temperatures and indicating use of sulfide or a sulfide metabolite as sulfur donor. Comparative genomic evidence linked non-Cys THI4s with proteins from families that bind, transport, or metabolize cobalt or other heavy metals. The crystal structure of the aerotolerant bacterial *Thermovibrio ammonificans* THI4 was determined to probe the molecular basis of aerotolerance. The structure suggested no large deviations compared to the structures of THI4s from O_2_-sensitive methanogens, but is consistent with an alternative catalytic metal. Together with complementation data, the use of cobalt rather than iron was supported. We conclude that catalytic THI4s can indeed operate aerobically and that the metal cofactor inserted is a likely natural determinant of aerotolerance.

## INTRODUCTION

Biosynthesis of the adenylated carboxythiazole (ADT) precursor of thiamin is chemically complex and energetically expensive [1,2]. Plants, fungi, and some prokaryotes make ADT via the thiazole synthase THI4, a single-turnover suicide enzyme [3–6]. In a reaction requiring iron (yeast) or zinc (Arabidopsis), these THI4s form ADT from NAD, glycine, and a sulfur atom stripped from an active-site Cys residue [3,5,7,8]. The sulfur loss converts Cys to dehydroalanine and irreversibly inactivates the enzyme [3,5] (Figure 1). Such THI4s must therefore be replaced after just one reaction cycle, and this – plus the high demand for thiazole [9] – makes THI4 one of the shortest-lived proteins in plant leaves [10,11].

**Figure 1.**
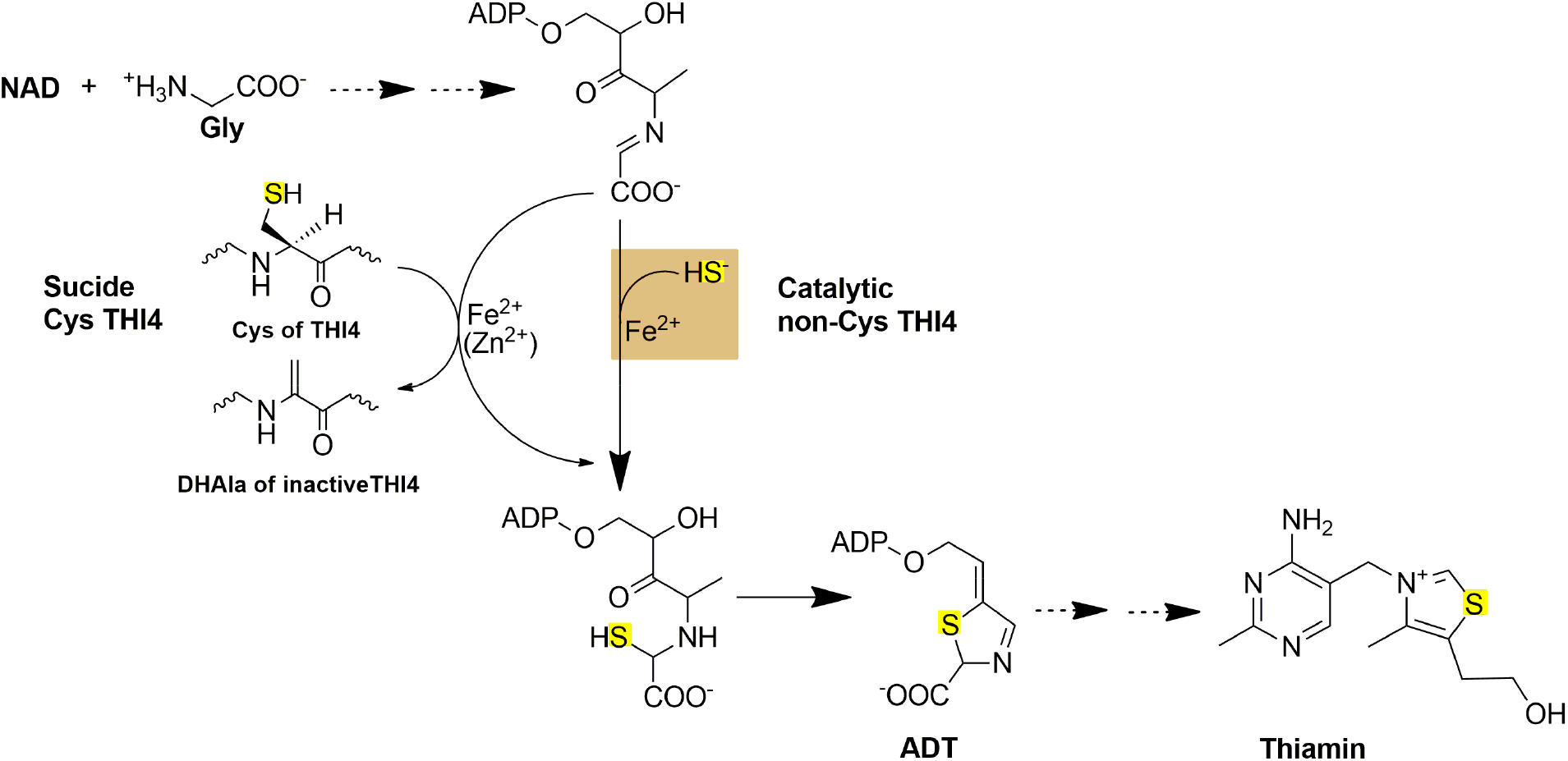
Biosynthesis of the thiazole precursor of thiamin by suicide and catalytic THI4s. THI4s form the adenylated carboxythiazole (ADT) precursor of thiamin from NAD, glycine, and a sulfur atom that in yeast and plant THI4s comes from an active-site cysteine residue and in methanococcal THI4s comes from sulfide (HS^-^). Sulfur loss from the active-site cysteine leaves a dehydroalanine (DHAla) residue that is not reconverted to cysteine, making yeast and plant THI4s suicide enzymes. In contrast, THI4s that use sulfide mediate multiple turnovers, i.e. are true catalysts.

Bioenergetic calculations indicate that the cost of THI4 degradation and resynthesis in plants reduces biomass accumulation by 2-4% [2]. Crop biomass gains of this order could therefore result from engineered replacement of a suicide THI4 with a catalytic THI4 that, like most enzymes, mediates thousands of turnovers in its lifetime [12,13]. But do catalytic THI4s that can operate in the mild, aerobic conditions typical of plant cells exist in nature? And if so, what characteristics confer this ability?

Catalytic THI4s have been reported from strictly anaerobic, O_2_-sensitive, thermophilic methanogens from hydrothermal vents, where sulfide levels are high enough (millimolar) to kill plants and most other organisms [7,14]. These THI4s use sulfide as the sulfur donor, have His in place of Cys in the active site, and prefer iron as cofactor *in vitro* (Figure 1) [7,14]. An exploratory survey [15] of prokaryote genomes identified THI4s with no active-site Cys (non-Cys THI4s) in several diverse organisms, and preliminary tests showed that two such THI4s complemented an *Escherichia coli* thiazole synthase (Δ*thiG*) mutant in aerobic conditions [15]. By indicating that non-CysTHI4s can have at least some activity in mild conditions in the presence of O_2_, these pilot data prompted further research.

In this work we deeply surveyed the diversity and genomic contexts of prokaryote non-Cys THI4s and ran complementation assays of thiazole synthase activity on a representative set. In addition, the crystal structure of a THI4 with aerobic complementing activity was determined and this THI4’s *in vivo* metal preference was explored. The results implicated the cofactor metal as a determinant of O_2_-tolerance.

## MATERIALS AND METHODS

### Chemicals and media

Chemicals and reagents were from Sigma-Aldrich or Fisher Scientific unless otherwise indicated. MOPS minimal medium was prepared as described [16] except that it was supplemented with the concentrations of micronutrients as specified in [17].

### Bioinformatics

Microbial THI4 sequences were identified in the SEED [18] and UniRef90 [19] databases using *Thermovibrio ammonificans* THI4 as query sequence. Comparative genomics analyses were performed using SEED and GenBank resources. Sequence similarity networks (SSNs) were constructed by submitting 199 THI4 sequences to the EFI-EST webtool using the FASTA option [20]. An E value of 10^-5^ was used to delimit sequence similarity. A final SSN was generated with an alignment score cutoff set such that each connection (edge) represents ~80% sequence identity. At this setting, some sequences remained as singletons. Network layouts were created and visualized using Cytoscape 3.4.

### Knockout strain and clone construction

An *E. coli* MG1655 Δ*thiG* strain was made by recombineering [15,21] using the Δ*thiG* cassette from the corresponding Keio collection strain [22]. Selected THI4 genes were recoded for expression in *E. coli* or yeast and synthesized by GenScript (Piscataway, NJ) or Twist Biosciences (San Francisco, CA). Recoded nucleotide sequences are given in Supplementary Table 1. For *E. coli*, recoded sequences with an added N-terminal His_6_ tag were cloned between the EcoRI and XbaI sites in pBAD24 [23]. For yeast (*Saccharomyces cerevisiae*), the recoded *Thermovibrio ammonificans* THI4 sequence (preceded by the putative yeast THI4 targeting peptide MSATSTATSTSASQLHLNSTPVTHCLSDGG plus a GG linker) or the native yeast THI4 was inserted into the *CEN6/ARS4* nuclear plasmid carrying the *his3* marker and the TDH3 promoter to drive THI4 expression.

### THI4 protein expression analysis

pBAD24 constructs were introduced into the *E. coli* MG1655 Δ*thiG* strain and single colonies were used to inoculate 3 ml of MOPS medium containing 0.2% (w/v) glycerol, 100 nM thiamin, 100 μg/ml ampicillin and 50 μg/ml kanamycin. The next day, 25-ml cultures were grown at 37°C in MOPS-glycerol medium with 100 nM thiamin supplementation until OD_600nm_ reached 0.8. Cells were then induced by adding 0.02% (w/v) arabinose and incubated for another 4 h at 37°C. Cells were harvested by centrifugation (6 000***g***, 15 min, 4°C), flash-frozen in liquid nitrogen, and stored at −80°C. Cell pellets were extracted by sonicating in 1 ml of 100 mM potassium phosphate (pH 7.2) containing 2 mM β-mercapto-ethanol, and separated into soluble and insoluble fractions by centrifugation (17 000***g***, 10 min, 4°C). Proteins in the pellet were solubilized by boiling for 5 min in 0.5 ml of SDS sample buffer. Aliquots (10 μl) of the insoluble fraction extract or tenfold-diluted soluble protein extract were separated by SDS-PAGE on 15% gels; proteins were detected by Coomassie Blue staining. The identity of the THI4 bands was confirmed by immunoblotting using anti-His_6_ tag antibodies (Thermo Fisher Scientific MA1-21315). The THI4 Coomassie band in soluble and insoluble fractions was quantified using Licor Image Studio Lite software. A 3441-pixel area around the band was used to calculate signal intensity. The method was calibrated using a standard curve for purified recombinant *T. ammonificans* THI4.

### Functional complementation assays in E. coli and yeast

For assays in *E. coli*, three independent clones of each construct were used to inoculate 3 ml of MOPS medium containing 0.2% glycerol (w/v), 100 nM thiamin, and 50 μg/ml kanamycin. After incubation at 37°C for 18 h, cells were harvested by centrifugation, washed five times with thiamin-free MOPS medium, resuspended in 500 μl of the same medium, serially diluted in ten-fold steps, and spotted on MOPS medium plates containing 0.2% glycerol, 0.02% arabinose, plus or minus 1 mM Cys. Plates were then incubated at 37°C in aerobic and near-anerobic (N_2_ containing ~1 ppm O_2_) conditions as described [15]. Images were captured after 7 d. For complementation assays with *T. ammonificans* THI4 in yeast, three independent clones of strain Δ*THI4* BY4741 (*MATa his3*Δ*1 leu2*Δ*0 met15*Δ*0 ura3*Δ*0*; *THI4*Δ::KanMX) transformed with the *CEN6/ARS4* plasmid alone or containing *T. ammonificans* THI4 or yeast THI4 were used to inoculate 3 ml of synthetic, complete medium (SC; yeast nitrogen base, USBiological cat. no. Y2036), drop-out mix (USBiological cat. no. D9540) minus histidine supplemented with 20 g/l glucose, 5 g/l ammonium sulfate, and 300 nM thiamin. After 48 h of incubation at 30 °C and 220 rpm, cells were pelleted (3 000***g***, 5 min), washed five times with thiamin-free SC minus histidine medium, resuspended in the same medium, and used to inoculate 3 ml of thiamin-free SC minus histidine medium to an OD_600nm_ of 0.05. Growth was then monitored at OD_600nm_.

### Purification and anaerobic reconstitution of T. ammonificans THI4

*T. ammonificans* THI4 cloned in pBAD24 with an N-terminal His_6_-tag was transformed into *E. coli* BL21(DE3). A starter culture (15 ml LB plus 100 μg/ml ampicillin) was inoculated into 6 l of LB; THI4 expression was initiated at an OD_600_ of 0.6 with arabinose (0.02% w/v) and incubation was continued at 22°C for 20 h. Cells were then harvested by centrifugation; pellets were resuspended in 50 ml of lysis buffer (20 mM Tris-HCl, pH 8.0, 500 mM NaCl, 2 mM 2-mercaptoethanol) and lysed in a microfluidizer cell (14,000 psi, M-110L Pneumatic). The lysate was clarified by centrifugation (18 000***g***, 30 min, 4°C) and applied to a 1-ml Ni-NTA column (HisPur, ThermoFisher Scientific). After incubation at 4°C for 1 h the resin was washed with 50 ml of lysis buffer and eluted with 6 ml lysis buffer plus 250 mM imidazole. The eluate was dialyzed against 1 l of 20 mM Tris-HCl, pH 8.0, 100 mM NaCl, 2 mM 2-mercaptoethanol for 12 h and purified by anion exchange chromatography (HiTrap Q, GE Healthcare) with a linear gradient of 0 to 500 mM NaCl over 40 min and size-exclusion chromatography (HiLoad 16/600 Superdex 200, GE Healthcare) in 20 mM Tris-HCl, pH 8.0, 100 mM NaCl, 2 mM 2-mercaptoethanol. Purified protein fractions were concentrated to 500 μl (14 mg/ml) and transferred to an anaerobic glovebox. Protein reconstitution and crystallization experiments were performed under pure argon and all solutions used were degassed and purged with argon before use. Purified enzyme (500 μl, 14 mg/ml) was incubated with 10 molar equivalents of ferrous ammonium sulfate at 22°C for 30 min, followed by incubation with 10 molar equivalents each of NAD^+^ and glycine for 1 h.

### Crystallization and structure solution of T. ammonificans THI4

Initial crystallization conditions were obtained using the hanging-drop method at 22°C, with 2 μl reconstituted enzyme and 2 μl of 100 mM HEPES-NaOH, pH 7.5, 200 mM NaCl and 35% 2-methyl-2,4-pentanediol. Optimization of crystal morphology resulted in cubic shaped crystals in ~2 weeks with 100 mM HEPES-NaOH, pH 7.5, 200 mM NaCl, 35% 2-methyl-2,4-pentanediol, 10 mM praseodymium acetate hydrate as the precipitant.

### Data collection, processing, and structure refinement

Diffraction data were collected on beamline 23-ID-D of LS-CAT at Argonne National Laboratory Advanced Photon Source at a wavelength of 1.033 Å. Data were collected at 100°K and processed using XDS [24] to 2.3 Å resolution in space group I121 (Supplementary Table 2). The structure was solved by molecular replacement using a homology model of *Methanococcus igneus* THI4 (PDB: 4Y4N, ~50% sequence identity). A model of *T. ammonificans* THI4 (four monomers per asymmetric unit) was iteratively built by combining AutoBuild (PHENIX [25]) with manual building in COOT [26]. Structure refinement was performed in PHENIX and REFMAC [25,27]. The parameter file for the bound glycine imine intermediate was generated using phenix.elbow [28]. The coordinates and structure factors are available from the Protein Data Bank (PDB ID: 7RK0).

## RESULTS

### Diversity of non-Cys THI4s and selection of representatives

We first surveyed ~1,000 prokaryotic THI4 proteins in the SEED [18] and UniRef90 [19] databases using BlastP and non-Cys THI4s [15] as query sequences. After removing entries that were truncated, redundant, or from unidentified organisms, there remained 199 unique sequences in which the activesite Cys position was occupied by His (171 cases) or by Met, Leu, Pro, Ala, Ser, Glu, Asp, Tyr, or Trp (28 cases) (Supplementary Table 3). The 199 sequences shared only 47% identity on average.

To analyze this sequence diversity and help select representatives to test for activity, we built sequence similarity networks (SSNs) using the Enzyme Function Initiative webtool [20,29]. The final SSN (E value = 10^-5^, alignment score = 80) contained a series of clusters plus various singletons (Figure 2). Some clusters included non-Cys THI4s that have been tested for activity [7,14,15] but others did not, indicating that much non-Cys THI4 sequence space remained to be sampled.

**Figure 2.**
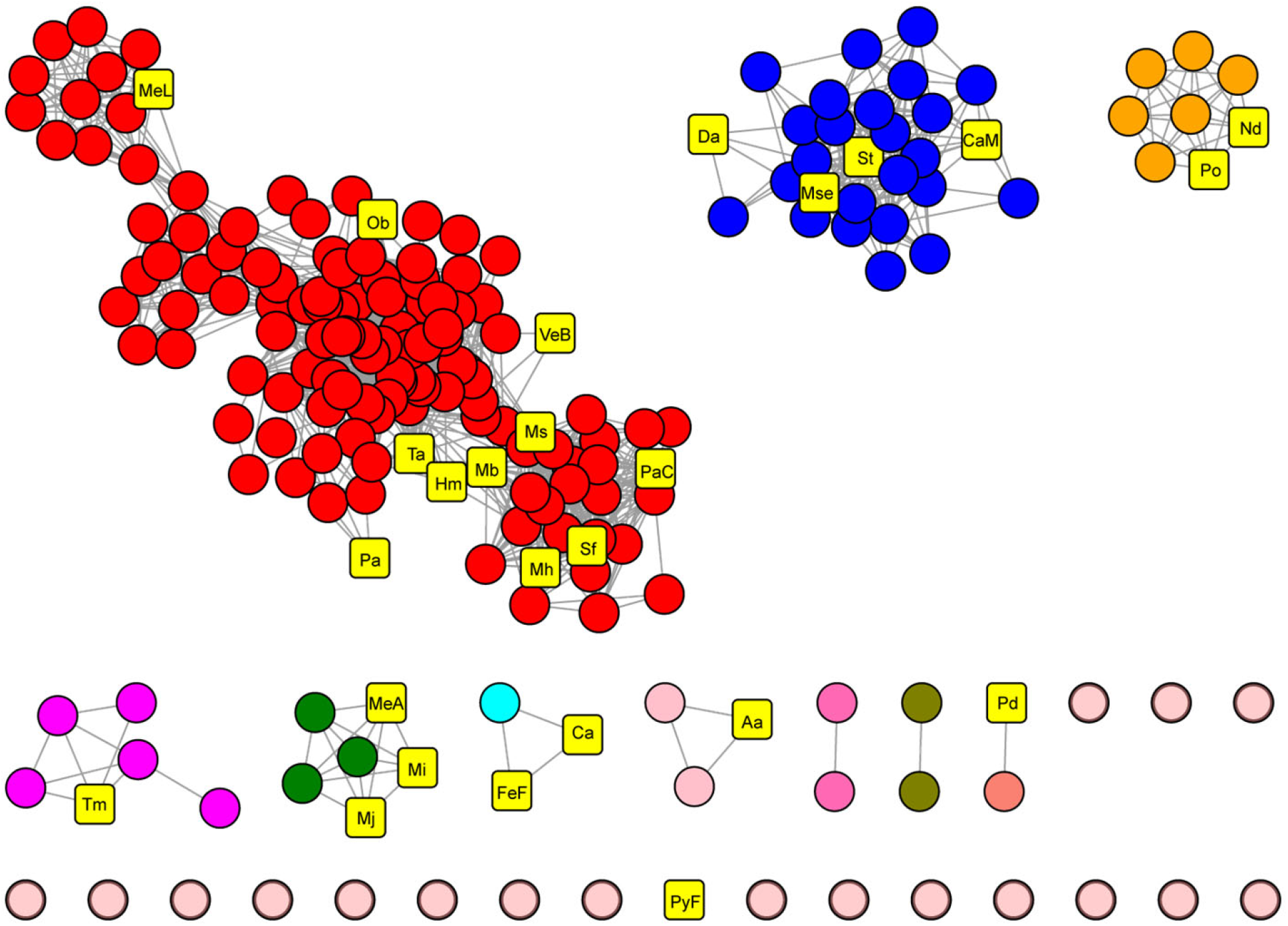
Sequence similarity network (SSN) of 199 diverse non-Cys THI4s. Each node in the SSN corresponds to a single sequence; each edge (gray lines) represents a pairwise connection between two sequences at a BLAST E-value <1 × 10^-5^. Lengths of edges are not significant, except that tightly clustered groups share more similarity than sequences with only a few connections. The 26 representative sequences selected for testing are shown as yellow squares; organism name abbreviations are as in Table 1.

We selected 26 representative sequences from throughout the SSN and from organisms with different ecophysiologies (Figure 2 and Table 1). These sequences were about as diverse (45% average identity) as the whole set of 199 and included THI4s described previously [7,14,15]. Fifteen of the selected sequences were bacterial and 11 were archaeal, eight were from organisms that require or tolerate O_2_, 11 were from mesophiles, 15 were from thermophiles, 22 were from habitats known or likely to be sulfide-rich and four were from habitats likely to be sulfide-poor (Table 1).

**Table 1.**
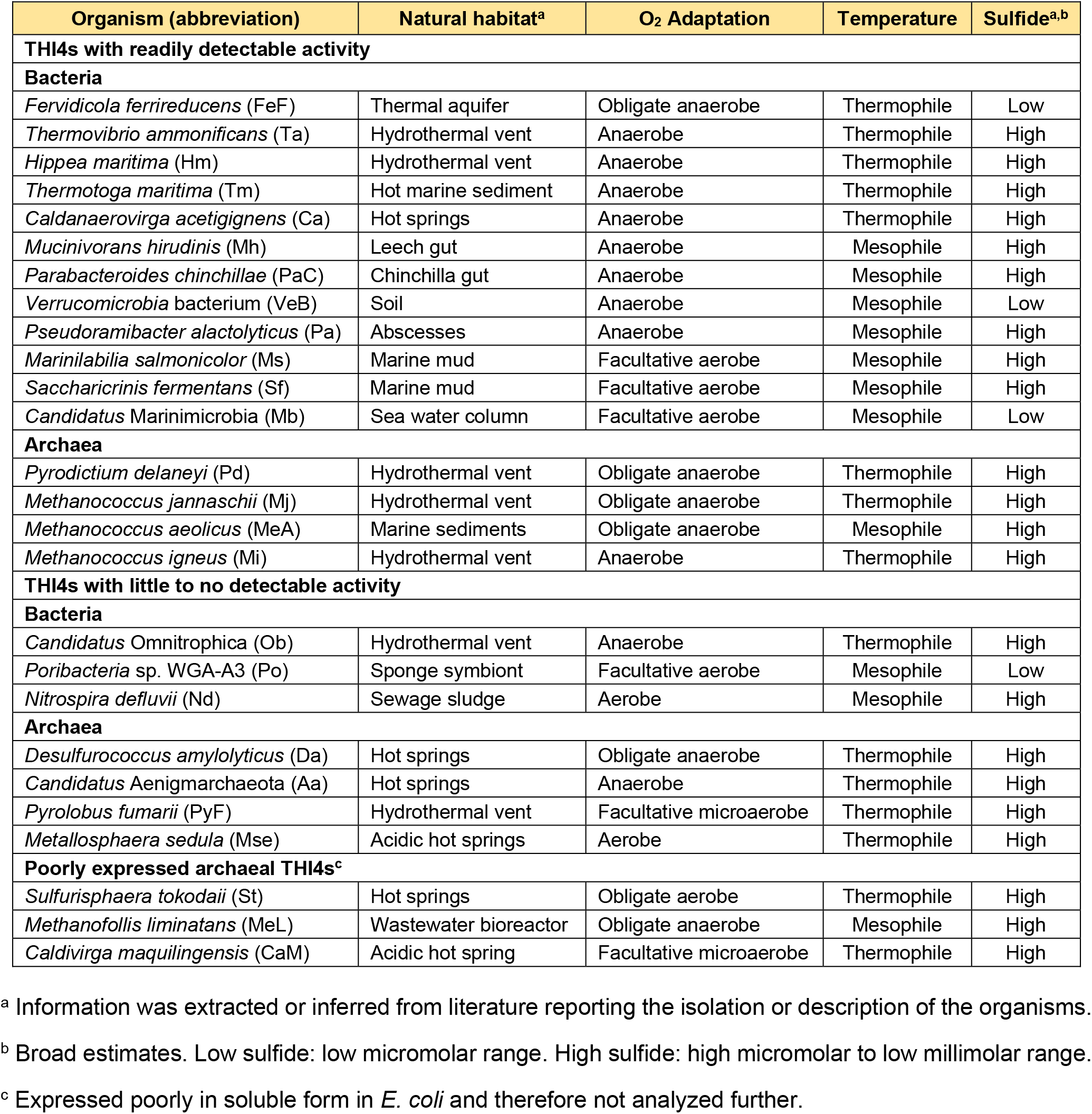
Ecophysiology of the prokaryote sources of the THI4s selected for testing

### Genomic context of non-Cys THI4 genes

The selected THI4s all came from genomes that encode ThiE or ThiN (Supplementary Figure 1), indicating capacity to synthesize thiamin from its thiazole and pyrimidine precursors. All genomes also had ThiD and all but one had ThiC (Supplementary Figure 1), respectively indicating capacity to utilize or produce the pyrimidine precursor. Nearly all the selected THI4s therefore came from organisms with complete thiamin synthesis pathways. Only six genomes encoded the alternative thiazole synthase ThiG, so that THI4 was typically the only identifiable endogenous source of the thiazole precursor. Three of the selected archaeal *THI4* genes abut a gene coding for a protein from the TRASH family, whose members bind nonferrous heavy metals, with characterized examples involved in zinc, copper, cadmium and/or mercury transport/resistance [30–35] (Supplementary Figure 1). Similar clustering occurs in many other archaea and bacteria (Figure 3A). The TRASH proteins in these clusters have two adjacent Cys residues in addition to the four canonical metal-binding Cys residues in the TRASH motif (Cys-Xaa_2_-Cys-Xaa_19–22_-Cys-Xaa_3_-Cys) [30] (Figure 3B); these extra Cys could help ligand a six-coordinate metal such as cobalt, nickel, or iron. Furthermore, non-Cys THI4 and TRASH genes cluster with genes for proteins from families that transport or metabolize cobalt or nickel [36–39] (Figure 3A). Non-Cys THI4s are thus more strongly genomically associated with nonferrous metals than with iron.

**Figure 3.**
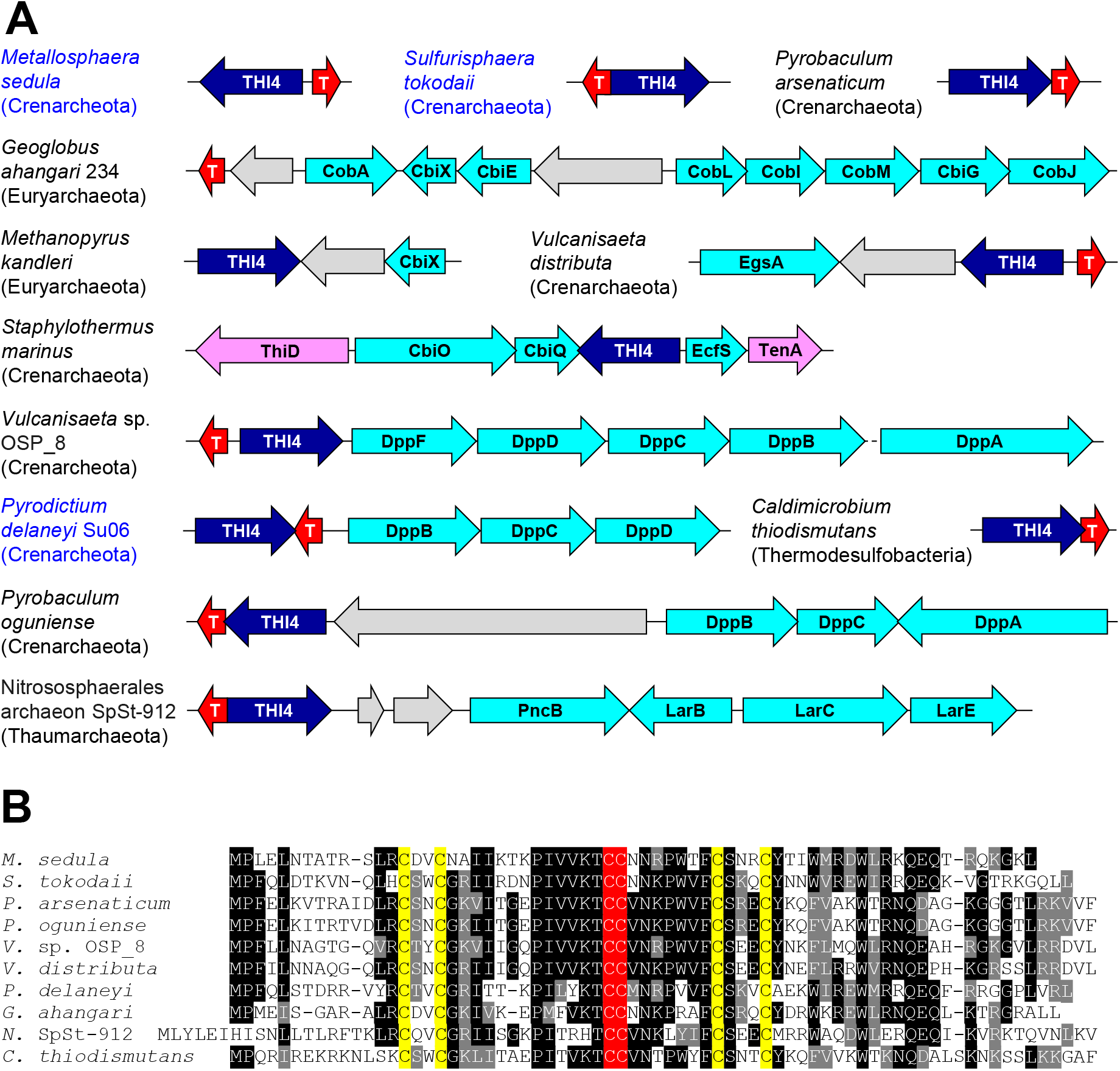
Association of THI4 genes with TRASH genes and cobalt- or nickel-related genes. (**A**) Chromosomal gene clustering arrangements. Names of organisms whose THI4s were tested are in blue. T, TRASH protein gene. Genes colored aqua encode homologs of nickel- and/or cobalt-related proteins: CobA, uroporphyrin-III C-methyltransferase, involved in cobalamin and sirohaem synthesis; CbiX, sirohydrochlorin nickel/cobalt chelatase, which inserts nickel or cobalt into tetrapyrroles; CbiE, involved in cobalamin synthesis; a cobalamin synthesis operon encoding CobL, CobI, CobM, CbiG, CobJ, plus (not shown) CbiC and CbiD; CbiO, CbiQ, and EcfS, the A+A’, T, and S components of a cobalt energy coupling-factor (ECF) transporter. DppA-D,F, subunits of ABC transporters for oligo-peptides, nickel, or (rarely) cobalt; LarBCE, nickel-pincer nucleotide (NPN) cofactor synthesis genes; PncB, synthesis enzyme for the NPN precursor NaAD; EgsA, glycerol-1-phosphate dehydrogenase, which has a nickel or zinc co-factor; gray genes have no known nickel or cobalt associations. Genes colored lilac encode thiamin synthesis or salvage enzymes (**B**) Alignment of the TRASH proteins encoded by genes in part A. The Cys residues of the extended TRASH motif (Cys-Xaa_2_-Cys-Xaa_19–22_-Cys-Xaa_3_-Cys) are in yellow; the two extra Cys residues are in red.

### Functional complementation tests of THI4 activity

The 26 selected THI4s were recoded for *E. coli* and inserted into pBAD24 [23]; the resulting constructs were then introduced into an *E. coli* Δ*thiG* strain [15]. To check THI4 expression, cells were grown in thiamin-supplemented minimal medium and harvested for gel analysis of the soluble and insoluble fractions (Figure 4A and Supplementary Figure 2). Of the 26 THI4s, 23 expressed well, with ≥85% in soluble form (Table 1) and were advanced to testing for thiazole synthase activity by complementation.

**Figure 4.**
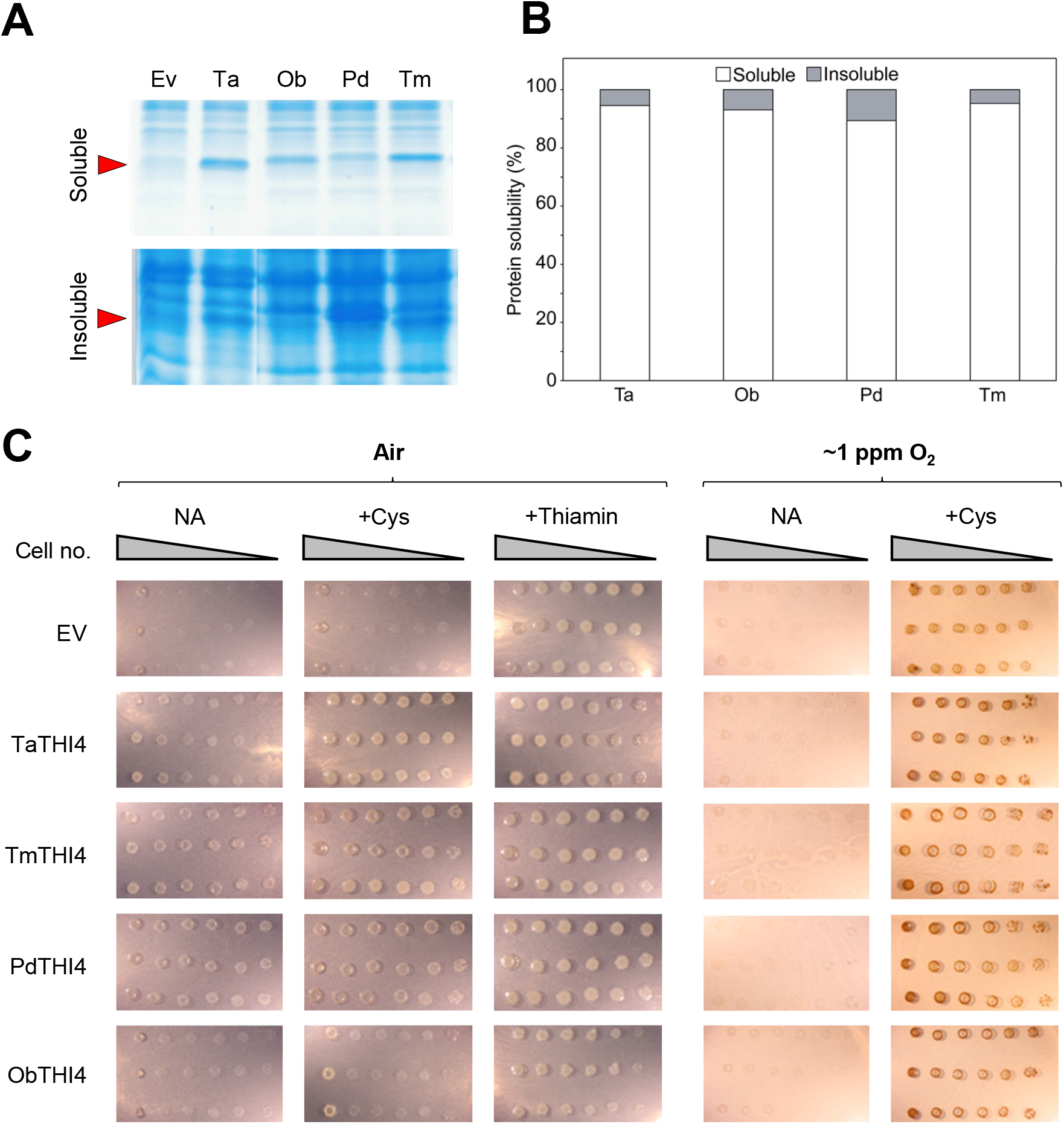
Soluble expression and functional complementation tests of non-Cys THI4s. (**A**) Gel analysis of soluble and insoluble expression in *E. coli* of representative THI4s. Soluble and insoluble fractions of cells were run on 15% gels, stained with Coomassie blue, and scanned to quantify the THI4 band, for which purified *Thermovibrio ammonificans* (Ta) THI4 served as a marker (arrow). Organism abbreviations are as in Table 1. (**B**) Quantification of the solubility (%) of the THI4s from part A. (**C**) Tests of functional complementation of an *E. coli* Δ*thiG* strain by the THI4s from part A or the empty vector (EV). TaTHI4, TmTHI4, and PdTHI4 had clearly detectable activity in air while ObTHI4 did not; none had more than slight activity in ~1 ppm O_2_. Overnight cultures of three independent clones per construct were 10-fold serially diluted and spotted on plates of MOPS minimal medium containing 0.2% (w/v) glycerol and 0.02% (w/v) arabinose with no additions (NA) or plus 1 mM Cys or 100 nM thiamin. The medium used for culture in ~1 ppm O_2_ contained 40 mM nitrate as electron acceptor. Cells were cultured in air or under N_2_ containing ~1 ppm O_2_. Images were captured after incubation at 37°C for 7 d. The high background in the ~1 ppm O_2_ +Cys treatment is due to staining of the inoculum cells.

Complementation tests were run in air or in N_2_ containing ~1 ppm O_2_, plus or minus supplementation with 1 mM Cys to increase intracellular sulfide level [40]. Sixteen strains showed clear growth in air, particularly when supplemented with Cys while seven did not, and no strain showed clear growth in ~1 ppm O_2_ but not in air (Figure 4B and Supplementary Figure 3). The enhanced growth with Cys supplementation fits with use of sulfide as sulfur donor [5,15]. The complementation tests in air thus split the THI4s into one group with readily detectable activity (henceforth: aerotolerant THI4s) and another with little or none. Both groups included THI4s from mesophiles and thermophiles, aerobes and anaerobes, and organisms from high- and low-sulfide habitats (Table 1). There was hence no evident correlation between THI4 aerotolerance and source organism ecophysiology. Neither was aerotolerance correlated with the residue (His, Met, Leu, Tyr, or Ser) that replaces Cys in the active site or with the number of Cys and Met residues, which can affect oxidative instability [41,42] (Supplementary Table 3). To summarize: the complementation data establish that some catalytic THI4s operate quite well (although not optimally, see below) in aerobic conditions but do not suggest why others do not. One of many possible causes is that the redox environment in *E. coli* differs enough from that in the THI4 source, especially for extremophiles (Table 1) [43–45], to disrupt protein disulfide formation or metal insertion and coordination [45–47] and thus prevent THI4 expression in its native form.

It is important to note that supplementation with thiamin markedly stimulated growth of all strains (compare NA and +Thi columns in air in Figure 4B and Supplementary Figure 3). This stimulation shows that the *in vivo* activity of THI4s with complementing activity did not fully meet demand for ADT despite their high expression levels (Figure 4A and Supplementary Figure 2), i.e. that there is room for activity improvement, at least when no Cys is supplied. We revisit this point later.

### Protein structure of the aerotolerant Thermovibrio ammonificans THI4

As structural data for THI4s with little or no complementing activity in air would be hard to interpret in terms of aerotolerance alone since failure has other possible causes, we explored molecular features associated with aerotolerance by determining the crystal structure of the complementation-active THI4 from *T. ammonificans* (TaTHI4) (PDB: 7RK0). This bacterium is an anaerobe but is likely to be intermittently exposed to O_2_ in its habitat near the oxic/anoxic interface in hydrothermal vents, and encodes enzymes to detoxify reactive oxygen species (ROS), including catalase/peroxidase, cytochrome *c* peroxidase, and cytochrome *bd* complex [48,49]. This makes the TaTHI4 structure a potentially informative ‘routinely O_2_-exposed’ comparison with the structures of ‘never O_2_-exposed’ THI4s from the archaeal methanogens *M. igneus, M. jannaschii*, and *Methanothermococcus thermolithotrophicus* [7,50]. Like *T. ammonificans*, these organisms are anaerobic thermophiles from hydrothermal vents [48,51] but, unlike *T. ammonificans*, they inhabit the anoxic region of the vent plume and lack the ROS defense genes present in *T. ammonificans* as well as heme- and manganese-catalase genes (Supplementary Table 4). Furthermore, *M. igneus* and *M. jannaschii* THI4s had similar complementing activity in *E. coli* to TaTHI4 (Supplementary Figure 3), meaning that differences among their recombinant protein structures are unlikely to be artifacts of misfolding in the heterologous host.

Protein crystals of TaTHI4 diffracted to 2.3 Å resolution in space group I121 and the structure was solved by molecular replacement (Supplementary Table 2) using *M. igneus* THI4 as a search model (PDB: 4Y4N) [7]. In common with the methanogen THI4s, TaTHI4 is an overall homooctamer with four monomers per asymmetric unit (Figure 5A). The octamer is tightly packed as a two-layer ring structure with approximate dimensions 73 × 86 Å (height × width) enclosing a ~26 Å diameter pore. The monomer structure (Figure 5B) consists of a long α-helix (residues 6-24) and a barrel-like core domain sandwiched by helix bundles (Figure 5C). The barrel-like core comprises a central five-stranded β-sheet with β6, β2, β1, β12, β13 topology, flanked by a twisted antiparallel β-sheet (β7, β10, β11) connecting α6 and a four-helix bundle (α9, α12, α2, α5). The barrel core is capped by a variation of the Rossmann fold β-hairpin motif. The C-terminal topology is similar to the canonical Rossman fold with an inversion of strands. TaTHI4 is thus structurally homologous to the previously solved methanogen THI4s [7,50].

**Figure 5.**
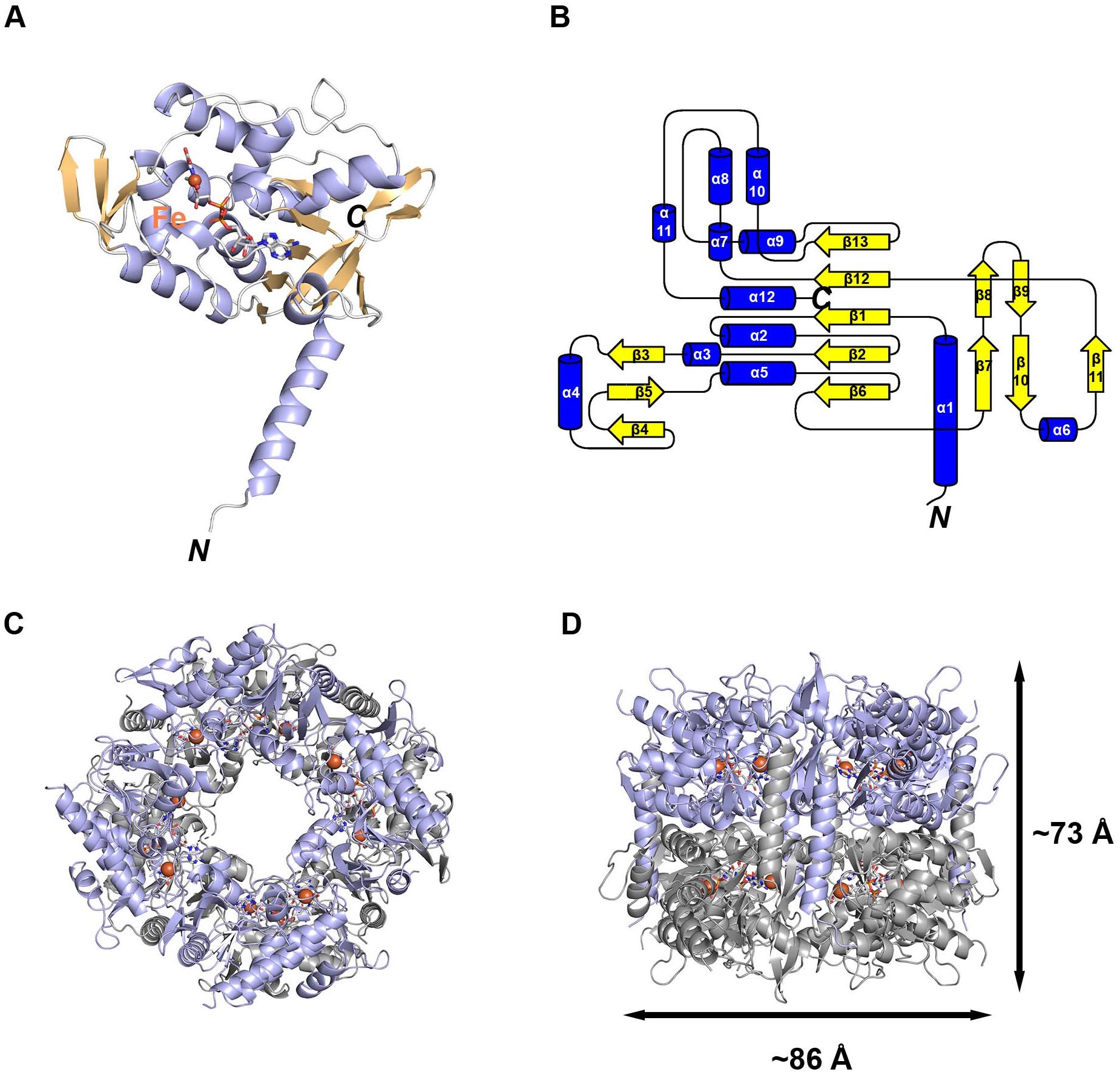
Structure of the aerotolerant non-Cys THI4 from *Thermovibrio ammonificans*. (**A**) Ribbon diagram of the TaTHI4 monomer with bound glycine imine shown as sticks; α-helices are colored blue, β-strands are colored beige; Fe(II) is shown as a brown sphere. (**B**) Topology diagram of TaTHI4 with α-helices in blue and β-strands in yellow. (**C**,**D**) The biologically relevant TaTHI4 homooctamer shown in ribbon representation in alternate top and side views with the metal Fe(II) indicated as a brown sphere.

The enzyme active site lies in a channel formed by two helix bundles and is capped by a loop from an adjoining monomer. Clear density for the TaTHI4 treated with NAD and glycine shows the expected glycine imine intermediate [7] in the active site (Figure 6A). This intermediate adopts an elongated conformation with the adenine moiety positioned in the barrel-like core domain. Binding interactions between the intermediate and TaTHI4 include a hydrogen bond between N6 of adenine and the side chain of Ser178, a bidentate hydrogen bond with the 2’ and 3’ hydroxyls and Glu56, a second hydrogen bond between Arg58 and the 2’ hydroxyl, along with a stabilizing interaction of the glycine imine carboxylate and Arg239 (Figure 6A). Based on our reconstitution conditions, the bound metal in the active site can be assigned as ferrous iron, Fe(II). The iron is coordinated in a pseudooctahedral geometry with three sites occupied by the pincer-type ligand of the bound glycine imine, axial ligands of His175 and Asp160, and a predicted water molecule (Figure 6B,C).

**Figure 6.**
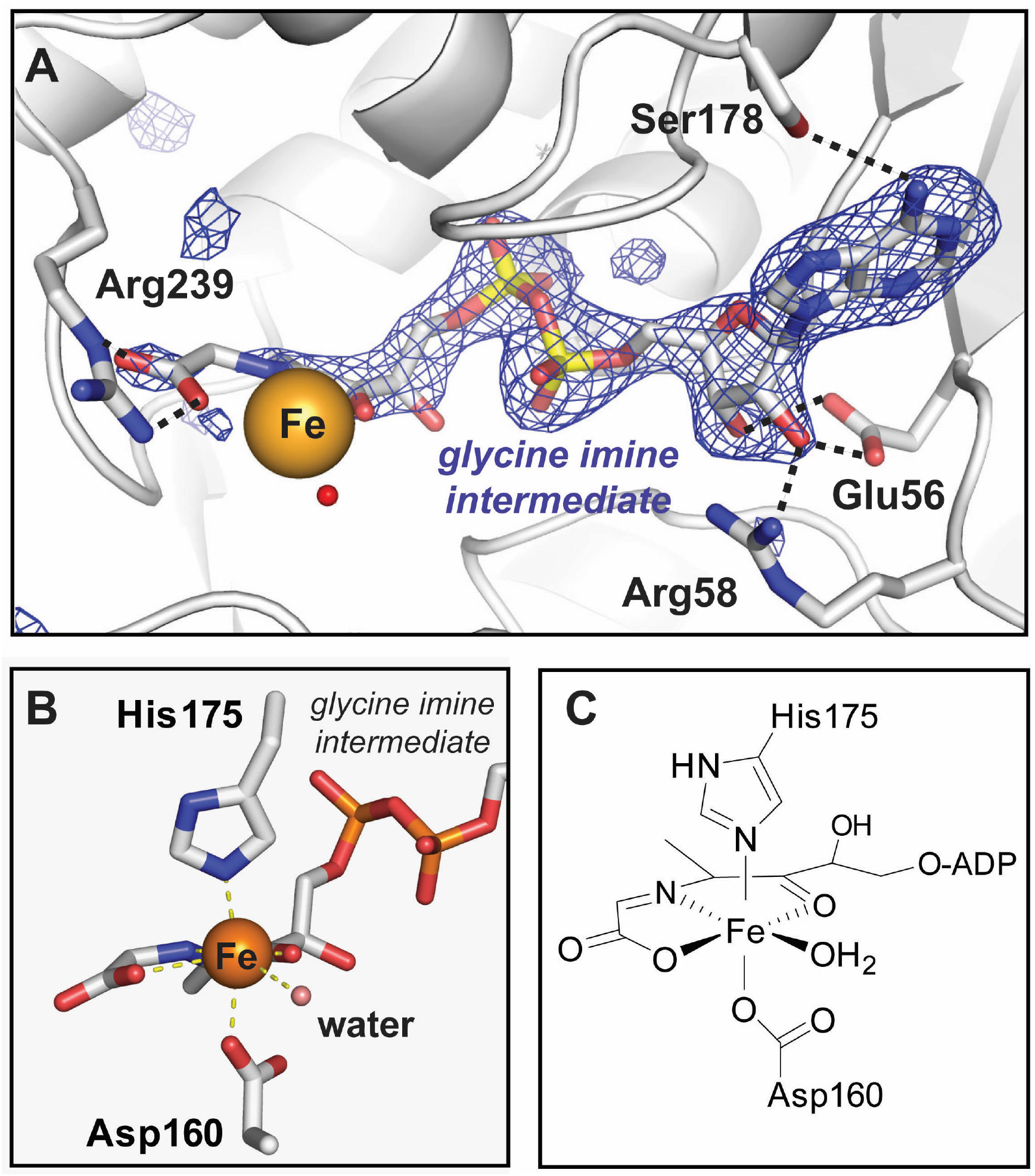
The active site of *Thermovibrio ammonificans* THI4. (**A**) Interactions of the bound glycine intermediate with the TaThi4 protein active site. A glycine imine intermediate omit map, Fo-Fc contoured at 3.5σ, is shown in blue. (**B**) Ligand geometry around the metal center with a near-octahedral coordination environment as diagrammed in (**C**).

To probe the role of the Met158 residue that in TaTHI4 occupies the position of the sacrificial Cys, we changed Met158 to His, Lys, or Cys and assayed complementing activity (Supplementary Figure 4). The mutant enzymes performed similarly to wild-type TaTHI4 in both air and ~1 ppm O_2_ atmospheres, with or without Cys supplementation. This result confirms the inference from natural variation in this residue (Supplementary Table 3) that it has no role in catalysis. That the Cys form did not have greater activity in the absence of Cys supplementation suggests that it cannot operate efficiently in suicide mode and still depends on sulfide as sulfur source, as was the case for *M. jannaschii* THI4 [7].

### Investigation of TaTHI4 metal preference in vivo in E. coli and yeast

As we could not obtain quantifiable *in vitro* activity [14] from reconstituted TaTHI4, we sought to test metal cofactor preference *in vivo*. Adding 100 μM cobalt, nickel, zinc, or manganese to the medium of *E. coli* expressing TaTHI4 (or five other THI4s) did not detectably affect complementing activity in air (Supplementary Figure 5). This result does not indicate which metal is preferred, but does show that the supply of this metal is not limiting in *E. coli* grown with normal trace metal supplementation.

We also tested metal preference using yeast, which resembles *E. coli* in having native cobalt-dependent enzymes and cobalt uptake systems [52] but differs in having no native nickel enzymes or high-affinity nickel uptake system and in needing an added nickel transporter to express a foreign nickel enzyme in active form [53]. We therefore tested TaTHI4 for ability to complement a yeast Δ*THI4* strain. The observed complementation (Supplementary Figure 6) proves that TaThi4 has access to its metal cofactor and implies that this metal is more likely to be cobalt than nickel if it is not iron.

## DISCUSSION

The biochemically characterized catalytic THI4s come from archaea of anoxic, sulfide-rich, reducing environments, use an O_2_-sensitive sulfur donor (sulfide) and cofactor (ferrous iron), and are thermophilic [7,14]. It was consequently *a priori* doubtful whether catalytic prokaryotic THI4s could function in mild, fully aerobic conditions, and recent evidence [5] that putatively catalytic plant THI4s are expressed only at severely hypoxic stages of seed development reinforced this doubt.

However, our survey of THI4 sequences and their provenances began to dispel the doubt because non-Cys THI4s confidently predicted from genomic context to be functional thiazole synthases were found in aerobic or aerotolerant organisms from mild environments. Confirmation that certain non-Cys THI4s – the majority of those tested, in fact – can function in air came from complementation experiments, which also indicated that these THI4s use sulfide, or a sulfide metabolite, as sulfur donor.

Having established that certain non-Cys THI4s can – unexpectedly – operate in mild, fully aerobic conditions, we investigated characteristics that enable them to do so. We could dismiss any role, positive or negative, for thermophily because complementing activity varied similarly in thermophiles and mesophiles (Table 1). There was likewise no association between complementing activity and the O_2_ adaptation of the source organism (Table 1). Nor did the residue that replaces the sacrificial Cys or the number of Cys or Met residues appear to be important (Supplementary Table 3).

What then, might confer aerotolerance? Comparative genomic analysis provided a clue by associating non-Cys THI4s with proteins that bind, transport, or metabolize nonferrous transition metals, notably cobalt or nickel (Figure 3A), and the observed complementing activity of TaTHI4 in yeast favored the possibility that the metal cofactor is cobalt (Supplementary Figure 6). Neither the comparative genomics nor the yeast complementation data ruled out a ferrous iron cofactor, however.

Comparing the TaTHI4 structure – the first deposited for a bacterial THI4 – with those of the ecologically less O_2_-exposed THI4s from archaeal methanogens (PDBs: 4Y4M, 4Y4N, 6HK1) [7,50] did not provide strong evidence on the aerotolerance phenomenon. The active sites are largely identical, with one major difference, the variation of residues (Met *vs*. His) replacing the Cys residue of suicide THI4s. Variants at this position, however, do not correlate with aerotolerance (Supplementary Table 3). The octamer structures show similar dimensions of the overall barrel-like architecture and similar surface electrostatic charge. Notable differences are two extended loop regions in TaTHI4 at the entrance to the large active-site pore (residues 134-142 and 186-197); however, based on the pore diameter, these changes are unlikely to affect O_2_ diffusion. An analogous comparision of the structures of O_2_-sensitive and O_2_-tolerant homologs of [NiFe] hydrogenase showed distinct, well-defined tunnels to the metal center [54,55]. In this case, structural variations of the tunnel were predicted to regulate O_2_ diffusion to the reactive metal center. All THI4s have a large pore (~26 Å) linking the active site to solvent, precluding variation in O_2_ diffusion. The lack of dissimilar structural features favors the alternative possibility of a role for the active site metal in O_2_ tolerance.

Semi-quantitative *in vitro* assay of *M. jannaschii* THI4 activity [14] showed that cobalt supported ~60%, and nickel ~25%, of the activity given by iron. The presented TaTHI4 structure contains an obligate bound Fe(II) atom, based on Fe(II) being the only metal present during protein reconstitution. The ligand geometry in the structure (Figure 6) does not preclude other metals, including Co(II) or Ni(II). A bound Ni(II) would be predicted to favor a square pyrimidal geometry [56] over the octahedral although the active site could possibly accommodate a minor shift in ligand position to square pyrimidal. There is solid precedent for octahedral Co(II), including colbalamin enzymes and methylmalonyl-CoA carboxytransferase [57]. In terms of THI4 chemistry, Co(II) is less likely to react deleteriously with molecular O_2_ than Ni(II) and Fe(II). Contrasting with many examples in which mononuclear Fe(II) reacts with O_2_ to generate reactive O-atom transfer species in metalloenzymes and model compounds [58,59], reaction of Co(II) with O_2_ has strong precedent for reversible formation of Co(III)-superoxide species in the absence of additional reducing equivalents or protonation [60], implying that replacing iron with cobalt could provide a mechanism for the observed O_2_ tolerance. In addition, a Ni(II)-SH intermediate in the catalytic cycle would be particularly stable [61], disfavoring catalysis. Other metals are known to be able to perform THI4 chemistry, notably Zn(II), but the ligand geometry and metal preference analysis of *M. jannaschii* THI4 [14] argue against a tetrahedral liganded metal.

In summary, the comparative genomics, functional complementation, and structural evidence collectively implicate the bound metal as a natural determinant of THI4 aerotolerance, with Co(II) the best candidate although Ni(II) cannot be rigorously excluded. Definitive evidence on this point will require development of quantitative *in vitro* assays for THI4 activity. Besides the nature of the metal inserted, there may well be other determinants of aerotolerance. We hope to identify such determinants, and to gain insight into the metal cofactor, from ongoing continuous directed evolution experiments [62] to improve the complementing activity of native non-Cys THI4s. If – as seems likely [63] – such improvement is possible, there is a realistic prospect of replacing suicidal plant THI4s with catalytic THI4s that work well in aerobic conditions and thus slash the energy cost of thiamin synthesis.

## Abbreviations

ADT: adenylated carboxythiazole
IPTG: Isopropyl β-D-1-thiogalactopyranoside
ROS: reactive oxygen species
SSN: sequence similarity network

## Data Availability

Coordinates and structure factors of the *T. ammonificans* THI4 crystal structure have been deposited in PDB under code 7RK0.

## Acknowledgments

We thank Prof. Leslie J. Murray for helpful discussions.

## Author Contribution

A.D.H., J.J., and S.D.B. designed the research; J.J., J.-D.G.-G., and B.J.L. performed assays; J.J. and A.D.H. ran bioinformatic analyses; Q.L., Y.H., and S.D.B. carried out structural analysis; A.D.H. wrote the article with input from J.J. and S.D.B.

