## Supplementary Table 2 for "Structure and function of aerotolerant, multiple-turnover THI4 thiazole synthases"

**Supplementary Table 2  *T. ammonificans* THI4 data collection and refinement statistics**

| | TaThi4 |
|---|---|
| **Data collection** | |
| Space group | I121 |
| Cell dimensions | |
| *a*, *b*, *c* (Å) | 89.89, 89.69, 131.81 |
| α, β, γ (°) | 90, 96.98, 90 |
| Resolution (Å) | 43.63-2.30 (2.36-2.30)* |
| $R_{merge}$ | 0.092 (1.552) |
| $I / \sigma I$ | 8.23 (0.77) |
| Completeness (%) | 99.46 (96.62) |
| Redundancy | 2.0 (2.0) |
| **Refinement** | |
| Resolution (Å) | 43.63-2.30 |
| No. reflections | 93685 |
| $R_{work}$ / $R_{free}$ | 0.217/0.275 |
| No. atoms | |
| Protein | 7835 |
| Ligand/ion | 160 |
| Water | 58 |
| *B*-factors | |
| Protein | 50.69 |
| Ligand/ion | 41.32 |
| Water | 47.34 |
| R.m.s. deviations | |
| Bond lengths (Å) | 0.009 |
| Bond angles (°) | 1.12 |

*Values in parentheses are for highest-resolution shell.
