## Supplementary Table 3 for "Structure and function of aerotolerant, multiple-turnover THI4 thiazole synthases"

**Supplementary Table 3 Sequences of 199 non-Cys THI4s from SEED and UniRef90 databases**

The 26 sequences selected for testing are boxed. Those that had little to no complementing activity are in red font. Those that were poorly expressed in soluble form in *E. coli* and were not tested are in gray font. The residue that replaces Cys in the active site is highlighted in cyan. Met and non-active-site Cys residues in each sequence are highlighted in yellow; the number per sequence is given in the header. Mean values were: Active THI4s: 10.6 Met, 2.3 Cys. Inactive THI4s: 9.1 Met, 3.1 Cys.

```
>Caldanaerovirga acetigignens (2 Cys, 11 Met)
MKSFSIPVPDTKVSSLIMKHYFKDLEDAVKSDVIVAGAGPSGLTCAWTLADQGYKVTVLDRRLAPGGGIWGGAMSFNKVVLQKDVEWILKEADVPFV
EDEGALVVSAPLFASKLIAKAAAHPGIRFFNMLTVVDLHSSGDRITGVVVNNSAIEMAGLHVDPMVLTAKAVLDATGHDAVLANLYSRRAGTGLIRE
SFMNAEKGEEDVVANTRMLAPGLFVAGMAANNVEGGCRMGPIFGGMLLSGKKAARLIINYLSENSK

>Candidatus Marinimicrobia bacterium (4 Cys, 12 Met)
MEKIVSFGIIDSYQKKLKENLEVDVAIVGGGPSGLIAAKYLAQAGKKVVLFERKLAPGGGMWGGAMMFNQIVVQEDAISILEDVDISYNLYEEGYYV
CDSVEATAALIFSAKKAGATIFNCFSVEDVVFQRGSVAGVVVNWASVHREGMYVDPLVIMAKAVLDSTGHSCEVASILARKNEVKLMTRTGNIMGER
SLSIEEGELTTIENTKEIFPGLYVSGMAANAVSGSFRMGPIFGGMLMSGKKVAGLINDRLSKRDGCK

>Fervidicola ferrireducens (2 Cys, 11 Met)
MKSFSVPVPDTKVSSLIMKHYFKDLEDAVKSDVIVAGAGPSGLTCAWTLADQGYKVTVLDRRLAPGGGIWGGAMSFNKVVLQKDVEWILKEADVPFV
EDEGALVVSAPLFASKLIAKAAAHPGIRFFNMLTVVDLHSTGDKITGVVVNNSAIEMAGLHVDPMVLTAKAVLDATGHDAVLANLYSRRAGTGLIRE
SFMNAEKGEEDVVANTRMLAPGLFVAGMAANNVEGGCRMGPIFGGMLLSGKKAARLIINYLSENSK

>Hippea maritima strain ATCC 700847 (2 Cys, 9 Met)
MNNLDERVISRAIVERYMNKLLDYLECDVTIVGGGPAGLVCAYYLAKANIKVAIFDKRLTIGGGMWGGAMLFNEIVVQEIGREILDEFGINYEKYTD
GYYTADSIEATTTLISKTVKAGAKIFNAIEVEDVVFKKIDGQYRVNGLVVGWTTVNMAGLLVDPLVVTSKYVIDATGHDADIANILTRKGGIKLNTP
EGVVIGEKPMWAEVGEQSTIEETQEVYPGLIVAGMAAVAVSGSHRMGPVFGGMLNSGKKAAQIVIESLKK

>Marinilabilia salmonicolor (2 Cys, 12 Met)
MEQIVSSGIIDSYFSKLKENLAVDVAIVGGGPSGLIAAYYLAKTGKKVALFERKLAPGGGMWGGAMMFNEIMVQKEALHILKELGIEYKHYRDDYYT
VDSVHATSALTYHATKAGARIFNCTSIEDVVFHNNIVSGLVINWAPVHREGMHVDPLIIMAKAVIDGTGHDCEIVHTVARKNDIKIDTPSGKVMGER
SLAVEEAERTTVDNTKEVFPGLFVSGMAANGTSGSYRMGPIFGGMLLSGQKVAGIISEKLAKEAIMESANN

>Methanocaldococcus jannaschii DSM 2661 (= Methanococcus jannaschii) (3 Cys, 10 Met)
MVNLMNIKDIKLNADETKTTKAILKASFDMWLDIVEADVVIVGAGPSGLTCARYLAKEGFKVVVLERHLAFGGGTWGGGMGFPYIVVEEPADELLRE
VGIKLIDMGDGYYVADSVEVPAKLAVAAMDAGAKILTGIVVEDLILREDGVAGVVINSYAIERAGLHIDPLTIRSKVVVDATGHEASIVNILVKKNK
LEADVPGEKSMWAEKGENALLRNTREVYPNLFVCGMAANASHGGYRMGAIFGGMYLSGKLCAELITEKLKNKE

>Methanotorris igneus (= Methanococcus igneus) (2 Cys, 8 Met)
MDVRLRADEYATTRAILKSAFDMWLDIIDVDVAIVGGGPSGLTAARYIAKEGYKVVVLERHLAFGGGTWGGGMGFPYIVVEEPADEILREVGVKLEK
VEGEDGLYTADSVEVPAKLAVGAIDAGAKVLTGIVVEDLVLRENRVAGVVINSYAIEKAGLHIDPITITAKYVVDATGHDASVVTTLSRKNPELGLE
VPGEKSMWAEKGENALLRNTREVYPGLFVCGMAANAVYAGHRMGAIFGGMYISGKKCAEMIVEKLKNNE

>Methanococcus aeolicus strain ATCC BAA-1280 (3 Cys, 8 Met)
MDISKIDLKADEKAVTKSIFKATYEMWMDNLEVDVVIVGGGPSGLTAGRYLADAGVKVLILERHLSFGGGTWGGGMGCPYITVQSPADEILSEVGIK
LEEGEDGLFVADSVEVPAKLGTGAIDAGAKVLTGIVVEDVILKEGKVSGVVINSYAINKAGLHIDPLTINAKYVIDATGHDASVACTLARKNEDLGL
VIPGEKSLWADEGENGLLKYTKELFPGLFVCGMASNATHGGYRMGAVFGGMYISGKIVADMILEKLKNE

>Mucinivorans hirudinis (3 Cys, 10 Met)
MEKIVSAGIVESYFDKLRRNLVLDVAIVGGGPSGLVAAYYLAKAGRRVALFERKLAPGGGMWGGAMMFNDIVVQSDALPILEELGVSYRHYRGDAYL
VDSVHATAALIYAATRAGATIFNCYSVEDVVFKDERVAGLVVNWAPVIREGMHVDPLVIMATAVLEGTGHDCAIARLVARKNGVRLNTPTGEVIGER
SLSIEEAERTTVENTKEIYPGLFVSGMAANGVSGSFRMGPIFGGMLLSGKKAAQMICDSL

>Parabacteroides chinchillae (3 Cys, 11 Met)
MEQIVSTGIIDSYFAKLKSNLSVDVAIVGGGPSGIVAAYYLAKAGKKVALFDRKLAPGGGMWGGAMMFNDIVVQEEAMPIVKELGVSYHAAGNGTYI
MDSVHTTSALIYQATKAGATIFNCYSVEDVVFHNDAVAGVVVNWAPVIREGMHVDPLTIMAKAVLEGTGHDCEVARTVARKNDIKLNTPTGGVIGER
SLNVELGESTTVENTKEIYPGLFVSGMAANGVSGSFRMGPIFGGMLMSGKKAAELICDKLGK

>Pseudoramibacter alactolyticus ATCC 23263 (2 Cys, 19 Met)
MLSDTKISEAILTTYTDRFKQMLSSDAVIVGGGPSGLIAAYYLGKAGVKTTLLDRRLSVGGGMWGGGMMMNQIVVQKSVLPILEEMGIACKAYDAEH
YTVSSVACISGLIFRAAQSGATIMNLVTMEDAVVREGRLEGLVINWSTVEMAHLMVDPLMMDARVVLDATGHDAALVTKLVERMGPVLNTPSGGLEG
EKPMWADHGEKQVVANTREVYPGLYVSGMAANATFGGQRMGPVFGGMLLSGKKAAEAMLRRLAQ

>Pyrodictium delaneyi (0 Cys, 10 Met)
MGIASFYPGELEKQYSEAKLARIALKVALEKLSYAVEADVAIAGAGPAGLTLAWLLAEQGLRVTLVEHRLSTGGGMKGGSMLFPVALVEEGLAAVIL
EKAGVRLHRVGEGLYAMDPVEAVAKLTARAVDAGAVILPGLHVEDLIVRGSGSNVRVAGIVVNWAPVVEAGWHVDPLYIEARAVVDATGHDAQLARL
LERRLPGSLKVPGMSSLDVWTGERQVVEHTGEIFPGLYAAGMSVAEVYNLRRMGPVFGGMIASAARLAEMLAERLAGKRMGLATGVARSG
```

>Saccharicrinis fermentans DSM 9555 (2 Cys, 10 Met)
MEQIVSVGIVDSYFKKLKENLTVDVAIVGGGPSGMVAAYYLARQGFKVSVYERKLAPGGGMWGGAMMFNEIVIQKEALPILDELNISYKHYDKDYYT
LDSVHATSALIYHATQAGATFFNCTSVEDVVFLDNKVSGVVLNWAPVHREKMHVDPLVIMAKAVIDGTGHDCDIARILERKNNIQLLTASGKVEGER
SLSIDEAERTTIENTKEIYPGLYVSGMASNGVSGGFRMGPIFGGMLLSGKKVANLIADNLNK

>Thermotoga maritima strain ATCC 43589 (2 Cys, 8 Met)
MRDVLISRLIVERYFEKLRNSLELDVAIVGAGPSGLTAAYELAKNGFRVAVFEERNTPGGGIWGGGMMFNEIVLEKELENFLKEVEIEYEVKEDHIV
VDSVHFASGLLYRATKAGAIVFNNVSVEDVAVQNGRVCGVVVNWGPTVRLGLHVDPITVKASFVVDGTGHPANVVSLLAKRGLVEMKTEFPMDADEA
EKFVVDNTGEIFPGLLVSGMAVCAVHGGPRMGPIFGGMILSGQKVARIVSERLR

>Thermovibrio ammonificans strain DSM 15698 (3 Cys, 10 Met)
MQNLSEVVISEAIITAFMEKLKSHLETDVAIVGGGPSGLVAGYYLAKKGYRVAIFERRLSIGGGMWAGAMFFNEIVVQEMGREILDEFGVNYREFKP
GYYLADAVEATTTIASKAVKAGATVFNGVTAEDVVLKQVNGQYRVCGLVINWTTVELNHLMVDPLVITAKYVVDATGHDASVVSTLQRKAGIKLNTE
TGCVVGEKPLWASVGEEDTVKNSKEVFPGIYVSGMAANATCGSHRMGPVFGGMLMSGKKVAEEIAAKLNQNKEA

>Verrucomicrobia bacterium (1 Cys, 11 Met)
MLNEVTISRAIIDAYFKKLTRHLEVDVAIVGGGPSGLVAGHDLARAGKKVALFESKLAIGGGIWGGGMGFNEIVVQEAAREMLVEFGLRATEFEPGY
YTLDAVHVAAALAARAMEAGLTVFNLTSMEDVVIQKDRVAGLVLNWTAIRHLKWHVDPLTIHSRFVLDATGHPASVAETLVRKMNVRLDTTTGGLVG
EKPMAAEDGERQTVENTREVYPGLFVSGMAAITVCGGHRMGPVFGGMLLSGRKAAQAMLAELGT

>Candidatus Omnitrophica bacterium 4484_171 (8 Cys, 9 Met)
MLEETIISKAIIDSYHNKLSSIIDVDAAICGGGPSGLVCAASLAAAGKKVVLFEKKLSLGGGMWGGGMMFNEIVVQKKAKKILDEFSVRTKKYKENY
YLADSSECVCALGYNAVHSGAVIINGVFAEDVMVKKNRICGLVINWSAAASANLHVDPLTVRAKFVVDATGHPSEVVKVVEKKSGVKIKTKTGKVLG
EKSMWAHAAENTIEKNTRQIAPGLFVTGMCANAVCGAPRMGPIFGGMLLSGKKCAKIILSRL

>Poribacteria sp. WGA-A3 (5 Cys, 10 Met)
MDNLQPAPLRERDVTRHIAREFYKEFDQLIESDVIIVGGGPSGLVCAHDLATQGFRTLLIEQSLALGGGFWSGGYLMNKATLCEPAHSILENMGVPC
KPVKDCAGMRIVDPPHATARLIASAYEAGAKVLNLTRVVDLILHGEGVLEGVVVNNTTAEMAGHDMIHVDPIALESRVVVDATGHDAVVVGLLNQRG
LYATVPGNGAMWVARSEAMVVDNTREVFPNCFVTGLAVAAVDGSPRMGPAFGSMLLSGRRAADLVRHKLKGE

>Nitrospira defluvii (6 Cys, 12 Met)
MEELARSRKACSTAVEGEYRMGKPKPAPLRERDITRQIAREYYKEFDQLIESDVIIVGAGPSGLICAHDLGRMGIKTLIVEQSLALGGGFWSGGYLM
NKATICAPAHKILKEVGVPCKQIKECPGMYMVDPPHATGALIAAAYNAGAKIMNLTRVVDLILRREGVLEGVVVNSTTAEMAGHDIIHVDPIALESK
IVVDATGHDAVVVNLLHKRGLYQQVPGNGAMWVSRSEEEVMDRTGEVSPNCFVIGLAVAAVFGTPRMGPAFGSMLLSGRYGAELIRDKLKNR

>Desulfurococcus amylolyticus strain DSM 18924 (1 Cys, 5 Met)
MSLESHITRVIWEEASRDWVELSSCDIVVVGAGPSGLTAAKYLAEKGLKTLVLERRLSFGGGIGGGGMLLHKTVVDERGLGILRDFNIRYKPSSIKG
LYVVDTAELTAKLAAGALDAGAKIIPGISVEDVIVRYNPFRVQGVVVEWSAVQLSGLHVDPLFIESKAVIDATGHDAEVLRILEKKNPESKVKIPGE
KSAYSEKADVDVVEYTGRVIPGLYATGMAVAAVRGLNRMGPIFTGMLLSGRKVAEAVIRDLESAPK

>Candidatus Aenigmarchaeota archaeon (2 Cys, 10 Met)
MGEIFSKVSEKEVTSAIVSGFIKEFEKIIESDVIIVGGGPSGLMAGKELSSKGKKVVIIERNNYLGGGFWTGGYLMNKITVRHPGEEILKDLGIPFE
EFGEGLYLADGPHACSKLIAATCDAGVKILNMTTLEDVVLKEKGAVGGVVINWTPIETLPREIASVDPIALESKVVIDATGHDANVVKKIEERGILK
TKGYGAMWVEKSEDMVVKYTGEVHPGLVVTGMAVSTFFGLPRMGPTFGAMLLSGKKAAEVTMEILSR

>Pyrolobus fumarii strain DSM 11204 (0 Cys, 10 Met)
MVIPGHMTTRRTAMPGLDAIITRVIIEEASKELVEYAESDVIVVGAGPAGLTAAFYLAKRGFRVLVLERRLSVGGGIGGGGMLFHKVLVQEEALPVL
NDMGIRVHPTSVKGIYSLDSVALITGLASAAVNAGAKIILGLEAVDLVVRKEGERHRVAGVMALWSAVGIANLHVDPLMFEAKAVVDATGHDARLAR
IAHQKLRGEAPEVPGDGPAWAEEGEKLVVKATGELIPGLYVAGMAATAVKGYYRMGPIFGGMLLSGKKVADLITEKLRGK

>Metallosphaera sedula strain ATCC 51363 (0 Cys, 8 Met)
MNIKQVDEIKITRYILKATFEDWMDFSVNDVVIVGAGPSGLAAAYYSAKAGLKTTVFERRLSFGGGIGGGAMLFHKIVIESPADEILREIGVKLQKF
EEGVYVVDSSEFMAKLAAATIDAGAKIIHGVTVDDVIFRENPLRVTGVAVEWTATQMASLHVDPLFISAKAVVDATGHDAEVISVASRKIPELGIVI
PGEKSAYSEIAEQLTVEQSGEVAPGLYAAGMAVTEIKAIPRMGPIFGAMLLSGKKVAEDIIKNLQANSATLKSVQKE

>Caldivirga maquilingensis strain ATCC 700844 (0 Cys, 10 Met)
MAGISIREASITRAIVNSALKLLSEYSSVDVAIVGAGPSGMTAAYYLAKAGLKTLVLERRFSFGGGIGGAASHLPSIIVEHPVSEILSKDFGIKIMD
MGDGLFTVDPAEMIAKLAVKAIDAGAKFLLGVHVDDVIYRDNPPRITGLALYWATIQMAGVHTDPFFIESNAVVDATGHDAEVAAVASRKIPELGIV
VRGEKSAYVGVAEDLVVKYTGKVIDGLYVTGMAVAAVHGLPRMGPIFGSMIMSGKRVAEIIIEDLKGNH

>Methanofollis liminatans DSM 4140 (5 Cys, 12 Met)
MELDEVTISRAILATQMETMVEYLDLDVAVVGGGPSGITCAALLAEKGVKVGLFEKKLSIGGGMWGGGMMFPRIVVQAEAKRILDRFGIASKEFEPG
YHVAKSVEAVSKLTAAACTAGAEFFNLIAVEDVVIKGDGRLAGLVVNWSPVEMAGLHIDPLTIRCKAVVDASGHDATIAHMVAKKGGDLPIRGEGFM
WADRAEGNILEHTREVFPGLFVCGMAANAVAGECRMGPIFGGMLLSGERAADLAAAAVLHP

>Sulfurisphaera tokodaii strain DSM 16993 (0 Cys, 10 Met)
MDSNSIKVKQVDEVKISKYILKYTFQDWEDIVESDVVIVGAGPSGMTAAYYLAKAGLKTVVFERRLSFGGGIGGGAMLFHKIVIESPADEILKEMKI
KLNKVEEGVYIVDSAEFMAKLAASAIDAGAKIIHGVTVDDVIFRENPLKVVGVAVEWTATQMAGLHVDPLFISAKAVVDATGHDAEVISVAARKIPE
LNIVIPGEKSAYSETAEELTVENTGMVAPGLYAAGMAVTEVKGLPRMGPIFGAMVLSGKRVAEIIIKDLRYS

>Deltaproteobacteria bacterium HGW-Deltaproteobacteria-1
MVLDEIVISKAIIERFLEKLLQATDVDVAIVGGGPSGLVAAYYLASAGKKVALFERKLSLGGGMWGGGMMFNEIVVQDEAREILDVFDIRYREYQQG
YYTADAVLAVTSICSQAARAGASIFNCVSVEDVMIREGRVTGLVINWSPVEMAGLHVDPLTIAAGSVIDTTGHATEVLKVIERKADMQLATPSGKLV
GERSMWAEKAERLTMDNTRQICPGVYVAGMSANAAFGGPRMGPIFGGMLLSGRKVAELILASS

>Thermogladius calderae (strain DSM 22663 / VKM B-2946 / 1633)
MELESIITRLVVEESARELVELSESDVLVVGAGPSGLTAAKYLADKHLKVVVLEKRLSYGGGIGGGGSLFHKVVVDERALPVLGDFKVRYKAAGVAG
YYVVDSAELMSKLAAGALDSGAKIILGAEVEDLVVRDNPLRVVGVMFKWSAITAAGLHVDPLFALSRAVVDATGHEAVLVSILSRKNRVAGVAVPGE
RSGFAERAERDVVEYTGRMVPGLYVAGMSVAAVHGLHRMGPIFTGMLLSGRKVAEAIARDLGVPQ

>Acetomicrobium thermoterrenum DSM 13490
MKLDELVITKAIVEGYFKKLMNCLELDVAIVGGGPSGLVAALELAKAGKKVALYERKLSVGGGMWGGGMLFNEIVIQHEAKEILEGVGVNVRPYEVE
GYYTADSVEAVSTLTSKAVKAGATIFNALSVEDVVVDDEERINGLVVNWTAVEMAGLHVDPLSIHCKYVIDATGHDTEVVRVVARKMPGRLFTATGN
IEGEKFMSPDRAEKLTIVNTREVFPGLYVAGMAANATFGGPRMGPIFGGMLLSGVKAAREILSKI

>Acidianus hospitalis (strain W1)
MQSIRIKQVNEVKISKYILKYTFEDWNNLVESDVVIVGAGPSGMTAAYYLAKAGLKTVIFERRLSFGGGIGGGAMNFHKIVIETPADEIIKELKIRY
IEPEEGIFIIDSAEFMAKLATAAIDAGAKIIHGVTVDDVIFRENPLRVAGVAVEWTSTQMSGLHVDPLFISAKAVVDATGHDAEIISVASRKVPELG
IAVPGEKSAYSEIAEELVVENTGKVAPGLYATGMAVCEVKSLPRMGPIFGAMILSGKKVAEEIIKDLRNS

>Acidithiobacillales bacterium SM23_46
MCCQSLEAWRSEPEERKRNADVVVVGAGPSGMTAAIHLARERHRVILLEKRLSPGGGIWGGGMAMSEAIVQDDALPWLDDLGVRHKPSRGGLHSADA
VELAAALCLKTVQSGTVLFNLLTVEDVCIHQDRVTGVVVNRSMIAGALPVDPIAFRTNAVIDATGHEAVVVEAVHKRGLLAHPAVAKPLGEGPMDAA
SGEAFVVENVKEVYPGLWICGMSVCATLGGPRMGPIFGGMLLSGQRVAALVSSALTEFAQKDRESRK

>Aciduliprofundum boonei (strain DSM 19572 / T469)
MLDEVEITKLIVENYMKDLMEYADLDVAIVGAGPSGLTAAYYLATAKKKVAIFDRRLSIGGGMWGGGMMFNKIVVQEDAKHILDDFSINYERFGDYY
VADSVHSVTSLAYHATKEGAKIFNLIGAEDVIIKNNRVSGLVINWSVIGELPIDPLSIYAKYVIDATGHESEVIKTLVRKNNIKLNTPTGSIEGEHS
MDADTAESVIVDNVKEVYPGIFVTGMAANAVFGSPRMGPIFGGMLLSGKKVADEIIRRLS

>Actinobacteria bacterium HGW-Actinobacteria-3
MPLSEIEVTRGILEGFSRDFLSSLQSDVAIAGAGPSGMVCAYYLAREGLKVSVFERNLHVGGGMWGGGMLFPRIIIQEAAREIVEEFGVRLKPFKEG
YFVGDSVETVSKVTAAAIDAGVRVWVGVSVEDVLIREENRLAGVVLNWRAVELANLHVDPLAVEAKVVVDATGHEAGVVRTVARKIPGCRLNTDTGG
VIGEMPMWAQVGEELIVGNTREVYANLLVTGMAANAVYGAPRMGAIFGGMFLSGYKCAHLAADIVRKA

>Alistipes inops
MIETKVSQGIVSTYFDKLQKNLELDVAIVGGGPSGIVAAYYLAKAGLRVAQFDRKLAPGGGMWGGAMMFNQIVIQEEAMDIVREFGINYAPFGEGLY
VMDSVESTSALLYHAVHAGATVFNCYSVEDVVYKENRVSGVVVNWTPVLREGLHVDPLNILARVVIDGTGHDSEIAATVARKNGARLNTETGGVVGE
RSLDVTAGEDEVVKGTKEIYPGLYVCGMAASAVSGTPRMGPIFGGMLMSGKKVADEIIARLKK

>Ammonifex degensii (strain DSM 10501 / KC4)
MAGGAIDERLVSRAIIQTYSEELLQLTDFDVAVVGAGPSGLTAAYYLAQGGLKTVVFERRLSVGGGMWGGAMMFNYLVFQEEARPIFETMGVRYREY
QPGYYVAHSVEAVAAFTLAACRAGARIMNLITVEDLVLRDNRVAGLVLNWTAVDMAGMHIDPLAVHCRYVVDATGHDAEVVRILTQKNQVTVKVPGG
HVQGEKSMWSERGEKQTLDHSGEVFPGLYVAGMAANAVAGGYRMGPIFGGMVLSGKKVAELILEAHRREKSQTL

>Ancylomarina sp. 16SWW S1-10-2
MEQIVSAGIVDSYFKKLKENLSVDVAIVGGGPSGLVASYYLAKKGFKVALYETKLAPGGGMWGGAMMFNEIVVQKDALHILDELNVSYTNYQGDYYT
LDSVHATSALIYHATQAGVKIFNCSSIEDVVFQNNKVCGVVLNWSPVRREGLHVDPLVIMAKAVVDGTGHECDIVSTLERKNGVKLNTKTGKVMGEC
SLSIDEAERTTVENTKEVYPGLYVSGMASNGVSGGFRMGPIFGGMLLSGEKLAGLIAENLSK

>ANME-2 cluster archaeon HR1
MELDEITITRAIIEDFTSDFLQSIDTDVALVGGGPANLIAARTLARAGVKTVLFERKLEVGGGMWGGGMMMPRIVVQEEARHILDDLGVRYRKYEEG
YYVADSIECTGKLIYEAASSGASIYNLISVEDVMIREGDAVTGLVINRTIVDMQKLHVDPITIRAKVVIDGTGHDAEICTTLSRKIPGALHVAGEKP
MWADVAERIILDNTKEVYPGLIVTGMAANAVAGAPRMGPIFGGMLLSGEKAAQIAIAKLGL

>Archaeoglobales archaeon ex4484_92
MEARISKAIIEEVAKDWSNISQVDVVIVGAGPSGLTAGKYLAEKGLKTLILERRLCFGGGIGGGGMLFHKIVIEKFAREILDDFDVRYYEHDNLLVA
DVAEFMAKLAVGCVNAGTKIIHGVSVCDVIFRLEPIRITGVCIQWSAVELSGLHVDPMFIESKAVLDATGHDAEVVSIASKKVPLDLNVTGEKSAYA
ELGEKLVVEKTGKVVEGLYATGMAVCSVFNLPRMGPIFGGMLQSGKKAAEIIYNDLK

>Archaeoglobus fulgidus
MEAEITKAIVETASEEWVEYAESDVIVVGAGPSGLTAARYLAEKGLKTLVLERRLSFGGGIGGGGMLFHKVVVEREAKDILDDFGIRYTEHRNFLVA
DSAEFMAKLAAKAIDAGAKIIHGVSVEDVIFRDDPLGVRGVCIQWSAVEISGLHVDPLFLRSRAVVDATGHDAEVISVAARKIPLEVSVVGERSAYS
EVAEREIVEKTGKIVKGLYAAGMAVAAVHNLPRMGPIFGGMLLSGKKVAEIVAEDLKL

>Armatimonadetes bacterium
MSFEKDSFKWDELTVTRGIVETFMADFLDSIDLDVAVVGAGPSGITAARILAGQGHRVGIFERNLHIGGGIWGGGMLFPRIVIEEEAAPLMEAAGVK
LRPWKDGTVIADAVESATKMTAAAIDAGARIFVGIEAEDVVVDDSDRVCGVVINWGAVTAAKLHVDPLAVHSKVLIESTGHPCEVGDVLLRKIPGAR
LDTGETCVPGEASMNARAGEAALIANTREIYPGVVVAGMAANAVSRSPRMGAIFGGMLLSGQKAAEISAQIIADLG

>bacterium (Candidatus Ratteibacteria) CG23_combo_of_CG06-09_8_20_14_all_48_7
MLDDVVISRAIVETYFQDLLNYLENDVAIAGAGPAGLTAAYFLAKKGRKVAIFERQLRVGGGMPGGGMMFNKIVIQEETKKILDEFGIRYQKYQNGY
YVADSLETTSLLTSKAIQAGAKIFNLIAVEDLSLQEGRVNGLVLNWSAVKTAGLHVDPITIRAKAVVDATGHDAVLCRLLVDKGKVTLRTPTGKVAG
EGPMWAEKGEEMIIANTGEVFPGLFIAGMTVNAVCGGPRMGPIFGGMLLSGERLSLLIP

>bacterium (Candidatus Stahlbacteria) CG23_combo_of_CG06-09_8_20_14_all_40_9
MMLDETIISRAIIETYKEKFVNILKSDVAIAGGGPSGLIAGYYLKKKKPDLKVVLFERKLSIGGGMWGGGMMMNEIVVQEEGKKILDEFGVKSKKFE
NGYYTADSIETVSALALNTVKAGVTILNAISVEDTIIEDNIAKGLVINWTSALDIGLHVDPLALRADHIIDATGHPCEIAHLIEKKGKKLFTRTGKI
IGEGAMYADKGERVIIENTKELFPNVWACGMAANAVFGGPRMGPIFGGMLLSGKIVAEKILQKN

>bacterium 42_11
MKDILISKAILESFFNKLRDSLELDVAIVGAGPSGLVASYELAKKKVKIAIFEERNTPGGGIWGGGIMFNEVVLEKELEDFLKELDIKYKYVEDYIV
VDSTHFASALIYHTTTWGTRIFNSISVEDIAMQNNRVCGVVINWGPVKKLGLHVDPITIKASYVVDGTGHPANVVSLLVKRGLLEKKTEFPMNAEEA
EKFVVEKTGEVFPGLLVSGMAVCEVYGGPRMGPIFGGMVLSGRRIAEIITERVNKR

>Bacteroidales bacterium 6E
MEQIVSSGIIDSYFKKIKESLSVDVAIVGGGPSGLVAAYYLAKKGLKVAMFERKLAPGGGMWGGAMMFNEIVVQKGALQILDEFKIDYTHYEGDYYT
LDSVQATSSLIYHAGKAGARIFNCTSVEDVVFHNNKVSGIVLNWAPVHRERLHVDPLVIMAKAVIDGTGHDCDIARILERKNNIQLNTVSGKVQGER
SLSIDEAERTTVENTKEIFPGLYVSGMAANGVSGGFRMGPIFGGMLLSGEKVAALIMNQLNK

>Bacteroides cellulosilyticus
MIETKVSKGIISTYFEKLERNLDLDVAIVGGGPSGIVAAYYLAKAGLKVAQFDRKLAPGGGMWGGAMMFNQIVIQEEAIDIVKEFNINHEKYEDGLY
VMDSVESTSALLYHAVHAGATVFNCYSVEDVIFKNNTVSGVVVNWTPVLREGMHVDPLNILAKIVIDGTGHDSEIAATVARKNGSRLATETGGVIGE
RSLDVIAGEEEVVNGTKEIYPGLYVCGMAASAVSGTPRMGPIFGGMLMSGKKVAEEIIAKLKK

>Bacteroides sp. 3_1_19
MEQIVSSGIIDSYFEKLKSNLSVDVAIVGGGPSGIVAAYFLAKAGKKVALFDRKLAPGGGMWGGAMMFNDIVVQEEAMPIIKELGVSYKEGANGTYI
MDSVHTTSALIYQATKAGATIFNCYSVEDVVFHNDAVAGVVVNWAPVIREGMHVDPLTIMAKAVLEGTGHDCEIARVVARKNDIQLNTPTGGVIGER
SLNVELGEQTTVENTKEIYPGLFVSGMAANGVSGSFRMGPIFGGMLMSGKKAAELICEKLGN

>Bacteroides stercorirosoris
MIETKVSKGIISTYFEKLERNLDLDVAIVGGGPSGIVAAYYLAKAGLKVAQFDRKLAPGGGMWGGAMMFNQIVIQEEAIDIVKEFNINHEKYEDGLY
VMDSVESTSALLYQAVHAGATIFNCYSVEDVIFKNNTVNGVVVNWTPVLREGMHVDPLNILAKVVVDGTGHDSEIAATVARKNGIRLATETGGVIGE
RSLDVVAGEDEVVNGTKEIYPGLYVCGMAASAVSGTPRMGPIFGGMLMSGKKVADEIIAKLKK

>Bacteroidetes bacterium ADurb.Bin035
MEQIVSNGIIDSYFNKLKQYLSVDVAIVGGGPSGLVASYYLAKNNYKVAIYERKLAPGGGMWGGAMMFNEIVVQKAALHILDELSIEYREFEHDFYV
IDSVHAASALIYNASKAGVKIFNCTSVEDVVFLDNKVCGIVLNWAPVAREHLEIDPLVIMAKVVVDSTGHDCDVAHTLERKNNIKLNTETGKVIGER
SLSINEGENSTIDNTKEIYPGLFVTGMAANGVSGSFRMGPIFGGMILSGKKVADLIINKIKNT

>Brockia lithotrophica
MFDERVITRAIVETYLEEFRSIVDLDVAVVGSGPSGLVAARELARRGYRVAVFEKRLSVGGGLWGGGMLMNRIVVQEAARPILEEFGVRTREYAPGV
YVAHSVETVAALVFGALQAGAYVFNGVAMEDVVVRDGRVAGLVLNWGAVHAAGLHVDPLAIHARAVLDATGHDAEVVRKLAAKNGVSLSVSGERSMW
ADRGEEEAVALTGEVYPGLFASGMAATNVYGGHRMGPIFGGMLLSGVRAAELIAEALGS

>Caldimicrobium thiodismutans
MELQINQAIIREGMRDLEDFSDVDVLIAGAGPSGLTAAKYLAERGFKVLIYERRLSFGGGIGGGGNMIPKIVVQTEALPIVEDFKIRAKKVENGFFT
IDPAELIAKLATGALDAGAKIFLGVNVDDVIVRDAPPRVVGVLWHWTAIQLSGLHVDPLYTHCKALVDATGHDAELIAIAGRKNPELGIEMAGEKSN
WSEVSERLVVEHTGRVAQGLYVTGIAVCAVYGLPRMGPIFGGMLMSGKKLAEIIEKDLKNASKGKGKGKKAG

>candidate division MSBL1 archaeon SCGC-AAA259E19
MLDEEVITKAIVEEYMRTFLENTDVEAALGGAGPANLVAAKKLAEKGVKTAVYEEKLNVGGGMWGGGMMYPRIVVQEEAKRILKEFGINFSEYEKGY
YVASSIESVARLASEAVKAGAEIFNLTKVEDVMVREDDHIAGVVLNWSAVEKANLHVDPLTVKADVVIDGTGHDAEICRVTQRKIPDADLQVRGEKP
MWAEKGEKALMDTTKEVYPGLIVSGMAANAVSAGPRMGPVFGGMLLSGEKAAELALERIEEK

>candidate division TA06 bacterium DG_78
MIEETIISKAIVESYLKDVLDSIESDVIIAGAGPSGLCAGYYLAKHKYKTVLFERGLKLGGGMPGGGIMFNKIVVQDEGKATLDEFGITYHQYEPGY
FVADSLETTACLTEKALKAGLKIFNLVSVEDVMIREGCVTGVVINWSAVEMARLHIDPISFKSKVVIDATGHPSEIVHIVEKKSGGKLLTPSGRIEG
EKSMWAEVAERLSLENTKEVYPNLYVCGMAANAVFGGPRMGPIFGGMLLSGKQVAELVMKKV

>candidate division WOR_3 bacterium SM1_77
MIEETVISRAIIESYLHDLLDSIQSDVIIGGAGPSGLCASYYLAKKGHKVVLFERALKLGGGMPGGGIMFNRIVVQETARRILDEFAIRYEEYKPGY
YVANSLEATAALTDKAIKAGAKIFNLITIEDVVVREQVVSGVVINWSSVEMAKLHIDPISFESKFVIDATGHPSEIARIVERKNSGTLLTPSGKIEG
EKSMWAEVAEQFTVENTKEIFPNLYVCGMAANAVFGGPRMGPIFGGMLISGMKVAELVMKKI

>candidate division Zixibacteria bacterium 4484_93
MNFTDIKISKAIIDDYFRKLTSSLELDVAIVGAGPAGLTAGYFIAKQGYRVSLFERKLSAGGGMWGGGMMFNTIVVGKGANDILDEFGIRYRRTEDS
LYIADAVESVGALIYRSTQAGLRIFNLITVEDLLVKDGKVEGLVINWSPVEMASLHIDPLTVQSRYSVDATGHPLEVVRTLCKKSGVQLFTETGDVL
GERSMCADEGEKFVLEKTGEVAPNMFVAGMAACAAFGGPRMGPIFGGMLLSGKKVADIITKRLKEKK

>Candidatus Acetothermia bacterium
MKIDDVLVSRLLIEEYMADFLDCLHIDVAIVGAGPAGLTAAYYLAKAGAKVAVYERKLAIGGGMWGGGAMFSRIVIQEEAKQILDQFKITSIAKEDG
YYVADAIEAITLLAAGAIQAGAKVFNLIHIEDLLLRNDRVEGLVLQWSPVEMGGLHVDPITIGAREVIDATGHDCEVVKKLLHKGGVKIDTETGGML
GERPMWAEKGEKMTVQYTKQIYPGLYVAGMAVGAVFGTPRMGPIFGGMLLSGKKAAQIIAQTLS

>Candidatus Acidianus copahuensis
MKVKQVDEGKISKYILKFTFEDWENIIDSDVIIVGAGPSGMTAAYYLAKAGLKTVLFERRLSFGGGIGGGAMLFHKIIIESPADEILKELGIRLVKA
EEDVYAVDTAEFMAKLASSAIDAGAKFIHGITVDDVIFREEPLKVAGVAVEWTATQMSSLHVDPIFISAKAVVDATGHDAEVISVASRKLPELEISI
PGEKSAYSEVAEQQVVDGTGKVAPGLYAAGMAVCEIKGLPRMGPIFGAMVLSGKKVADEIINDIRKS

>Candidatus Altiarchaeum sp. CG2_30_32_3053
MIEETKITELIVRNAVDDFLNNLDVDVVVAGGGPAGLTTARYLAKAKKKVVLFERKLSIGGGMWGGGMMFPRVVLQKGGKEILEECDVKCKKSGGLW
VADSIECVTKMTAKAIDEGVKIFNLVSIEDVIIRNNEDNKNNKNRKNKTKICGVVINWTAVQMANLHVDPLSVKSKFVVDATGHEASISHLVVKKVG
NLNTKTGDVLGERSMWAEKGESDIMGNTGEVYPGLFVSGMAANAVYGSERMGAIFGGMLMSGKKVSELILKKI

>Candidatus Aminicenantes bacterium 4484_214
MIDEIVISRAITEAYLKEFLDCLESDVIISGAGPAGLCAALNLAQEGYKVVVFERTLRPGGGVPGGGMMFNKIVIQEEARPLLEELDVTLKPYQENY
YVVQALELLGALLVKAIKQGVYLFNCISVEDVLIYDKKVSGVVINWSAAQAAGLHVDPLTARTKFVVDATGHAAEVAEIVSRKSGCKLFTSTGKVTG
EKPMWAEEGEKILLTNTKEVYPHLYVCGMAANAVFGGPRMGPIFGGMLLSGQKVAQLIAARLKTK

>Candidatus Aramenus sulfurataquae
MQSIRIKQVDEVKISRYILKYTFEDWYSLVDSDVVVVGAGPSGLATAYFTAKAGLKTVVFERRLSFGGGIGGGAMNFHKIVIESPADELLREWKVKL
VEAEEGVFIVDAAEFMAKLGAAAIDAGAKVIHGINVDDVIFRDKPLRVAGVAVEWTSTQMSGLHVDPLFVSAKAVVDATGHDAEILSVASRKIPELG
IVIPGEKSAYSEVAEELVVNNAGKVAEGLYTTGMAVCEVKSLPRMGPIFGAMVLSGKKVAEDIINDLRNS

>Candidatus Aureabacteria bacterium SURF_26
MPLDDLIVSRAIIDEYHTTLVDALNMDVAIVGGGPAGLVAGYYLAKQGYKVSLFERKLSIGGGMWGGGIMFNKIVLQEDALKVLNEFNVRVKKYRDN
YYVADSVETVGALIYHATQAGLQIFNCMSIEDVKITADDAVCGLVVNWTSVELTNMHVDPITFGAKFVIDATGHSCDMANIILKRIGKVLFTPTGDI
MGEGSMNAELAEKVVAENSREIYKNLYVTGMAANAIWGSKRMGPIFGGMLLSGKKVADDISARLAQEAVNA

>Candidatus Bathyarchaeota archaeon B26-2
MRIKEVDEAVVTKAILEGSLKYLHELTEVDVAVVGAGPAGLTASRYLAKAGLKTVVFERRLSFGGGIGGGGMQLPMLVVQSPADEILREVGCNLTTY
REGVYLANSSEMLAKLAYSAVKAGAHIILGVTVDDLIYRSEENRTRIVGVVVQWSSVIISGLHVDPLAFKAGAVVDCTGHDAEVLSVASRKIPELNL
MIQGEKAMWVSESERLIVEKTGEVSPGLYVAGMAVATLNQTPRMGPIFGGMLLSGKKVAEIIIERYKQNRTQG

>Candidatus Desantisbacteria bacterium CG2_30_40_21
MQLDDVVISRAIIESWNKDLLDSLDIDVAIVGGGPAGLTCGYYLSKKGLKVVLFERNLSIGGGMWGGGMMYNKCVFQQESLPILNEFGIRTQEYQDG
YFITDSLETVTTLCSGALKAGLKIFNLIGVEDVMIRQEGVTGLVLNWTAVTMAKLHIDPLTIRAKAIVDSTGHAAEVAGIIVRKIGKKLLTETGEML
GEKPMWAEVGERTIAENTKEIYPGVFVAGMSANAVFGGPRMGPIFGGMLLSGKQAAEIIAARL

>Candidatus Desulforudis sp.
MKIDETVVSRAIIERYTQKLLSCLDVDVEIVGGGPAGLTAAHYLAKHGVKTTLYERKLSVGGGMWGGAIMMNEIVFQEQARPLFEEFGIRINPYSDG
YYTASSVECVAALTLQACQAGANIINLMTVEDVVLHEDRVSGLVLNWTAVDIAGLHVDPIATRSKYVIDCTGHDMEVANILSCKAGVKLVTPSGEPV
GEKPMWADVGENQLIGHTIEVYPGLFVAGMAANAVNGGYRMGAIFGGMVLSGRRAGELILKRLQS

>Candidatus Fermentibacteria bacterium
MENIVTSAIAEDYHRKIQESVTAHVAIAGGGPSGLVAAAELASRGLSVNLYEKNLTPGGGMWGGAMLFNSILIEEEFAETAEELGMKLRKYRDSVLL
ADSVQATAALISRACSAGVRMFNGMAVEDVTVDRYDRVNGVVVNWAPVMKLGMMVDPLMTCAGAVLDATGHPAEIVTRFAAKNNTEITVPGEASLNV
EMGEKHTVEHTGMVHPGLFVSGMSACATAGGYRMGPVFGGMLKSGLKAAEEITEYLNKSQ

>Candidatus Korarchaeota archaeon
MFPVEESVISSAIIERGSKFLVDLVKSDVIIVGAGPSGLVAGRYIAKTGLKTCIIERRLSFGGGIGGGGMLFPRIVVQEPAQEILEEVGVKLEPYSK
GVWIADVAETIAKLAAGAIDSGARILLGANVEDLIVRNSRVCGVVVQWSAVTSAGLHVDPLAFESRAVIDCTGHNAEVVAIAARKNPELGIRVLGEH
SMDAVRAEKEVVELTGEVLPGLWVAGMAAAAVRGGPRMGPIFGGMLLSGKKVAELVSRELEVL

>Candidatus Nitrospira inopinata
MHKPKPAPLRERDVTRHIAREYYKEFDQLIESDVIIVGAGPSGLLCAHDLAAMGFRTLIVEQSLALGGGFWHGGYLMNKATICEPANEILEELGVPC
KRIADCDGMYMVDPPHATGALVAAAYRAGAKILNLTRVVDLILRQDGLLEGIVVNNTTAEMAGHDVIHVDPIALESKIVVDATGHDAVVVELLHKRN
LYKPVPGNGAMWVSRSEEEVVDRTGEVYPNCFVVGLAVAAVYGTPRMGPAFGSMLLSGRYGAQLIKKKLKQE

>Candidatus Nitrospira nitrificans
MAKPRPAPLRERDITRHIAREYYKEFDQLIESDVIIVGAGPSGLICAHDLAAMGFRTVLIEQSLALGGGFWSGGYLMNKATICEPANEILEEVGVPC
KKIKECEGMYMVDPPHATGALIAAAYKGGAKIMNLTRVVDLILRNGGLLEGVVVNSTTAEMAGHDLIHVDPIALESKIVVDATGHDAVLVELLHKRN
LYNKVPGNGAMWVARSEEEVMDRTGEVYPNCFVIGLAVAAVYGTPRMGPAFGSMLLSGRYGAELIKKKLKQE

>Candidatus Nitrospira nitrosa
MTKPRPAPLRERDITRQIAREYYKEFDQLIESDVIIVGAGPSGLICAHDLAKMGFRTLIIEQSLALGGGFWHGGYLMNKATICEPANEILEEIGVPC
KKIRECEGMYMVDPPHATGALIASAYKAGAKVLNLTRVVDLILRRDGILEGVVVNNTTAEMAGHDVIHVDPIALESKIVVDATGHDAIVVELLHKRN
LYQKIPGNGAMWVSRSEEEVMDRTGEVYPNCFVIGLAVAAVYGTPRMGPAFGSMLLSGRYGAGLIAKKLKNE

>Candidatus Syntrophoarchaeum butanivorans
MDEVTISKAITESYMKDLIDSMVLDTVVVGAGPAGLLAAYNLAREGVKVAVFERRLSVGGGMWGGGMMFSRIVVQDAGREILDEIGVRCSEYEPGYY
IADAIEAVTTITSEVIRAGARIFNLMSVEDVVVRDDDRIHGVVINWSAVELSKLHVDPMTVIADYVIDATGHAAEVARIVEQKLGGGALTVRGERPM
WAEAGEAAVVENTKEIYPGLIVAGMAANAVLGSPRMGPVFGGMLLSGRKAAELVLSRL

>Chloroflexi bacterium RBG_13_51_52
MVKFSPVGEVVITRAIVEEFAKEFNEYVESDCIIIGGGPSGLVAGRDIARAGKKVVIIERNNYLGGGFWSGGYLMPKVTVRHPGEKILDELGVPYKT
VAKGLVVCDAPHACAALIAAACAAGVKIFNMTMLEDLVVKDGRVCGAVINWSPIASLPRQVAALDPVAIEAKVVIDATGHDATVVAKLEKRNLIKMK
GEGAMWIEKSEDLIVEHTGECFPGLIVTGMAVGAVYGLPRMGPTFGSMFLSGEVAAKVALEKMK

>Clostridium drakei
MYLEDTKISKAIIDTYKDKLEDILHSDVIIVGGGPSGLVGASYLAKAGIKTTLLERNLSIGGGMWGGGMMMNQIVIQESAKSILDEFNIGYKKYEEN
YYTADSIECVSALTLSASQSGARILNSISVEDVIVKDKCISGLVINWTAVEKTRMPIDPIMIESKYVLDATGHDASVVNKLVTRMGNVLNTPNGTLE
GEKPMWADRGEEQVIKNTREVYPGLYVSGMAANATFGGQRMGPIFGGMLISGQKVAQELIQKIKNC

>Clostridium ragsdalei P11
MYLEDTKISKAIIDTYKNKLEDVLHSDVIIVGGGPSGLVAASYLAKAGIKTTLLERSLSIGGGMWGGGMMMNQIVIQESAKSILDESNIGYKKYEEN
YYTADSIECVSGLTFNAAQAGARILNLITVEDVIVKDKCISGLVINWAAVEKTRMPIDPIMIESKYVLDATGHDASVVNKLVTRMGNVLNTPNGTLE
GEKPMWANRGEEQVVENTREVYPGLYVSGMAANATFGGQRMGPIFGGMLISGQKVAQELIKKIKNC

>Clostridium sp. JN500901
MYLEDTKISKAIIDTYKDKLEDILYSDVIIVGGGPSGLVAAAYLAEAGVKTTIIERSLSIGGGMWGGGMMMNQIVIQESARSILDDFNVNYKKYEEN
YYTVDSIECVSALTLKAVKAGAKILNLISVEDVIEKDNCIAGLVINWAAVEKTKMPIDPIMIESKYILDATGHDASIVNKLVAKQGNVLNTENGSFV
GEKTMWADRGEDQVVKNTGEVYPGLYVSGMAANATLGGQRMGPVFGGMLLSGQKAAKQLIEKLHAAK

>Coxiella sp. DG_40
MEQITTLGIVDSYYQKLKDNLFIDVAIVGGGPSALVAAYYLAKLQKKVAIFERKLAPGGGMWGGGMMFNQIVVQSEALSILDEFKISYALFKDNYYL
VDSIESTASLIYHTIHAGAKVFNCYSVEDIVLKNKKVSGIVVNWGTVHDQGLHVDPLVVVAKCVIEATGHSCEVAKVLAKKNGIKLHTETGGVVGEK
SLAMEQAERSTIENTKEIYPGLYVCGMAANGVSGDFRMGPIFGGMLMSGKKVAEIIVKDIT

>Dehalococcoidia bacterium
MPLFHPVTEGEITRAIVNSFLRQFEEYVSSDVIIVGGGPSGLMAGRELGKAGLKVIVIERNNYLGGGFWAGGYFMNKLTLREPAQEVLDELGVPYSR
AGEGLYVADAPHACSKLIGAAADSGVKFFNLTLLEDLVVREDKRVAGAVINWSPIAYLPREIAALDPVPLETKVIIDATGHDASVARKLERRGMLKL
AGEGALWIEESEEAVVEHTGEVYPGLVVTGMAVASVYGLPRMGPTFGGMLLSGKRAAEVALAAVLTDSR

>Desulfacinum hydrothermale DSM 13146
MALDERIITRAIMDRYIAKLKEAIDLDVAIVGAGPSGLVAGMLLAEAGKKVALFERKLSVGGGMWGGGMLFNEIVVQEEAKTILDQVGIRAQHYTDG
YYTADAVESVSTLTSRSVKAGARIFNCVSVEDVMMRPERIMGLVLNWSAVEMAGLHVDPLAVRCQVVVDATGHDTEVVKVVERKVPGSLSTPSGKRA
GERSMWAEEAERLTLENTCQVYPGLYVAGMAANATFGGPRMGPIFGGMLLSGQKVARLILEQLQS

>Desulfarculus sp.
MLEEVTITRAIIRRYLGKLDQSLELDAAIVGGGPAGLVAGKKLAQAGYKTALFERKLSVGGGMWGGGMMLNEIVVQQEARRILEEFGVPSSEFAPGY
YTADSVLATSTLCSVAAKAGLTIFNLVSVEDVVIRAQRVTSLVINWSAVQMAGLHVDPLTIKARWVIDATGHDSEVLHVIARKVDAELLTASGKVMG
ERSLWAEQAESDTLANTREAFPGVYTAGMCANAVFGSYRMGPVFGGMLLSGEKAAAEVAARLAAGE

>Desulfatibacillum aliphaticivorans
MEERITSAIVRTYFEKLQNFLEVDLAIVGAGPSGLVAAAALAKEGKKVAIFERLLAPGGGVWGGGMLFNEIVIQEEALHILDDFNISYKSAGDGLYT
ADSVEVASGLIFGAKKAGVMINNAVSVEDVVCREGRICGVVVNWTPVERLGMHVDPLVVMSKAVLDGTGHPGEITDLATRKAGIKIDTPTGKIMGEK
PMWMELGEASTVENTKCLYPGLYVSGMAANNASGGFRMGPIFGGMFMSGRKVAKMILEDIDG

>Desulfobacca acetoxidans (strain ATCC 700848 / DSM 11109 / ASRB2)
MGLDEIIISRAIIERFMEKFLDNLELDVAIVGGGVSGLVAGWRLAQKGRKAAIFERKLSVGGGMWGGGMMFNEIVVQEEAKHLLDELGITSRPYDRG
YYTADAIESTTTLASQAMKAGVKIFNLIHVEDVMVRENRIDGLVILWTAVNMAGLHVDPLTIRAKHVIDCTGHDVEVIKIFLRKNQPASLKTETGGI
MGERSMWAEVGEAKTVEYTSEVYPGLWVAGMTATGTLGTFRMGPIFGGMMLSGEKAANLIDERLKKG

>Desulfobacteraceae bacterium
MQLDDVAISKSILDAYFEKLLARLDVDVALVGAGPANLVAGYYLGKSGFKAVVFESKLAPGGGMWGGGMMFNEIVLQDDAVHIAEELGIHCNPGGDG
YYTMDSVEATSTIISRCVRAGTVIFNLIKVEDVLFRQDDRQPRVSGLVINWSPVEKLGLYVDPLSIRASFVVDGTGHPADICRTVARKMDVKLNTKT
GNVVGEMPLWAEKGEQFTVTNTAEVFPGLYVAGMAANNAFGGPRMGPIFGGMLRSGKKVAEILAEKLRSD

>Desulfobacterium sp. 4572_20
MAINEVVISKAIIDRFSQKFMEYTEVDTAIVGAGPSGLIAAYFLARAGQKVALFERQLSIGGGMWGGGMMFNEIVVQTQGKELLEMFGISAREYEPG
YYTADAVECVTTICSNAVKAGAKIFNCMSVEDVSIREDRVMGLVLTWSAVEAARMHVDPLTIAAKYVIDATGHDTEVIRLIEKKADIALQTETGKIM
GERSMWADKAEQLTIENTKEICPGVFVSGMAANAAFGGPRMGPIFGGMLLSGKKVAELIMAKEGSAFETSAEDFDSWFNRNQAIFESELLAQQQFIY
SL

>Desulfocarbo indianensis
MLDEITITRAIIDRYFEKLNRNLELDAAIVGGGPSGLIAGYKLAKAGYRVAMFERKLSIGGGMWGGGMMMNEIVVQEEAKRILDEVQVPTREFQPGY
YTADSVLCTSTLCSQAAKAGLTIFNLVSVEDVMVREQRVVGLVINWTAVEMAGLHVDPLTIRAKYTIDATGHAAEVMHVIARKVDAKLFTDDGKVAG
ERSLWAEVAETNTVNNTREAFGGVFTAGMCCNATFGSYRMGPVFGGMLLSGEKAAQLVAERLQAEK

>Desulfococcus oleovorans (strain DSM 6200 / Hxd3)
MELNEVTISRAIIDRFYEKLIANLEVDVAVVGGGPSGLVAAWRLARAGRKVALFERKLSIGGGMWGGAMLFNEIVVQKSALHVLDAMEIGYRLYAED
YYTADAVEAISTLTSQAAKAGVAIFNCVTVEDVMIRPDRIVGLVLNWSPVEMAGLHVDPLAMRASFVIDATGHATEVVHVVAKKVPGTLRTDSGKIE
GEKSMWSDRAESLTLENTREVYPGLYVAGMAGNATFGGPRMGAIFGGMLLSGEKVAAEILERLE

>Desulfofundulus australicus DSM 11792
MMHLEDVVISKAIISRYQEELLEALESDVAVVGGGPSGLVAAYYLARANKKVVLFERKLSIGGGMWGGGMMFNQIVIQDEALPLLEEFKISYRVFEE
GYYTASSVEAVAALTLGAVRAGAKIFNLISVEDIMVRDNRVAGLVINWTPVDLGRLHVDPLTVQSSYVIDCTGHDAQVAGMIVKKMGAVLKTTTGTL
EGEKPMWAARGEMATVANTREVYPGLIVAGMAANAVCGGHRMGPVFGGMLLSGQRAARIILEGDKT

>Desulfofustis glycolicus DSM 9705
MLNEVTISTAIINRYMTKLTSALDLDVAIVGGGPSGLVAGYYLAKAGRKVALFDRKLSIGGGIWGGGMMFNEIVVQEAGAAVLAEFGLAGSPFEPGY
YTLDSVYTTATLVHKAMAAGLLIFNLIGVDDVVIKDERVAGLVINWGAVSTLGWHIDPLTLFARYVLDATGHDAEIASVLVRKMGVRLNTETGGLVG
EKSMAAERAERETVTNTREVYPGLFVSGMAANAVCGGYRMGPVFGGMLLSGKRAAESILEGLA

>Desulfonatronospira thiodismutans ASO3-1
MALDEIIISRAIIETYTKKLMDSLELDVAICGAGPSGMVAAYYLASAGKKTAVFERNLAPGGGMWGGGMMFNEVVVQEEAREILDELDIKSVEYTPG
YYTADSVEAVCTLGSKAAKAGARFFNLVCIEDVMIRENRITGLVINWSAVESAGLHVDPLTVRADYVVEATGHPVEIMQVIESKMDTRLNTPSGRLE
GEKSMWAEKAEEHTIENTTEAFPGVYVCGMSANATFGSFRMGPVFGGMLRSGKKVAQEIINKAK

>Desulfonauticus sp. 38_4375
MSLDEKIISEAIISKYFEDFKRCLNLDVAIVGGGPSGLTAAYHLAKEGFKVALFERKLSIGGGMWGGGMTFNYIVVQEQGKQILEEMDIICEEYKPG
YYVVDAVLATTTLASKACKAGAKIFNCMSVEDVVIREEGKVKRVAGLVVNYSPVEIAGLHVDPLVLETKFVIEATGHDTEVLKTLVRKNDIKLFTPS
GGIEGEKSMWAEVAEENTLKNTREAFPGIYVCGMAANACFGSYRMGPIFGGMLLSGVKVAEEISTRLKEGK

>Desulforudis audaxviator (strain MP104C)
MKLDETIISRAIIESYVTRLLSCLEVDVEIVGGGPSGLTAAYYLARAGLKTTVYERKLSVGGGMWGGAAMMNEIVFQETARPVFEEFGVTIKKYRDN
YYTASSVECVAALTLGACRAGANIMNLLTVEDVVLHNNRVSGLVLNWSAVEISGLHVDPIATRSKFVVDATGHDVSVVGVLARKAGVQLDTPSGKVQ
GEKPMWADLGEAQIMENTSEIFPGLYVVGMAANAVHGGYRMGAVFGGMVLSGRRVAEMIIDRLKV

>Desulfovibrionaceae bacterium CG1_02_65_16
MIIDERIVSEAIASTYFGKFKSCLDLDVAIVGGGPSGLTAAWKLAKAGRKVALFERKLSIGGGMWGGGMTWNSIVVQESAKSILEDAGVALSEFKPG
YFTADSVAATAALAYQATHAGAHVFNCMSVEDVVLREVEGVKRVIGLVVNSSPVEIARLHVDPLVLHCKHAIECTGHDVEMLKTLVRKNDVRLDTPS
GGIEGEQSMWADVAEANTVRYTREVFPGVWVAGMAANAAFGSYRMGPIFGGMLLSGVKVAETIDALL

>Desulfovibrionales bacterium GWA2_65_9
MIIDERIVTEAIASAYFEKFKQCLDLDVAIVGGGPSGLTAAWKLAEAGRKVALFERKLSVGGGMWGGGMTWNYIVVQEEAKGILEEAGCAMSEYKPG
YFLADSVAATAALAYRATKAGAHVFNCMSVEDVVLREIDGEKRVMGLVVNSSPVEMARLHVDPLVLHCKHAIECTGHDVEMLKTLVRKNGVKLNTPS
GGIEGEQSMWADVAEANTVRHTREVFPGVWVAGMAANATYGSYRMGPIFGGMLLSGVKVAEEINARL

>Desulfurella amilsii
MALDERIISRAIIERYFQKLLANIDCDCAIVGAGPAGLVCGYELVKNGLKVTLFDKRLSVGGGMWGGAMMFNEIVVQEEGKLILDEFDIKCSLFEPN
YYTTDSIEAITTLISKTVKAGVKIFNGIEIEDVVLKKVDGDYKVGGVVINWTTVNMAHLPVDPIVISSSFTVDATGHDAHLAETLVRKGGVKLNTDS
GAVIGEKPMWAQIGEQDTVNHTKEIFSGLYVCGMAANAVSGAHRMGPIFGGMLNSGKKCAQLILEKWSRK

>Desulfurella multipotens
MALDERIISKAIIERYSQKLLSQLDCDCVIVGGGPAGLICGYELAKNGLKVTLFDKRLSVGGGMWGGAMMFNEIVVQEDGKAILDEFDIKTVLYEPN
YYTADSIEAISTLISKTVKAGVKIFNGIEIEDVVLKKVDGQYRVGGVVINWTTVNMASLPVDPIVVSASFTVDATGHDAHLAQTLVRKAGVKLNTDS
GGVPGEKPMWADVGEQDTVNHTKEIYNGLYVCGMAANACSGAHRMGPIFGGMLNSGKKCASLILEKWGKK

>Desulfurobacterium atlanticum
MELSEVVISRAIVERFMNKLLSNLKVDVAIVGGGPSGLVAAYYLAKEGFKVSLFERKLSIGGGMWGGAMLFNEIVVQEMGREILDEFDVGYEKFQEG
YYTTDSVEAVTTIASKAVKAGAKVFNGVTVEDVVLKKENGDYRVCGLVINWTPVEITGMHVDPLTIESKFVIDATGHDAYVVSTLQKKAGIRLDTKT
GCVVGEKPLWASVGEEDTVKNSREVYPGIYVSGMAANAVCGSHRMGPVFGGMLMSGKKIAKEIAERLKHNVEE

>Desulfurobacterium indicum
MENLSEVKISKAIIERFTEKLLSNLEVDVAIVGGGPSGLVAAYYLAKEGLKVSLFERKLSIGGGMWAGAMFFNEIVVQEMGREILDEFSVSYRKYDE
GYYTADAVEAVTTIASKAMKAGAKIFNGVTAEDVVLKKVNGQYRVCGLVINWSTVDMTGLMVDPLVVTSNYVIDATGHDATIVSTLQKKAGIRLDTE
TGCVVGEKPLWASVGEEDTVKNSREVFPGIYVSGMAANATCGSHRMGPVFGGMLMSGKKIAMEIAQKLKS

>Desulfurococcales archaeon ex4484_42
MVKELESRVTELIVKHASRDWAELASTDVVIVGAGPSGLTAAKYLAEDGIKVVIFERRLSFGGGIGGGGMLFHKVVVEDFALDILKDFGIRYVEDGG
LYVVDASELMAKLAVGALNAGAKIIHGVTVEDVIFRTNPLRITGVAIQWSAVPLANLHVDPLLIYSKAVIDATGHDAEVVRVASRKIPELRLKVTTE
KSAYSELGERLVVEKTGKVVPGLYVTGMAVAALNNLPRMGPIFGGMLLSGKKVANEVLKDLRT

>Desulfuromonas sp. SDB
MKDINITNHIIAEFYKDIQDRVSSDVIIIGAGPSGLVASYLLAQDNFKVTVFEKRNQPGGGIWGGGMMFNQLVLPDDLQDFLNQMSIKFKLHPDNLI
SVDSVHFSSALLYHATEVGVKIFNNIGVEDLLVVDDMVRGVVINWNDVIKNKIPIDPLTFEAKAVVDSTGHPADGVEKLARRGLVEISQEFPMNADV
AEKFVVEATGQLYPGLYVSGMAATAAKGGPRMGPIFGGMIKSGIKIANLIKQTWRSS

>Dethiosulfatarculus sandiegensis
MMLDEVTITRAIIDRYMEKLHANLDLDVAIVGGGPSGLVAGYYLAKKGYNVAMFERKLSIGGGMWGGGMMMNEIVVQEEAKRILDEFGIPCREYVEG
YYTADSVVSTSTLTSKATLAGLSVFNLITVEDVMVRDNRVNGLVINWSPVEMAGLHVDPLTLRARYTIDATGHPAEVLNVISKKVDAKLSTDTGKVI
GERSLWAEVAESTTIENTKEAFPGVYTAGMCANAVFGAHRMGPVFGGMLLSGEKVAQVLDRKLQEED

>Dethiosulfovibrio peptidovorans DSM 11002
MELDERVISKAIVSRFFERLTDHLENDVVIVGGGPAGLVAGYVLADAGVKVSLFDRRLSLGGGMWGGGMLFNEIVVQSEGARILDDLGVSLREFEPG
YYTAGSVEAVSTLISSAVRAGVTVFNGMVAEDVVMREDRVIGLVINWSTVETSGLLVDPLAVRSDFIIDATGHDSNVTSTVEKKVPGRLLTETGKVE
GEKSLWCERAEKLTVDNTKEVYPGLFVAGMSANAVFGGPRMGPIFGGMLLSGEKAAKEILLRLNGKRVS

>Dissulfuribacter thermophilus
MREIDITKAIIDKHIEELNKCLCSDVVIVGAGPSGLVAGSILAQKGYTITIFEKRLAPGGGIWGGGMGFKYVIIQKEALDIVEEFNIPYEKYSDDLY
AVDAINFASGLILEAGKRGVHIFNLIAVEDLLVREGRVQGVVINNTFAKMNQFPIDPLTIEAKAVVDATGHEHEVVKTLSQKNDVTLNTPTGKPLGE
RSLFAETAEKAVVTNTKEVYPGLYVCGMATAAVYGGYRMGPIFGGMLMSGKKLAGLLEEALKA

>Elusimicrobia bacterium CG_4_10_14_0_8_um_filter_37_32
MKLDDIVISKAIMETFTKDFVDYLEVDVAIVGGGPAGLTAGYYLAKKGKKVVLFERKLSIGGGMWGGGMMYNKCVFQEDAKKILDEFGVTTHKYQEG
YYVTDSLETVSVLCSKAIKAGLKIFNLISVEDVMIRKEKITGLVLNWSAVQLAKLHVDPMTIRAKYVIDATGHDAEVVKIVVRKIGKKLYTKTGDML
GEKPMWAEVGEKDIIKNTKECYPGLYICGMASNAVFGGPRMGPIFGGMLLSGKRISGLVT

>Euryarchaeota archaeon ADurb.Bin165
MTLDEVTISRAIITDHLNTLVQYMEMDVAVIGAGPSGLVCATILAEKGLKVGLIEKKLSVGGGMWGGGMMFPRIVVQQGAKRLLDRFGIRSSEFSPG
YYTARSIEAVAKLAAAASDADVEFFNLTTVEDVMVKGDGLLSGLVINWQPVEATGLHVDPLTVRCRMTVDATGHDAIIAHYVSKKCGGLEIKGEGTM
WADNAEAAVVAHTKEVYPGLYVCGMAANAVMGGNRMGPVFGGMLLSGESAAEQILSRF

>Ferroglobus placidus (strain DSM 10642 / AEDII12DO)
MPFSEKNITRVIVREAAKEWEEISETDVVVVGAGPAGLTAAHYLADFGFDVVVFERRLSFGGGIGGGGMLFHKIVVEKEAKEIAEEFGIKTREVEDG
LYVIDAAEMLAKLSAGAIDSGAKVILGVTVDDVIYRPEPLRISGVLVQWSAVQIAGLHVDPLMIESKAVVDATGHDAEVVSVAARKIPELEIYVAGE
KSAYSELSEKLVVEKTGKVVDGLYVAGMAVSAVYGLPRMGPIFGGMLLSGRKVAEQIMFDLKK

>Fervidicoccus fontis
MSENLEFKITKLILEHSMKDLIEFADSDVIIVGAGPSGMTAAKYLADKKLKVLVLERKLSFGGGIGGGGNLMHKIVIKSDALKIIKDFEIEYKKTEF
EDLYTLDASELISKLATGAINSGAKILFGYSVEDLIVREKPLRVSGVVVKWSAIDLAQLHVDPIFFTGKAILDATGHDAELIKILAKKNPSFAINVK
NESSAHAELGEKQVVEFSGKVCDGLYAAGMSVATLHGLYRMGPIFSGMLISGKKVAELISKELGK

>Fervidobacterium changbaicum
MGKDLTISKLIVENFFEKLSNALEVDVAIAGCGPSALTLSLELSKKGYKVAIFEAKNEPGGGIWGGGMMFNEVVLESELEGYLKELGIRFKKFDEFI
VTDSVHLASALLYHTTLAGTMIFNNVFVEDLVMYDRRVSGVVINWTPTLREKLHVDPISIVSKFTVDGTGHPANLVKLLSKRGIISSIGGSTEASYN
FGIVGYEFPMDAENGERFVVENTREIYPGLYIVGMAAVSVGAGPRMGPIFGGMIMSGLRAAELISNELRKMGGSDDER

>Gemmatimonas sp. SG8_17
MRGRRGGGVTDFSEGQITRAIITAYHEKLWGQVVGDVVVVGAGPSGLVAATDLARRGLKVTVLEKRLSPGGGIWGGAMAMNEVVVQDAALPLLAEFS
VSFRSVKGGLHVINAVELASALSLKAVQTGAVILNLTVAEDVCVHRGRVTGVVANRTNLAEALPVDPVSFEAKAVLDATGHDAALVQMLQRRGLLKG
LTEMQGEGPMDAAGGESFVVDKVTEVYPGLWVSGMAVVATLGGPRMGPIFGGMLLSGKRAADLISDTLSGE

>Geoglobus acetivorans
MSYSERNITRIIVREAAKDWEDISDTDVVIVGAGPAGLTAAYYLREFGFDVVVFERRLSFGGGIGGGGMLFHKIVIEEEAKEIAEGFGMKLKEVESG
LYSVDSSDFLAKLSYSAVESGAKVLLGVTVDDVVFRPDPLRISGVLVQWSAVQISGLHVDPLMIESRAVVDATGHDAEVISIAARKIPELEIFIHGE
KSAYSEMSEKLVVEKTGKVADGLYAAGMAVAAVHGLPRMGPIFGGMLMSGKKVAEQIMFDLKK

>groundwater metagenome
MIDETKITELIVRSAVDDFLGNLKVDVVVVGGGPAGLTTARYLAKAKKRVVLFERKLSIGGGMWGGGMMFPRVVLQKGGKEILEECNVRYKKFDDLW
VADSIECVTKMTAKAIDEGVKIFNLISIEDVIIRSVQSNKNNKEGKTKICGVVLNWTAVQMANLHVDPLSVKSDFVVDATGHEASICHLVVKKVGNL
NTKTGDVLGERSMWAEKGESDIMNNTKEVYPGLFVSGMAANAVYGSERMGAIFGGMLMSGKKVSELILEKEK

>Hadesarchaea archaeon DG-33-1
MGGIEDTEITAAILKRFMRDFEDVTNLDVAIAGAGPSGITAASFLASGGAKVAVFERNLHVGGGMWGGGILFSRVVIQEAAKVMLEEVGVKLKPTAA
GYYTADSVEAVTKSTTAAVDAGARVMVGLTVEDVMIREKDRVAGIAVNWKAVELAGLHVDPVGISAKIVIDATGHDAMIARIVQRKVPNAKFPTSTG
GVVGEKPVWAEVGETEIVNNTREIYPGLIVTGMAANTVFGSPRMGPIFGGMLLSGRRAAEVALKV

>hydrocarbon metagenome
MWYLVELDERVISRAIIAVQMEKMLRYTDMDVAIVGGGPAGLTAASFLGAEGFSVALIEKKLSVGGGMWGGGMMFPRIVVQEEGRQLLDHFAIRYTR
YEEGYYVASSVEAVAKLTAAACDAGVEFFTLVTVEDVMVRSDKRLSGLVITWSPVEMAGLHVDPLTLGCRYTIDATGHDAVIARLVARKSGAVTVKG
EGFMWADRAESRITSHTREVFPGLIVAGMAANAVAGENRMGPVFGGMLLSGRHAAALVSRELASPKP

>Hyperthermus butylicus (strain DSM 5456 / JCM 9403 / PLM1-5)
MVNAVQAPEHSWLPHNVTSLREGALAALIIRKTAEKLTSITSVDVAIAGAGPAGLTAAWLLAEKGLRVVVVEHSLGVGGGMRGGSMLMPVGLVEDGL
PAELLRRAGARLDRVADGLYAVDPTEAVVKLAAKAIDAGAVILPGLHVEDLILWRSGSGYRVAGLVINLSPVVEAGWHVDPIYIEARATIDATGHDA
ELVKLLSKALGDSSIRVRGTRGMDVWEGEKLVVEYTGEVYPGLYAAGMAVSETYQLPRMGPVFGGMLASGARVAELVASRLSEQ

>Ignicoccus islandicus DSM 13165
MIDEGKVTSIIIEESSKELSQMAKGVDVVIVGAGPAGLTASHYLAKAGLKVLILERRVSLGGGISGGGSLFHKVVVEDVELEGYNPKEIAEELGVPL
KKVDDNLYTTDAAALVAKLSNASVSAGAKIVLGMHVEDLIYRIEEGVTKVKGVVALWSPIYLSGLHVDPIFFKAKAVVDATGHDAEILKIASKKLPN
VNFEVGREYGAWIDEAEKLVVKYTGKVLEGLYAAGMSVASFYRLPRMGPVFGGMLASGKKVAEKIIGDLEVS

>Ignisphaera aggregans (strain DSM 17230 / JCM 13409 / AQ1.S1)
MKELELRISRAILRNSVRELIEYSDVDVVIVGAGPSGLTAARYLAMNGFRVVVLERRLSFGGGIGGGGMLFHKIVVSSEALPILNDFDIKYYRDDEE
DLYMIDSSELMAKLAVGAINAGAKIFHGIHVEDVIYRENPLRITGVVIQWSAVVMSGLHVDPLFITSRAVVDATGHDAEVLQIVSRKIPEVGISLPG
ESSAYSELSEKIVVEKTGMVIPGLYVAGMAVAALYKLPRMGPIFSSMLLSGKRVAEIIANDLKKK

>Korarchaeum cryptofilum (strain OPF8)
MESLESRISKAIWESTYKDWLDIIDSDVVIVGAGPSGLTAASYLAKSGFKTTVIERRLSFGGGIGGGGMQLHKVVVDGRALKVLEDFKVRYSYLEKY
DLYVLDSAELMAKLASGAIDSGAKLIHGLTVEDLIVREDPFRVEGVVVQWSSVLLAGLHVDPLFIHSRVVVDATGHDAEVIRILERKNPSLGIKVPG
ERSAYSELSELSVVERTGKVVEGLYVTGMAVAALNQLHRMGPIFSGMLLSGRKVAEEIIRDLS

>Labilibaculum filiforme
MEQIVSAGIVDSYFKKLKENLSVDVAIVGGGPSGLVASYYLAKKGFKVALYESKLAPGGGMWGGAMMFNEIIVQKDALHILNELGVSYQHYQEDYYT
LDSVHATSALIYHATQAGVKIFNCSFIEDVVFQNDKVCGVVLNWSPVRREGLHVDPLVVMAKAVVDGTGHDCDIARTLERKNDVKLNTKTGKVMGEC
SLSIDEAERTTVENTKEIYPGLYVSGMASNGVSGGFRMGPIFGGMLRSGEKLAGLIAENLSK

>Latescibacteria bacterium DG_63
MKLDDVEISKAIIESFYAKLLDSLMCDVAVVGGGPAGLTAAYYLAKHGRKVVLFERKLSIGGGMWGGGIMFNEIVVQRDGKKILDEFGVRTTLVKEG
YFCADSVEAVSTICSKASQAGARIFNLFSVEDVMMTEERVTGLVINWSAVELSNLHVDPISIKAEHVIDATGHAAEVAHIIQTKSGSKLLTPTGTVI
GERPMCAEVAEKSILENTKEIFPGVLAAGMCCNAVFGAPRMGPIFGGMLMSGKKAAELIIDKPAAPKRRCLPDDE

>Lentisphaerae bacterium ADurb.Bin082
MAMENIITTAIIRQFADKLSAGTDLDVAFVGGGPSALVAAAKLAKKGLKTAIFEKSLAPGGGVWGGGMLFNEIVVQENVLGILEQIGISYQAVPDAK
GYYTVDSVEMASGLIFNAVKAGAKIFNAMSVEDIVFKEGRVNGLVINWAPVRKLAMPVDPLTVIAKAVVDATGHPCEIIRIACEKAQVKIATETGGV
LGERPMWVQHGEQQTVDSTAEYYPGLFACGMSATNVTGGYRMGPIFGGMILSGLKAADLIAKSLASLEKR

>Metallosphaera yellowstonensis MK1
MEIRQVDEVKITKYILKATFEDWMDIAENDVVIVGAGPSGLSAAYYLAKKGLKTTVFERRLSFGGGIGGGAMLFHKIVIESPADQVLREMNIRLQRV
EEGVYIVDSSEFMAKLASSAIDAGAKIVHGVTVDDVIFRENPLRVTGVAVEWTATQMASLHVDPLFIHAKAVVDATGHDAEVISVAARKIPELGIAI
PGEKSAYSEVAEKLTVDNTGEVAPGLYAAGMAVTEVKGLPRMGPIFGAMVLSGKKVAEDIASTLLMKARNT

>Methanobacteriales archaeon HGW-Methanobacteriales-1
MELDDITISRAIVEEFMNDFMDYMDIDVAIGGGGPAGLTAGYYLAKAGLKVALYERKLSIGGGMWGGGMMFNKIVVQEEGKRILDEFGIQSKKYQEN
YYVSDSVEATSTLCSKATQAGLKIFNLMSIEDVMIRGDDISGLVLNWSSVEMGGLHVDPLSIRSKAVIDATGHPCEVVKVVQNKIGPKLNTPTGEII
GEKSMWAEVGEPAIMENTREVYPNLYVAGMAANAVYGAPRMGPVFGGMLLSGEKIANMLIEKLK

>Methanobacterium subterraneum
MKLDDIIVSKGIVAGYMEELLDYMEMDVAIGGGGPSGLTAGYYLAKAGLKVALFEKKLSMGGGMWGGGMMFNKIVVQEEGKRILDEMGIRNQEYEEG
YYLADSVESASTICSKACQAGLKVFNLMEIEDVMIKGEGVEGLVINWSPVEMAGLHVDPITVGARAVIDATGHPCEVVKVLERKMEAPLKTETGKIM
GEKSMWADVAEQNIMGNVGEIYPGMYVTGMAANAVHGSPRMGPIFGGMLLSGEKVAEMLIEKLK

>Methanobrevibacter woesei
MKKLDDITVSKAIIQEYMNDFLDYTDMDVAIGGGGPSGVTAGYYLAKAGYKVALFERKLSIGGGMWGGGMMFNKVVVQEEGKRILDEFGIKSKKFED
NYYTVDSIECTSTLCSKATQAGLKIFNLMSIEDLMVRENGINGIVLNWSSVEMSGLHIDPLTVRAKAVIDATGHPTEITKIVEQKMGANLKTETGKI
MGEKSMWADRAEGKILDNVTEVYPGLWVTGMAANAVHGSQRMGPIFGGMLLSGEYVAQKIIEKLENE

>Methanocalculus sp. 52_23
MQLDEVTISRAILETHAEISSRYLDLDIAIVGGGPSGLVCAALAAEDGRKVAVIEKKLSVGGGMWGGGMTFPRIVVQEEGKRLLDQFGIRSRVYKPG
YHVASSVESVAKLTAAACDAGAEFFNLTSVEDVVIKEDGRVSGLVITTSPVEMTGLHVDPLTLAAKVTVDATGHDAVVAHCVLRKGGDITIHGESFM
WAERAETNIINHTREIFPGLIACGMAANAVAGEARMGPVFGGMLLSGEHAAVLAREISERV

>Methanocella conradii (strain DSM 24694 / JCM 17849 / CGMCC 1.5162 / HZ254)
MELDETLISRAIIDDFLRTLSDYVSVDVGIVGGGPSGLVCATYLARAGVKVAVFERKLSVGGGMWGGGMMFPRIVVQQEATRILDDFGIRYREYRPG
YYIAGSIEAVGRLTSAAAGAGAEIFNLMSVEDVMIRENKEVVGLVINWSAVDIAGLHVDPLTVRTRVVVDATGHPAEVCRIVERKVSGGAFKVPGEQ
SMWADRGERALISTTKEVYPGLVVAGMAANAVAGGPRMGPIFGGMLLSGEIAARIVKEKLGVS

>Methanococcoides burtonii (strain DSM 6242 / NBRC 107633 / OCM 468 / ACE-M)
MKLDEVTISRAIIEEFSKVFLDYTDVDVALVGGGPANLVAAKYLAEAGLKTVIYEKKLAVGGGMWAGGMMFPRIVVQEDALHILDEFGISYHEYENG
YYVANSIESVGKLISGATSAGAEIFNLVNVEDVMIRENDEICGLVINWTAVEIGKLHVDPLAIRSKVVVDGTGHPAVVCSTVQRKVPGAKLGELGVV
GEKPMWADVGEKMLLDTTKEVYPNLYVAGMAANAVAGAPRMGPVFGGMLLSGKQVAELIIERLG

>Methanococcoides methylutens MM1
MKLDEVTISRAIIDEFSKVFLDYTEVDVALVGGGPANLVAAKYLAEAGLKTVIYEKKLSVGGGMWAGGMMFPRIVVQEEARHILDDFGIDYHEYEEG
YYIANSVESVGKLISAATSAGTEIFNLVNVEDVMIRDNDEVCGLVINWTAVEIGRLHVDPLAIRAKVVVDGTGHEAAVCNTVQRKVPGAKLGELGVV
GEKPMWADVGERMLVETTREVYPNLYVDGMAANAVAGAPRMGPVFGGMLISGKQVADLIIERLK

>Methanococcoides vulcani
MKLDEVTISRAIIDEFSKVFLDYTEVDVALVGGGPANLVAAKYLAEAGLKTVIYEKKLSIGGGMWAGGMMFPRIVVQEEARHILDDFDITYHEYEKG
YYIANSVESVGKLISGATTAGTEIFNLVNVEDVMIRENDEVCGLVINWTAVEIGRLHVDPLAIRAKVVVDGTGHEAAVCNTVQRKVPGAKLGDLGVV
GEKPMWADVGERMLLETTKEVYPNLYVDGMAANAVAGAPRMGPVFGGMLLSGKQVAELIIERLK

>Methanococcus maripaludis
MDGKLRADEVAVTKSILKSTFDMWMDLIDVDVVIVGAGPSGLTAAKYLAQNGVKTVVLERHLSFGGGTWGGGMGFPNIVVEKPADEILREAGIKLDE
VIGEPELFTADSVEVPAKLGVAAIDAGAKILTGIVVEDLILKEDKVSGVVIQSYSIEKAGLHVDPITISAKYVIDSTGHDSSVIHTLARKNKDLGIE
VPGEKSMWADKGENSLTRNTREVFPGLYVCGMAANAYHAGYRMGAIFGGMYLSGKKCAELILEKLENK

>Methanocorpusculum labreanum (strain ATCC 43576 / DSM 4855 / Z)
MDLEVTKAITESWFARLQENLCFDAAIVGTGPSGLIAAVKLADAGYKVSMFESKLAPGGGMWGGAMLFSSIAVQNEAVYLLDELEIPYKRYNENLVV
CDSVLATSALIYQASKRGVVIHNGMSVEDVVFMDNRVSGVVVNWGPVVREGLHVDPLSFRAKIVVDATGHPCMISETAARKNNITLNTPTGKVCGEC
SLNAVEGEAMTVENTKEIYPGLYVCGMAANGVFGSPRMGPIFGGMLLSGEKVAKLIIEELK

>Methanoculleus thermophilus
MTLNEVTISRAILEESHRALIEHLEMDVAVVGGGPSGLACAALLGEKGLSCALIEKKLSIGGGMWGGGMMFPRIVVQEEARRLLDRFGIAYKEFEPG
YYVAKSVEAVAKLTAAACDAGVEFFNLTTVEDVMIRGDGRVGGLVINWTPVDMAGLHVDPLTVACTCTVDASGHDAVVARMIERKGGRLQVKGESFM
WAERAESRILDHTKEVFPGLFVAGMAANAVAGECRMGPIFGGMLLSGERAAELVAESLER

>Methanohalobium evestigatum (strain ATCC BAA-1072 / DSM 3721 / NBRC 107634 / OCM 161 / Z-7303)
MELDDITITKAIVDDFSKTFIDYTEVDVALVGGGPANMIAATRLAQEGYKVALFEKKLALGGGMWGGGMMFPRIVVQDEARKILEEFDINHYEYDNE
KGYYIANSIESVSRLINKTVTSGVQVFNLVNFEDVMIREDDRVTGIVINWTAVSIANLHVDPLTIRAKVVIDGTGHEAVVCNTVQRKIPNAKFEGVV
GERPMWADAGEKSLKETTREVYPGLIVTGMAANAVAGAPRMGPVFGGMLLSGEMAAKIAMSKLD

>Methanohalophilus euhalobius
MELDERIITRAIVEEFTNVFLDYTDVDVALVGGGPANLVAARYLAEAGLKTVLFEKKLSVGGGMWGGGMMFPRIVVQEEARRILDDFDVPYHEYEEG
YYVANSVGTVGKLISAAVSAGVEIFNLVSFEDVMIRDNDEVCGLVINWTAVEIARLHVDPLTIRSRLVLDGTGHEATVCNTVQRKIPGAFGGKEVVG
EKPMWADTGERLVMKNTREVYPGLIVTGMAANAVAGSPRMGPVFGGMLLSGEKAAQLAISRLKD

>Methanolacinia petrolearia (strain DSM 11571 / OCM 486 / SEBR 4847)
MKLDEVTISRAILSEQHKIMTEYLDIDCAVVGGGPSGITCAAILAQNGVKVALIEKKLSIGGGMWGGGMMFPRIVVQEEARRLLDHFGIKYTEYEKG
YYVASSVEAVSKSLAAACDAGAEVFNLTTVEDVVVKEDGGVSGLVINWTAVEMAGLHIDPLTMRTKVTVDATGHDSMIAHMVRKKGGALEIKGEGFM
WAERAETNILSHTKEVFPGLIVAGMAANAVGGETRMGPIFGGMLLSGEKAANMIIERLKK

>Methanolinea sp. SDB
MELSETTITRAIVSSQMKILLEYSELDVAVVGAGPSGLTAAAILGDAGYKVGVIEKKLSVGGGMWGGGMMFPRIVVQEPARRLLDRFEISYQPFEEG
YYVASSIEAVARLTSAACRGGAEFFNLTSVEDVMVKDDGRVSGLVINWTPVEMAGLHVDPLTIGCRYTIDATGHDAVVATLVERKGRNLEVKGEGFM
WADRAESEIISHTREVYPGLIVTGMAANAVAGEHRMGPVFGGMLLSGEFAASLVREKLNR

>Methanolobus profundi
MELDETIITRAIVEEYSKVFLDYIEVDVALVGGGPANLVAAKYLGEAGLKTVLFEKKLSIGGGMWGGGMMFPRIVVQEDAKHILDDFNINYHEYEKG
YYVASSIESVGKLICGATDAGAEIFNLIDVEDVMIRENDTVCGLVINWGPVSMNRLHVDPLAIRAKVVIDGTGHDAGICSTVQRKIPGTDIKLDVVG
EKPMWADVGEKILMDTTKEVYPGLIVTGMAANAVAGAPRMGPVFGGMMLSGKKAAELAIEKLRK

>Methanomassiliicoccales archaeon PtaB.Bin215
MEIDEVLVTRKIVERYTEEFLENVDVDVVIAGAGPSSLTAARYLAKAGLRVVIFERKLTPGGGMWGGGMTFPIIVVQEGSKDLLGEIGVRLRDAGDG
YFTADSVEASAKLISAAVTAGARLYNTISVEDVMIRQDSICGVVINSSAVEVAGLHVDPLAVRSKYVIDGTGHPAEVVHVVQRKVGRLNTPTGQIEG
EKSMWAEVGEQMTVENTVEVYNNLYVCGMASNAVMGAPRMGPIFGGMLLSGRKVAEMIIAREKKKGKKK

>Methanonatronarchaeum thermophilum
MNVDDKFVSKAIIDEFSKDFLDSLSVDVAIGGAGPAGMVAAKYLAENDIKTAVFERKLSVGGGMWGGGMMFPRIVIKEKSLPILDDLNINYREYQDG
YYIANSIESVGQTAAEAVKAGAEIYNLMTVEDLHYKENKVNGVVINWSSVDLAGLHVDPLTIESKITIDATGHDCELVKVAQERINKKLNTKTGKIM
GEKSMWAEQGEKDVVELTGEVLPGLYVTGMAVNAVHGKPRMGPIFEGMLLSGKKVAEQCIKKLK

>Methanoplanus limicola DSM 2279
MKTKVYESENKMKLDEVAISRAIVSEQSKVMLDYYDLDCAIVGAGPSGLTCAAMLGEEGLKVGVIEKKLSVGGGMWGGGMTFPRIVVQEEARRLLDH
FGIKYREYESGYFVSSSVEAVAKITSAACDAGAEFFNLTYVEDVVIKGDNRISGLVINQTPIQMTGLHIDPLTLATKVTIDATGHDSVVAHLVRDKG
GSVEIKGEGFMWADRAESNILSHTKEIFPGLIVTGMAANAVGGETRMGPVFGGMLLSGEKAAKLAKSALKK

>Methanopyrus kandleri (strain AV19 / DSM 6324 / JCM 9639 / NBRC 100938)
MEREITPIVLREGYEFINDCSESDVIVVGAGPAGLTCAYELAKSDVDVTIVERKLYVGGGMTGGGMLFPAGVIMEETAEVLEEVGVELRPAEAGLLA
FNPVEAAIKLANAALEAGARILVGIEVEDVIERRGRVCGVVVNWTAVKAANMHVDPLALEAEYTVDATGHEAAVCKLAGIEVKGEGPMWAERGEELV
VKHTQEVKPGLFVAGMAASAVKGAYRMGPIFGGMLESGKKAAEEILERLTE

>Methanoregula formicica (strain DSM 22288 / NBRC 105244 / SMSP)
MELDELTISRAILASQTNVLINHLELDAAVVGGGPAGLTCAALIAGQGKKVGVIEKKLSVGGGMWGGGMMFPRIVVQEEARRLLDLFGIRYTPFESG
YYVARSVEAVSKLTAAACDAGVEFFNLMSVEDVMIKADKRISGLVINWTAVEMGKLHVDPLVMGSRYTVDATGHDAVVARLVEKKGGDIRVKGEGFM
WADRAETNILNHTKEIFPGLVVAGMAANAVAGESRMGPVFGGMFLSGERAAQIVLREMKA

>Methanoregulaceae archaeon PtaB.Bin009
MELDEITISRAILSSQVEKLLEFMEMDVAVVGGGPSGLTAAALIGEQGFRVGLIEKKLSVGGGMWGGGMMFPRIVVQEEAKRLLDQFDIAHTSYTEG
YYVASSVEAVSKLTASACDAGVEFFNLFSVEDVMIRGDSRLSGLVVNWTPVEMAGLHVDPLTMGCRVAVDATGHDAVLARLVERKGGDVKVRGEGFM
WADRAESEIVSHTREVFPGLVVCGMAANAVAGEHRMGPVFGGMLLSGERAAALATSSLRQENSAA

>Methanosaeta harundinacea
MALDEVTITKAIVESYMESFLKYTDVDVALVGAGPANLVAAKKLAEADAKTVVFERNLSVGGGIWGGGMMFPRIVVQKEGCRILDEFGVWYREYEEG
YYIASSIETVAKLTAGVIDAGAEIINLVTVEDVMIREDERIAGLVINWEAVERTRLHVDPLSVRARVVIDGTGHDANICKVVQRKIPGAKVGSLGVV
GEKPMWADVGEKTVVEVTQEVYPGLIATGMAAAAVAGGPRMGPIFGGMLLSGEKAAMLALEKLGL

>Methanosalsum zhilinae (strain DSM 4017 / NBRC 107636 / OCM 62 / WeN5)
MELDEVVITRAIVDEFLNVFLDYTDVDVALAGGGPANLVAAKYLAEAGYKTVLFEKKLSIGGGMWGGGMMFPRIVVQEEARRILDDFNITYKEYEDG
YYVANSIESVSKLAAGATSAGAEIFNLVSVEDVMIRENDRVSGLVINWTAVGIGKLHVDPLTIRSKVVIDGTGHDASVCNIVQQKVPGAQLGELGVV
GEKPMWADVGEKLLMETTREIYPGLIVSGMAANAAAGAPRMGPVFGGMLLSGEKAAELAISKLD

>Methanosarcina acetivorans (strain ATCC 35395 / DSM 2834 / JCM 12185 / C2A)
MELDEVIITRAIFDEYSKTFLDYTDIDVALVGGGPANLVAAKYLAEAGVKVALYEQKLSLGGGMWAGGMMFPRIVVQEEATRILDDFGIRYKEYESG
YYVANSVESVGKLIAGATSAGAEVFNLVSFEDIMIRENDRVTGIVINWGPVTTQRLHVDPLMIRTKLVIDGTGHEAVVCNTILRKIPNAKIGELGLL
GEKPMWSEVGERLAVNATQEIYPGLIVAGMAANAATRAPRMGPVFGGMLLSGEKAAKLALDRLKTI

>Methanospirillum stamsii
MTLDEITISRAIISDYMHTLLEYMEMDVAIVGGGPSGLVCSALIAEKGYKVGLIEKKLSIGGGMWGGGMMFPRIVVQSEAKRLLERFNITHSEFSPG
YYTARSIEAVSKLTTAAVDAGVEFFNLTTVEDVMVKGDGRLSGLVINWQPVEATGLHVDPLTIRCRMIVDATGHDAVIAHYVSKKMGKPDIKGEGTM
WADNAESAVVTHTKEVFPGLFVCGMAANAVSGGHRMGPVFGGMFLSGESAAVQILQQL

>Methanothermobacter defluvii
MEIHAGVKMKLDDIKISRAIVEGYMEDLLDYMEMDVAIGGGGPSGLTAGYYLARAGLKVALFERKLSIGGGMWGGGMMFNKIVVQDEGREILDEFGI
RSEPYDEGYHVADSVEATSTLCSRACQAGLKIFNLMSIEDVMIRDEGITGLVLNWSSVEMAGLHVDPLTVRAGAVIDATGHDCEIVKVVERKIGPEL
NTPDGRIQGERSMWADVGEAALIENTREVYPNLYVAGMASNAVYGAPRMGPIFGGMLVSGRRVAEMIIEKLK

>Methanothermococcus okinawensis (strain DSM 14208 / JCM 11175 / IH1)
MDKFKIEEKDVTTSILKATFNMWMDIVDVDVVIVGAGPSGLTAARYLAKEGVKVVVVERHLSFGGGTWGGGMGHPYITVQKPADEILREVGVKLEEI
DGGLYVADSVEVPAKLGVGAIDAGVKILTGVIVEDLILKENKVSGVVINSYAIDKAGLHIDPLTINAKYVIDATGHDASVTNTLARKNKDLGLEVPG
EKSLWAEKAENSILRHTREIFPGLFVCGMAANATHGGYRMGAIFGGMYLSGKKVAELILEKLKNND

>Methanothermus fervidus (strain ATCC 43054 / DSM 2088 / JCM 10308 / V24 S)
MVLNEVTISKAIISKYMEELIDNTNLDVAIAGGGPSGITAGYYLAKEGFKVALFEKRVSIGGGAWGGGMMFNKIVVQEEGKKILDEFDVNTERYENN
YYVADAIEMITTLASKACKSGLKIFNLINIEDIVIKNKKISGIVVNWTAAEMAKIHVDPLVIKSKFVIDATGHDCEVVKAVEKKLGPVLNTETGRIV
GEKPMWAEKGEKAVIKNTGEVYPNLYVAGMAANSVYGSYRMGPIFGGMLLSGKKVAELIRERLL

>Methanothrix soehngenii (strain ATCC 5969 / DSM 3671 / JCM 10134 / NBRC 103675 / OCM 69 / GP-6)
MSLDEVMVTKAIVEGYLESFLENTEVEAALVGAGPANLVAAKRLAEANIKTVLFEKRLSVGGGLWGGGMMFPRIVVQQAAIRILEEYGIRYHEHCKG
YYVANSIETVAKLTARAIDAGAQIVNLVTVEDVMIREQDRVVGLVINWTAAEMAQIHVDPLCIRARYVIDGTGHEASVCRVVARKIPGAIIGIDGVK
GEKPMWAEVGERTVVEMTQEVYPGLVVAGMAAAAVCGGPRMGPIFGGMLQSGEKAAGIVIENLNK

>Methanothrix thermoacetophila (strain DSM 6194 / JCM 14653 / NBRC 101360 / PT)
MALDEVKITRAIVESYLESFLKCTDVDVALVGAGPANLVAAKRLAEADVRVVLFEKRLSVGGGLWGGGMMFPRIVVQKEACRILDEYDIWYREFEEG
YYVADSIEVVAKLTAGAIDAGAELINLVSVEDVMIREGDRIVGLVINWTAADMAGIHVDPLAIRARVVIDGTGHDAAVCRVVQKKIPGAIVGESGVI
GEKPMWAALGEKIVVDATREVYPGLIVAGMAATTVAAGPRMGPIFGGMLLSGEKAASIALEKLAQSVD

>Methanotorris formicicus Mc-S-70
MDLRLKADEYTTTKAILKSAFNMWMDIIDVDVAIVGGGPSGLTAARYIAKKGYKVVVLERHLAFGGGTWGGGMGFPYIVVEEPADEILREVGIKLEK
VDGEEGLYTADSVEVPAKLAVGSIDAGAKILTGIVVEDLILRENRVAGVVINSYAIEKAGLHIDPITITAKYVVDATGHDASVATTLSRKNPELGLE
VPGEKSMWAEKGENALLRNTREVYPGLFVCGMAANATYGGNRMGAIFGGMYLSGKKCAEMVVEKLKNNE

>Nitrospira bacterium SM23_35
MELDEVVITKAIVDQFCKKLTNHLETDVAIVGGGPSGLVAGYFLAKAGRKTVLFERKLSVGGGMWGGGMLFNEIVVQKAAVRILKEFGITYCEFQKN
YYTADSVESISTLISRAVQAGVTIFNCITAEDVLMRTSRVTGLVLNWSAVEMARLHVDPLAVRSRFVVDATGHETAVVRLVQNKVPGTLKTLSGKVE
GEKSMWSDKAESLTLKNTREVFPGLYVAGMAANATFGGPRMGPIFGGMLLSGEKVAKLLLQALSQKSK

>Nitrospira japonica
MGKPKPAPLRERDITRQIAREYYKEFDQLIESDVIIVGAGPSGLICAHDLADMGFKTVIVEQNLALGGGFWHGGYLMNKATICAPAHKILDEIDVPC
KRIKDCEGMYIVDPPHATGALIAAAYRAGAKVLNLTRVVDLILRRDGSLDGVVVNNTTAEMAGHDLIHVDPIALESKIVVDATGHDAVVVSLLHKRS
LYTEVPGNGAMWVSRSEEDVMDHTGEVYPNCFVIGLAVSAVHGTPRMGPAFGSMLLSGRYGAELIKKKLKQE

>Nitrospiraceae bacterium
MNPINVAPLRERDITRHIAREYYKEFDSLIESDIIIVGGGPSGLLCARDLATSGFRTLLIEQSLALGGGFWSGGFLMNKATICEPADQILEELGIPF
KPIKDCPGMTMVDPPHVTSRLISAAYEAGVKIMNLTKVVDLILRQDHRIEGVVVNNSTVEMAGHDTIHVDPIALESQIVVDATGHDAVVVNLLHKRN
LYQKVPGNGAMWVARSEALVVENTREIYPNCFVAGLAVAAVDGSPRMGPAFGSMLLSGRYAAELVRQKLKGE

>Nitrospirae bacterium
MPKPPTPAPLRERDITRHIAREYYKEFDQLIESDVIIVGAGPSGLICAHDLAAMGFKTVVVEQSLSLGGGFWSGGYLMNKATICEPANEILEEIGVP
CKKITECAGMYMVDPPHATGALIAAAYRGGAKIMNLTKVVDLIIRRDGILEGVVVNSTTAEMAGHDAIHVDPIALESKIVVDATGHDAIVVELLHKR
NLHKAVPGNGAMWVAQSEQEIMDRTGEVYPNCFVIGLAVAAVYGTPRMGPAFGSMLLSGRYGADLIKKKLKGA

>Omnitrophica bacterium RBG_13_46_9
MDEALISRAITESFTKDFIDAFNVDVAIAGAGPSGLICAYYLAKQNVKVAVFERHLRVGGGMPGGGMMFNRIVVQEEAMPILKEFGVSAKRYKKDLY
IVDALEAISTFCSKTIKRGAKIFNLINVEDVVIRKDRIAGVVLNWSAVSWAKLHVDPMAVRSKAVVDATGHDSEIARIVERKTGPVLRTETGGVIGE
KSMWAEIGEKMILENTKEIYPGLIVCGMAANAVFGSPRMGAIFGGMLLSGKKAAEVARKVIKSK

>Omnitrophica WOR_2 bacterium RIFCSPHIGHO2_02_FULL_68_15
MFARAQEAQITRAIVRAFAKEFDGLVRSDVLIVGAGPSGLVAAMDLARRGRRVLVVEQTNYLGGGLWLGGYLFNKLTVRAPAHRLLKELKVPCRQVQ
PGLYVADAPHVCARLIAAACDAGVKFAQMTEMVDVVVREGGRVEGLVINWSPVSALPKGLAHVDPVALEAKVVVDATGHDAAVVRLLAKRGLAAPVP
GDGAMWVERGEQAVMDKTGEVHPGLFAAGLAVSAVHGTPRMGPAFGAMLLSGRRCAQMIERAYFA

>Peptococcaceae bacterium SCADC1_2_3
MPLEDITISKAIITRYNQELLAALESDVAIAGGGPSGLVAAFYLAQQGAQVVLFERNLSLGGGMWGGGMMFNQIVVQEEAIPILDTFGVRYRTFEPG
YYTAHATETVAAIILGAVRKGVKILNLISAEDVMVRNERVCGLVLNWTAVGLARLHVDPIAASCSCVIDCTGHDAQIANIVVRKMGAVLKTSSGKIE
GEKPMWAERGEAAIIKNTGEIYPGLYVAGMAANAVYGNHRMGPVFGGMLLSGKRAAELIKGER

>Phorcysia thermohydrogeniphila
MQNLNEVIISQAIIESFMEKLKNSLEVDVAIVGGGPSGLVAGYYLAREGFKVSIYERHLAIGGGMWAGGMLFNEIVVQEMGREVLDEFGVRYREFQP
GYYVADSVEAVTTIASKAVKAGAVIFNGVTAEDVVLKKVNDEYRVCGLVINWTSVERSRLPVDPLVITAKYVIDATGHDASVVSTLQKKAGIKLATE
TGCVIGEKPLWASVGEEDTVKNTREVFPGIFVSGMAANATCGSHRMGPVFGGMLVSGKKAAQEIAEKLKGNKEE

>Planctomycetes bacterium DG_20
MDDFDETDVSQAILRAYYAKVADALQGDVLVVGAGPSGLVAAWRLAQAGHRVVVLEKRLSPGGGIWGGSLGMNEVAVQKHALAILDEAGVRHQPSGR
LFTADAMELASALCLKALHAGAVILNLMTAQDVCVRSGRVTGVVANRSLLGESLPIDPIVFSARAAIDATGHEAVLANCIQRRGLLKNSLGRLPGEG
PLDAPAGERFVVDHVAELYPGLWTTGMSVCASLGGPRMGPIFGGMLLSGEKVAALVGQALKKTRPQVRHE

>Porphyromonas sp. CAG:1061
MEKLVSQGIITRYFEKLNDCLDLDVAIVGGGPSGIVAAYYLAKAGLKVAQFDRKLSPGGGMWGGAMMFNEIVIQEEALEIIKEMGINYEPYQDKLYT
MDSVESTAALLYNAVHAGARIFNCYSVEDVVYKENRVSGVVVNWTPVLREGMHVDPLNIMAKYVIDGTGHDSEICRVVAKKNGATLNTSTGGVVGEQ
SLDVITGEKMVVEGTKEIYPGLYVCGMASSAVYGTPRMGPIFGGMLMSGKKVANLIIDQLK

>Prevotella amnii
MIEKEISKGIITTYFEKMEKSLDLDVAIVGGGPSGIVAAYYIAKAGLKVALFDRKLSPGGGMWGGAMMFNQIVIQKEALDIIKEFEINYEQYSDNLF
TTDSIECTAAILYKAVHAGATIFNCYSVEDVVFKNNIVSGVVVNWTPVLREGLHVDPLNIMAKFVIDGTGHDSEICKVVARKNNITLNTSTGKVVGE
RSLDVIEGEQQVVEGSKEIYPGLYVCGMASSAVGGTPRMGPIFGGMLMSGKKVADMLIKRIQS

>Prevotella nigrescens
MIEKKISKGIISTYFAKMEKCLELDVAIVGGGPSGIAAAYYMAKAGLKVALFDRKLSPGGGMWGGAMMFNQLVVQEEALEIIKDFDINYEPYEDGLY
TADSVESTSALLYKATHAGATIFNCYSVEDVVFKNNIVSGVVVNWTPVLREGLHVDPLNIMAKFVVDGTGHDSEMCQVVARKNGIKLNTATGDVIGE
RSLDVAEGERQVVEGTKEIYPGLYVCGMASSAVGGTPRMGPIFGGMLLSGKKVAEAIIERLK

>Prevotella stercorea DSM 18206
MIETQVSKGIITTYFDKLQNNLDLDVAIVGGGPSGIVAAYYMAKAGLKVAQFDRKLSPGGGMWGGAMMFNQIVIQEEAMHIVKDFDINYQAFEDGLY
TIDSVESTSSLLYHAVHAGATIFNCYSVEDVVFKNNVVSGVVVNWTPVLREGLHVDPLNIMAKCVIDGTGHDSEMCKVVARKNGIQLDTATGGVIGE
RSLDVVEGERMVVEGTREVYPGLYVCGMASSAVAGTPRMGPIFGGMMLSGKKVADMIIEKLKK

>Prosthecochloris sp. ZM
MEEKISKFIIQSFFAKLEDSLTVDVAIVGAGPSGLIAAKELAKAGKKVAIFESKLAPGGGVWGGGMLFNEIVLQENIIPILDEYAIRYKTTGEGYVT
ADAVEVSSALIYGAVHAGVRIFNAVRVEDLAMRDERVCGVVINWNPVSRLEMHVDPLVITSRAVLDGTGHPSELINLASNKAGITLDTPTGKVMGEK
PMWMENGESSTVINTKRLYPGLYASGMAANNAMGGFRMGPIFGGMFLSGKKVAGLILEDIQG

>Pseudothermotoga lettingae (strain ATCC BAA-301 / DSM 14385 / NBRC 107922 / TMO)
MKDTMISTLIVNRYFKKLRSFLELDVAIVGAGPSGLTAAYELAKKGFKVAIFEEKNTPGGGIWGGGMMFNEIVLEKELEDFLNELGITYVIQENHVL
VDSVHFASALLYRTTMVGATVFNNISVEDVAMQDGKVCGVVINWGPTMRLGLHVDPITVKASFVIDGTGHPANVASLLAKRGLIEMKMELPMNADEA
EQFVVENTGEIFPGLMASGMAACAVHGGPRMGPIFGGMILSGKKIAQIIEEKLR

>Pyrobaculum aerophilum
MELKIGRAIIRHALKDLDEYSDVDVAIVGAGPAGLTAAKYLAEKGLKVVVYERRFSFGGGIGPGGNMLPKIVVQEEAVPILRDFKVRYKPAEDGLYT
VDPAELIAKLAAGAVDAGAKIILGVHVDDVIFRGDPPRVTGLLWIWTPIQMSGMHVDPLYTQAKAVIDATGHDAEVVSVAARKVPELGIQVVGEKSA
WSEVSEKLVVEHTGRVAPGLYVAGIAVCAVYGLPRMGPIFGGMLMSGKKVAEVVYKDLMAEAHAVRA

>Pyrococcus abyssi (strain GE5 / Orsay)
MLREVTISRAIIESYYRDLLNNLELDVAIVGAGPSGMVAAYYLAKGGAKVAIFEKKLSIGGGIWGGGMGFNKVVVQEEAREILDEFDIRYEEFEKGY
YVADAIEVATTIASKTVKAGVKIFNMIEVEDLVVKDNRVSGIVINWTPVLMTGLHVDPLTVEAKYVIDSTGHGAQVAQFLLKRGLIERIPGEGAMWA
EQGERLTVENTREVFPGLYVTGMAANAIAGAPRMGPIFGGMFLSGKKAAQEILEKLNL

>Rikenella microfusus
MEKLVSLGIVENYFEKLKNNLSVDAAIVGGGPSGLVAAYYLAKAGKRVVLYERKLAPGGGMWGGAMMFNDIIVQQEALPILDELGVCYKPYREGACV
VDSVHATSALVYAATKAGATIFNCYSVEDVIFRDEAVAGLVVNWAPVMREGMHVDPLMVTAKTVLEGTGHDCMIARLVARKNNVRLNTPTGEVAGER
SLNVEQGERLTVENTKEIYPGLFVSGMAANGVSGSFRMGPIFGGMLMSGKKAAELMIAKING

>Saccharolobus solfataricus (strain 98/2)
MEVKIKQVDEVKISRYIIKETMEDWYQFVESDVVIVGAGPSGLSAAYYLAKAGLKTLVFERRLSFGGGIGGGAMLFHKLIIEKPADEILREVNIRLK
EVEEGVYVVDSAEFMAKLATAAIDAGAKIIHGVTVDDVIFRENPLRVAGVAVEWTATQMASLHVDPIFISAKAVVDATGHDAEVISVAARKIPELGI
VIPGEKSAYSERAEELTVINTGKVAEGLYATGMAVTEVKGLPRMGPIFGAMVLSGKAVAGEITKDLLKSEIRA

>Smithella sp. SDB
MLNETTISRAILDAYFKKLDSCLELDVAIVGGGPSGLVAGYYLAKAGRKVALFERRLSIGGGIWGGGMMFNAIVVQEAGRQLLEEFDLKGSEYAPGY
YVLDAVDVTATLIHKAVRAGLQVFNLIAMEDVVIKNERVAGLVINWGAVDTLKWHVDPLTIHARYVIDGTGHPANVTEVLVRKMGVRLNTSTGGIVG
EKSMEAEQGELQTVENTREVYPGLYVSGMAANAVFGGYRMGPVFGGMLLSGRKAAEEILAHL

>Spirochaetes bacterium ADurb.Bin215
MLDETVISRAIIETYMKKLTDNLSVDVAIVGAGPSGFVAGYFLAKAGRKVVIFERALAVGGGMWGGGMGFNEIVVQEEGKAVLDEFDLPAVRYVQGY
YTLDSVRAVSALALRAVEAGVTVFNLVGVEDVVLHDERVSGLVLNWGTVMKAGIPVDPLTVHSRCVLDATGHPAHVAEVLCRKMGVSLNTPTGKMMG
EMSMDAEKGEKQTVENTREAYPGLFVSGMAANAVFGGYRMGPVFGGMLLSGKRAAEVMLARLGER

>Staphylothermus hellenicus (strain DSM 12710 / JCM 10830 / BK20S6-10-b1 / P8)
MKFFPQNLYELSEGDLSKTLIDALYKKLSEIVKVDVAIVGAGPSGLTAAWKLGEKGYKVLVLERMLGVGGGMRGGSMLLPVGLIEDGEAAEIAREAG
ARINKIRNGLFVVDPSELAVRLASKAIENGAIIWPGVLVEDLITRGRGEDLVVKGVLINWTPIYEAGWHVDPFYIEANAVVDATGHDGSLLRVLAKR
HPELKINIPGMSSQNVWIGEEMVVEKTSMVVKGLFVTGMSVAELYNTNRMGAIFGGMLVSGRKVADLIDDYFGKTRTLREQ

>Sulfolobales archaeon SCGC AB-777_J03
MASVRRVPESKISRFIVEETMKDWMDIVESDVVIVGAGPSGMTAAYYLAKAGLKTVVFERRLGFGGGIGGGAMQFHRLVIEEPADEVLREFGVRLKK
VDEGVYVVDAAEFMAKLASKAIDAGAKIILGVTVDDVIFREDPPRVAGVAVEWTATQMSGLHVDPLFISAKAVVDATGHDAEVISVAARKLPELNIS
VPGEKSAYSEVAEQLVVDNTGPVAPGLYAAGMAVCEVKSLPRMGPIFGAMVLSGKRVAELIIQDLRKS

>Sulfolobus acidocaldarius
MSDSIKIKAIDEVKISRYIIKQTMEDWMNFVENDVVIVGAGPAGMSAAYYLAKHGLKTLVFERRLSFGGGIGGGAMLFHKLVIESPADEVLKEMNIR
LEKVEDGVYIVDSAEFMAKLAASAIDAGAKIIHGVTVDDVIFRENPLRVAGVAVEWTATQMAGLHVDPVFISAKAVVDATGHDAEVVAVASRKIPEL
GIVIPGERSAYSEMAEKLTVEQTGVVAPGLYVAGMSVTEVRGLPRMGPIFGSMVLSGKKVAEDIIKDLRNS

>Synergistaceae bacterium
MRLDEVTITKAIMERYFDKFMNNLELDVAIVGGGPSGLVAGYFLAKAGHRVALYERKLSVGGGMWGGGMLFNEIVVQEDAKRLLDELDVPTLPYKEA
GYYTADSVETVSTITSKAVKAGLVVFNCISVEDVVVKDDRISGLVINWTAVPMANLHVDPLSIRSRYVIDATGHDTEVVAMVAKKAPGRLLTPSKNI
EGEKFMNPEEAERLTLENTKEVFPGLYVAGMACNATFGGPRMGPIFGGMLLSGEKVARLILKELSK

>Syntrophaceae bacterium PtaB.Bin038
MALNEVTISRAIVETYTKKLLAHLDVDVAVVGGGPAGLVAAYFLAGAGRKVALYERKLSIGGGMWGGGMMFNEIVVQAEAKGILDHFGVRTQEYAPG
YYTADAIEAVTTICSRAIQAGAKVFNCITVEDVVIRDNRVMGLVITWSPVEMTGLHVDPLTIHAKAVIDATGHDTEVLHVIER
KADVTLNTPTGKLMGERSMWSEKAERLTIDNTREICPGVYVAGMSANAAFGGPRMGPIFGGMLLSGRKVAEQILAGG

>Syntrophobacter sp. DG_60
MALDELIITQAIVERFSEKLKGCLMLDVAIVGAGPSGLVAGYYLAKNGHRVAIFEKKLSIGGGMWGGGMMFNQIVVQTEGKRILDEFEIKTAPFSEG
YYTADAIEAITTICSKACQAGVNVFNTISVEDVLVREGRVIGLVINWSTVEMAKLDVDPLTIRAQYVIEATGHATEVVKVIEKKMGESLLTPTGKII
GEKSLWAEIGEADTIKNTKEAYPGLFVCGMAANATFGSYRMGPVFGGMLLSGERVAKLIHERLRTKT

>Syntrophobacteraceae bacterium
MELNEITITQAIVDRFLEKFRNSLETDVAIVGGGPAGLVAGYFLAKAGCKVSLYERKLSVGGGMWGGGMLFNEIVVQEEAKRLLDLFGVGSHHYRDN
YYTADSVEAISTLTSQAVKAGVMIFNCISVEDVVMRPERVIGLVLNWTAVEMAGLHVDPLAIRAKFVVDATGHDVEVVRVVKRKVPGKLLTPSGEIE
GEKSMWSEVAEKLTLDNTREVFPGLYVAGMAANATFGGPRMGPIFGGMLLSGEKVAQTLIDQLKK

>Syntrophobacterales bacterium CG_4_8_14_3_um_filter_49_14
MELNEITITKAIIERFSEKLIACTEVDTAIVGGGPAGLVAAYFLAKVGKKVAIFEKKLSIGGGMWGGGMMFNEIIVQPEARELLDLFDVRTRKYEAG
YYSADAIEAVSTICSYATKAGARVFNCITVEDVMIREGRVIGLVINWTPVSMTGLHVDPLTIAAKSTIDATGHATEVLRVIERKTDIRLNTPSGALM
GERSMWADRAERLTLENTREICPGVYVAGMSANAAFGGPRMGPIFGGMLLSGKKVAELIIANG

>Syntrophus sp. (in: Bacteria)
MELNEITITRAIIERFTEKFLACTEVDVAIVGGGPAGLVASYYLAKAGRKVALFERKLSIGGGMWGGGMMFNEIVVQEEAKEILDLFGITSRPYETD
YHTADAIEAVSTICSYAVKAGAKIFNCMSVEDVMIREGRVTGLVITWSPVEMAGLHVDPMTIGAKWIIDSTGHATEVLRVIERKADVRLFTETGKLM
GERSMWAEKAERMTLENTKEICPGVYVAGMSANAAFGGPRMGPIFGGMLLSGRKVAEQILSR

>Thermanaerovibrio acidaminovorans (strain ATCC 49978 / DSM 6589 / Su883)
MELDERRISAVIVRRFMDRLLDSMDLDVAIVGGGPAGLVAGHNLAREGFKVAMFERKLSLGGGMWGGGMMFNQIVVQEEGAQVLREFGVRVLDEGEG
YYSADSVEAVSTLISSATRAGLRVFNCVTAEDVTMREDRVVGLVITWTPVEMAGLHVDPLAIRSRFVIDATGHDINVVRVVERKVPGKLMTPTGRAE
GEKSLWSHRAEELTLENTREVFPGLYVAGMSANATFGGPRMGPIFGGMLLSGRKAAQLVSRALRGQGGRG

>Thermincola ferriacetica
MHLDETVISRGIVQKYMEELMDYMNTEVAIVGGGPSGMVAAYYLVKRGCKVALFDRKLAVGGGMWGGAMMFNKIVVQSAGKRILDEFAISCEEYERG
YYVADAVESVTTIASMTVKAGCKIFNLIGAEDVMVEDGRVTGLVLNWTPVQVNNYHVDPLVVRAKYVIDGTGHPAEVTQTLTRKMGVRLNTPTGGVA
GEKPMNALKGELDVVENTREVFPGLYVTGMAANAAFGSHRMGPVFGGMLLSGEKAAMEIAARLGK

>Thermococcales archaeon 44_46
MLRDVTISRAIIETYFKELLEHLNLDVAIVGAGPSGMVAAYYLAKGGAKVAIFEKKLSIGGGIWGGGMGFNKIVVEEEAKEILEEFGVRHEEFEEGY
YVADAIEVATTIASKNIKAGAKIFNMVEVEDLVVKENRVAGIVINWTPVKMTDLHVDPLTVEAKFVIDSTGHGAQVTQLLEKRGLIERVPGESAMWA
EMGEKLTVEHTKEIYPGLYVTGMAANAVAGAPRMGPIFGGMFLSGRKAAFEILEKLKK

>Thermococcus celer Vu 13 = JCM 8558
MLKDVEVSRAIIEAYTKDILDSLKLDVAVVGAGPSGMVAAYYLARGGAKVAIFEKKLSVGGGIWGGAMGFNRIVVEESAREILDEFGVDYEEFKPGL
YVADAIEVATTMASKTVKAGVKVFNMVEVEDLVVKDGRVAGVVVNWTPVKMTGLHVDPLTVEAEFVIDSTGHGAQITGHLLKRGLIEELPGEGPMWA
EMGERLTVEHTKEVFPGLYVTGMAANAVAGAPRMGPIFGGMFLSGRKAALDILERLGV

>Thermodesulfatator autotrophicus
MALDEIKISRAIIERYFKKLTDYLEMDVAVVGAGPSGLMAAYKLASEGFKVAVFERRLSIGGGIWGGGMMFNEIVVQEEGARLLKEIGVRTEPWNGG
EYYTADAVEVACILAAKSVQAGAKIFNLIMVEDVMVRDNRVVGLVLNWSATEIAGLHVDPLAVKAKYVVEATGHETAVLQVMQKKLGAKLNTETGKV
MGEKSMWAEVAENLTVDYTREVYPGVFVAGMAANATFGAYRMGPIFGGMLLSGERVAQLIAERLRQ

>Thermodesulfobacterium geofontis (strain OPF15)
MELKIQRAIVKFGMEDLYEYSDVDVLIVGAGPSGLTSAKYLADKGFKVLVYEKRLSFGGGIGGGGNMIPKIVVQEEALPILKDFKIKYKEAEKNLYT
IDPAELIAKLAVGALDAGAKIILGVHVEDVIVRDNPPRVTGVLWRWTAIEISGLHVDPLYTQSKALIDATGHGAEIVQIAAEKNPELNIIIKGEKSN
WSEVSEKLVVDYTGKVAEGLYVTGIAVCEVFGLPRMGPIFGGMLMSGKKIAEIIEKDLRG

>Thermodesulfobium acidiphilum
MTKHFLNPVTDVNVSKLILKHYFESITDALTSDVIIVGGGPSGLTAARELGNSGYKVVIMERKLSPGGGTWGGSMSFNKVVIQKDLKDYLNELEIPF
VEDLDALVVDSCLFASQLIAKALKTQNVKLFNLMTVVDLEYTNNAITGVVVNNTGIEIAGLHVDPMVFQTKAVLDATGHDAIAANIYSKRVQLPLRK
EHFMNAVQGEEDTVNNTKMLANGLFVSGMAANNVDGGSRMGPIFGGMIKSGLKAAKLIMEYIKTV

>Thermodesulforhabdus norvegica
MAELNEIIITRAIIDRYHAKITGNLDVDVAIVGAGPSGLVAGYHLAQKGYRVTIFERKLSVGGGIWGGGMLFNEIVVQDEARRILEEFGVRVNRYEE
NYYTADAIETVSLLAARAIQKGVTILNGITVEDVVMRPNRVIGLVILWSAVEIAGLHVDPLAIRAKYIVDATGHDTEVVKVVHKKVPGRLMTPTGNI
EGEKSMWSEEAEKLTLENTREVFPGLFVAGMAANATFGGPRMGPIFGGMLLSGEKVAHLIDERLKQGS

>Thermoplasmata archaeon HGW-Thermoplasmata-1
MTVIDEIVVTRKIFDRFSREFLDHLDVDVALVGGGPANMVAAHYLAKAGKKVVLFERKLAPGGGMWGGGMNFPVIVIQEAALPIMNEFGIKVEGSND
GYYTADSVECVAKLLAKSIDSGARVFNSMTVQDVMIRDDDNKVSGVVINWTGVDITKMHVDPLTIRAKFVVDGTGHPCEVCNVVAKKAGRLRTPSGK
VEGERSMWAEVGEETTVNNTVEIYPGLYVAGMAANGVMGAPRMGPIFGGMLLSGKKVAEMILQKLGA

>Thermoprotei archaeon ex4572_64
MSIVRKVSEDEISKAIINEALKELESLVDVDAVVVGSGPSGLTCSYYLAKYGLKTVLIERRLSFGGGIGGGGMLLPSIAIESPAAELIHDEFLVNIK
KVRDGLYVMNPAEFIAKLASKAINAGVKVLLGVSVEDVIFRSNPLRIAGVVINWSAVHISQLWVDPLFIKAKAVIDATGHDAEVVNIVSKKIPDFKL
AIKGEKSACSIEAEDLIISYSGKVVEGLYVTGMATAKVYGLPRMGPIFGGMVLSGKKTAEEVYRDLSRELNR

>Thermoproteus uzoniensis (strain 768-20)
MRKINLIWKLRGAMELKIGRAIIRHGAEDLYEYSDVDVAIVGAGPSGLTAARYLAEKGLKVIVLERRFSFGGGIGPGGNMYPKIIVQEEALPILRDF
KVRYKPAGDGLYAVDPAELIAKLAAGAIDAGAKILLGVHVDDVIFRGDPPRITGLLWIWTPIQMSGMHVDPLYIQTKAVVDATGHDAEVVSVAARKV
PELGIQLQGEKSAWSEVSEKLVVEHTGKVAPGLYVAGMAVAAVFGLPRMGPIFGGMLMSGKKVAEIVAKDLAAEVHAV

>Thermosipho africanus
MWDYEVSKIIVERFFEKLNDNLNVDVAIVGGGPSALSASYYLSKKGLKVAIFEAKNEPGGGTWGGGMMFNELVVENDIKSFLDELGMNYLIKDNFIS
VDSVHFASSLLYNATKAGAVLFNNVIVEDIAFYENKVNGIVINWAPVIRQKLHVDPITIMAKFVVDGTGHPANVVNMLVDRGIDIDLPIGKIREYPM
NAKEGEKFVVENTKEVFPGLYVMGMAAVSVGGGPRMGPIFGGMIKSGLKVAKEILEKLSI

>Thermosipho melanesiensis (strain DSM 12029 / CIP 104789 / BI429)
MWDLEISKIIVNGFFEKFNDALDVDVAIVGGGPSALTASYFLTKNGFKVVIFEEKNDPGGGTWGGGMLFNELVVEEELEWMLKEFGMNYKRLNGFIS
IDSVHFASSLLYNTTKVGTKIFNNVIVEDILMEENRLCGVVINWAPVIKQRLHVDPITVKAKYVVDGTGHPASVVQMIIDRNLEVELPLDKIREFPM
NAKEGENFVLKNTKEVFPGLFVMGMAAVSVGGGPRMGPIFGGMLKSGEKVANAIVEKLSVEVSK

>Thermosulfidibacter takaii (strain DSM 17441 / JCM 13301 / NBRC 103674 / ABI70S6)
MLDEKIITKAIIESYTQNLLDYIDMDVAIVGAGPAGLTCAYYLAKEGVKVGIFERKLSIGGGIWGGGAMFNEIVLQEEALPIVQEMEVSYKPYKEKG
YYVINAVEFACALGLKAIRAGAKIFNLWSAIDVKVKGEDERVNGLVLLWTPVDTAGLHVDPITVEAKYVVDGTGHDAEIANVVVKKLKKKLATPTGD
VAGERPMWAEFGEKATEEFTGEVYPGLFVIGMAAVACYGKHRMGPIFGGMLLSGKKAAKMILECLK

>Thermosulfurimonas dismutans
MALDEVKITQAIVERFTEKLKEALELDVAVVGAGPSGLMAAYKLAKEGFKVAIFERKLSIGGGMWGGGMMFNEIVVQEEGARLLKEIGVEARPWQED
YYTADSVETVCALGYAARKAGAMIFNLISVEDVMVRKDRVVGLVINWTAVEMGGLHVDPLAIRSKYVVESTGHELSVLHIMQKKLGVQLMTPSGKIE
GEKSLWADVAETTTLENTREVFPGVFVAGMAANATFGSYRMGPIFGGMLLSGEKVAQEIAARLKK

>Thermosyntropha lipolytica DSM 11003
MVINDIKITRSIIEEYYAFTRDFLDCDVVIVGGGPAGMTAAYYTAQQGLRTVVLESRLSPGGGMWGGGMFFNQIVFQPEAGEILQELGISYTANREG
YLVVPSYRAVASLILAADRAGARILNGITAEDIMVRENRVCGVVINWTAAVKLGMHVDPLCIGGKVVIDATGHDAGIVRTYLDKSGGSLPEDEEERI
RTSSMWAAKGEEMVVEYTRFITEGLIACGMSVSSLFNTPRMGPIFGGMLFSGRKAAELALDYIRKVKA

>Thermovirga lienii (strain ATCC BAA-1197 / DSM 17291 / Cas60314)
MKLDEKIISKAIITRYYQKILSHLQVDVAIVGGGPSGLVAGYYLAKEGHRVALFERKLSVGGGMWGGGMLFNEIVVQEDAKEILEDFGVRVQPWEDA
GYYTADAIESVCSITSKAIQAGLTVFNCISVEDVSVEGDRITGLVINWTPVEMSGLHVDPLSIGASFVIDATGHDTEVVHMVAKKAPGKLMTPSGDI
EGEKFMCPDEAEKKTVENTKEVFPGLYVAGMACNATFGGPRMGPIFGGMLLSGRKVAALISQRLK

>Treponema sp. CETP13
MLEYNVSKGILDSYHTKLKSALDSDAIIVGSGPSGLVAGYFLAKAGKKVVMFERELAPGGGIWGGGMFFNDVMVQEEAATILSEIGVELPEVKDNFY
TIDSVYLASTLISKAVEAGVTLLNMISIEDIIFAKDESIGGVVLNWAPVHKEHMHVDPLMAISRCVLDATGHPSEIVNVLTRKNEITLNTKTGKVMG
ERSLKCKKAELATAENTCEIYPRLFVSGMAANGVAGAYRMGPVFGGMIRSGKKVAEQMLQCIDTEAPIYD

>Vulcanisaeta distributa (strain DSM 14429 / JCM 11212 / NBRC 100878 / IC-017)
MAGIYISESSITRAIMRSALKMLDEYSSVDVAIVGAGPSGMTAAYYLAKAGLKTIVLERRFSFGGGIGGAASHLPSIVVEYPASDILSKDFGVRLQD
MGDGLFAVDPAEMIAKLAVRAIDAGAKFLLGVHVDDVIIRDNPPRVAGLAVYWSTVQMAGVHTDPFFIEAKAVVDATGHDAEVAAVTTRKNPDLGLA
IHGEKSAHASVAEDLVVKYTGRVMEGLYVTGMAVAAVYGLPRMGPIFGSMIMSGKRVAELIINDLRR

>Zestosphaera tikiterensis
MEPLEAKISKIIWKETLNDWLKLSNVDVVVVGAGPSGMVTAKYLADSGIKTLVLERRLSFGGGIGGGGMLMHKVVVDSKALNILDDFKIKYSRSDYE
GLYVVDASELMAKLAAGAIDSGAKIVNGITVEDLIVRDNPFRVEGVVIQWSAVNLSGLHVDPLFIYSKAVVDATGHDAEVLKVLSRKNPEVNLKIPG
EKSAYAELSEELVVKHSGKVLPGLYVSGMAVAALYGIYRMGPIFTGMLLSGKKVAEEIAKDLRGGSQ
