## Supplementary Table 4 for "Structure and function of aerotolerant, multiple-turnover THI4 thiazole synthases"

**Supplementary Table 4  Occurrence of ROS defense genes in genomes of prokaryotes whose non-Cys THI4s have crystal structures**

The predicted proteomes in NCBI were searched by BlastP with the indicated query sequences. Significant hits (e-value < 1e-05) are indicated with a plus sign.

| Enzyme | Query[a] | Organism | | | |
|---|---|---|---|---|---|
| | | *Thermovibrio ammonificans* | *Methanococcus igneus* | *Methanococcus jannaschii* | *Methanothermococcus thermolithotrophicus* |
| Catalase-peroxidase | WP_013536945 | + | - | - | - |
| Cytochrome *c* peroxidase | WP_013537906 | + | - | - | - |
| Cytochrome *bd* complex | WP_013537252<br>WP_013537253 | + | - | - | - |
| Heme-catalase | WP_000077872 | - | - | - | - |
| Mn-catalase | WP_000488336 | - | - | - | - |

[a] GenBank identifier.
