## Supplementary Figures for "Structure and function of aerotolerant, multiple-turnover THI4 thiazole synthases"

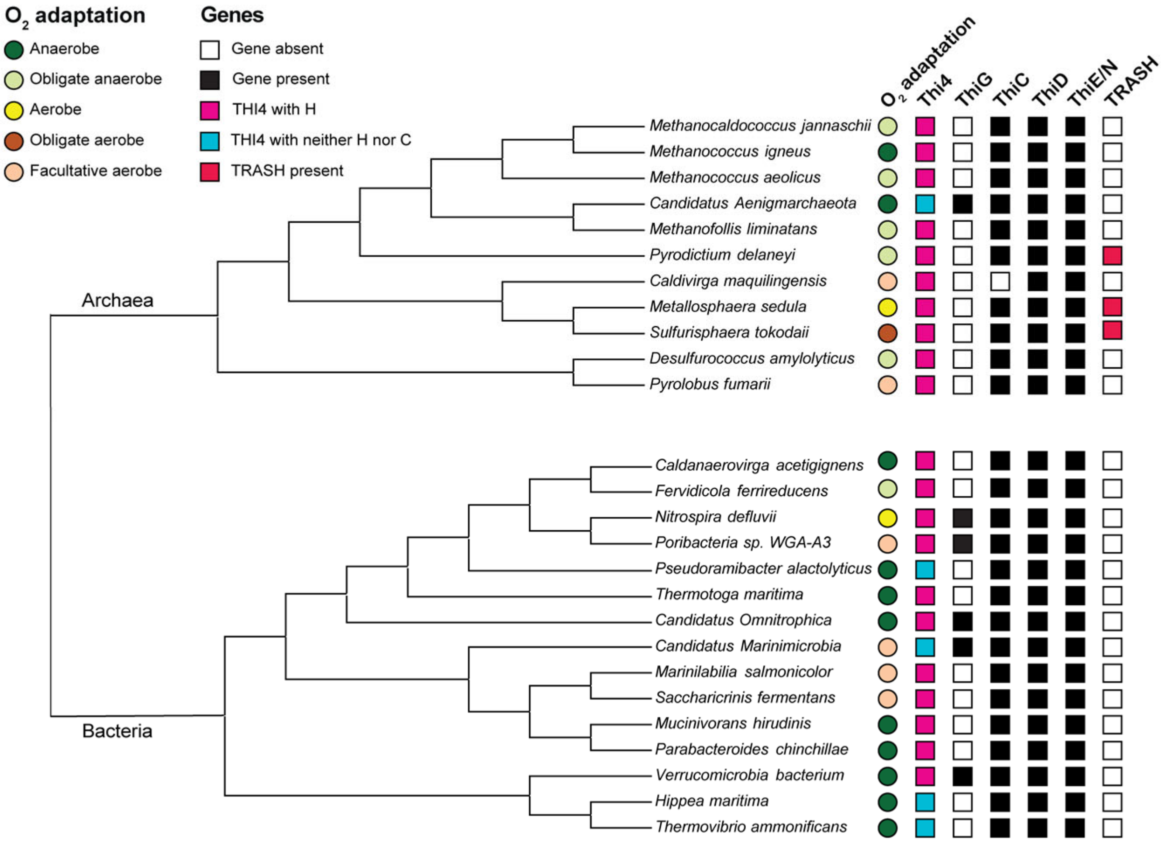


**Supplementary Figure 1. Ecology and genomic context of the 26 THI4s selected for testing.**

The first column (colored bullets) shows the $O_2$ adaptation of the 15 bacteria and 11 archaea whose THI4s were tested. The second column (colored squares) shows which residue replaces the active-site cysteine in each THI4. The next four columns (black or white squares) indicate the presence or absence of other thiamin synthesis enzymes (ThiG, ThiC, ThiD, and ThiE or ThiN). The last column (red or white squares) indicates presence or absence of a gene encoding a protein from the TRASH family (Trafficking, Resistance, And Sensing of Heavy metals) that is clustered with the THI4 gene.

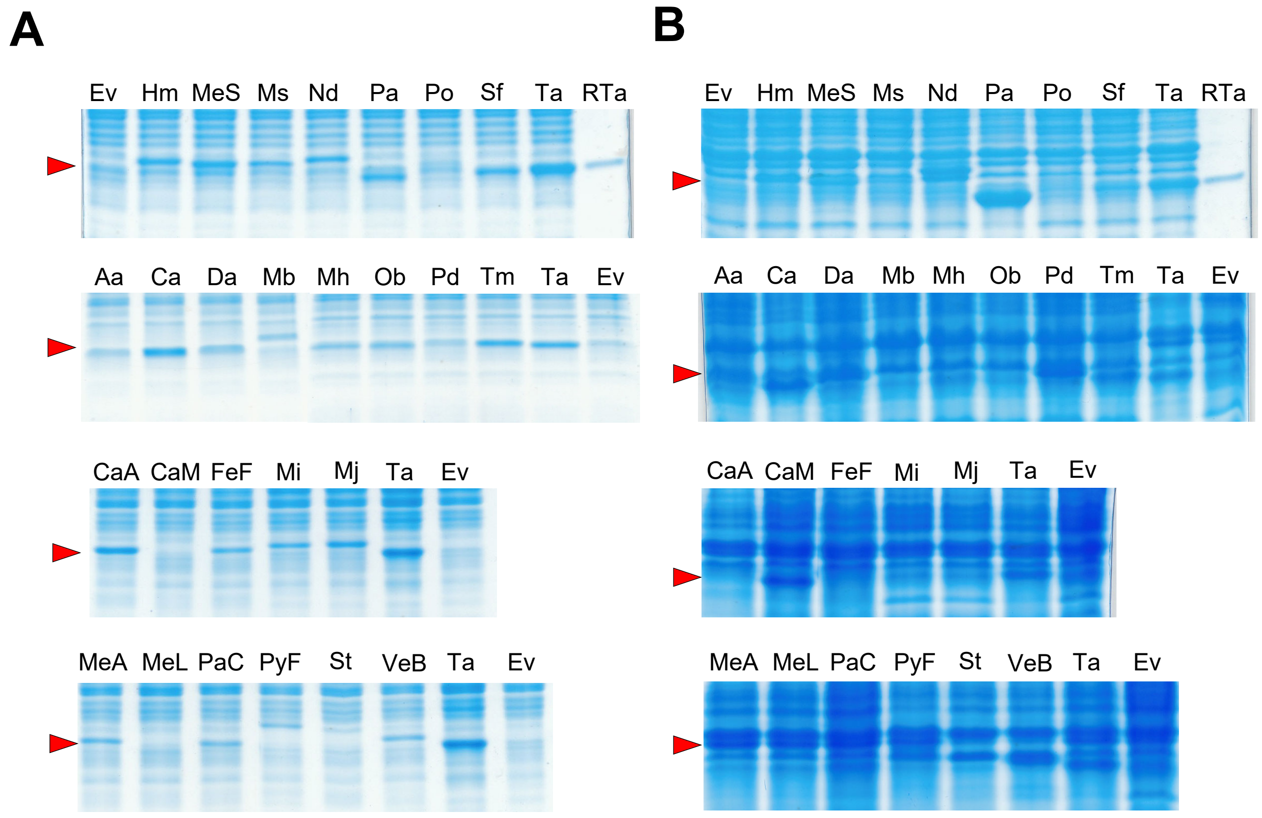


**Supplementary Figure 2. Soluble expression of non-Cys THI4s.**

Quantitative gel analysis of (**A**) soluble and (**B**) insoluble expression in *E.coli* of 26 selected non-Cys THI4s. Soluble and insoluble fractions of cells were run on 15% gels, stained with Coomassie blue, and scanned to quantify the THI4 band, for which purified recombinant *Thermovibrio ammonificans* THI4 (RTa, arrow) served as a marker. Organism abbreviations are as in Table 1.

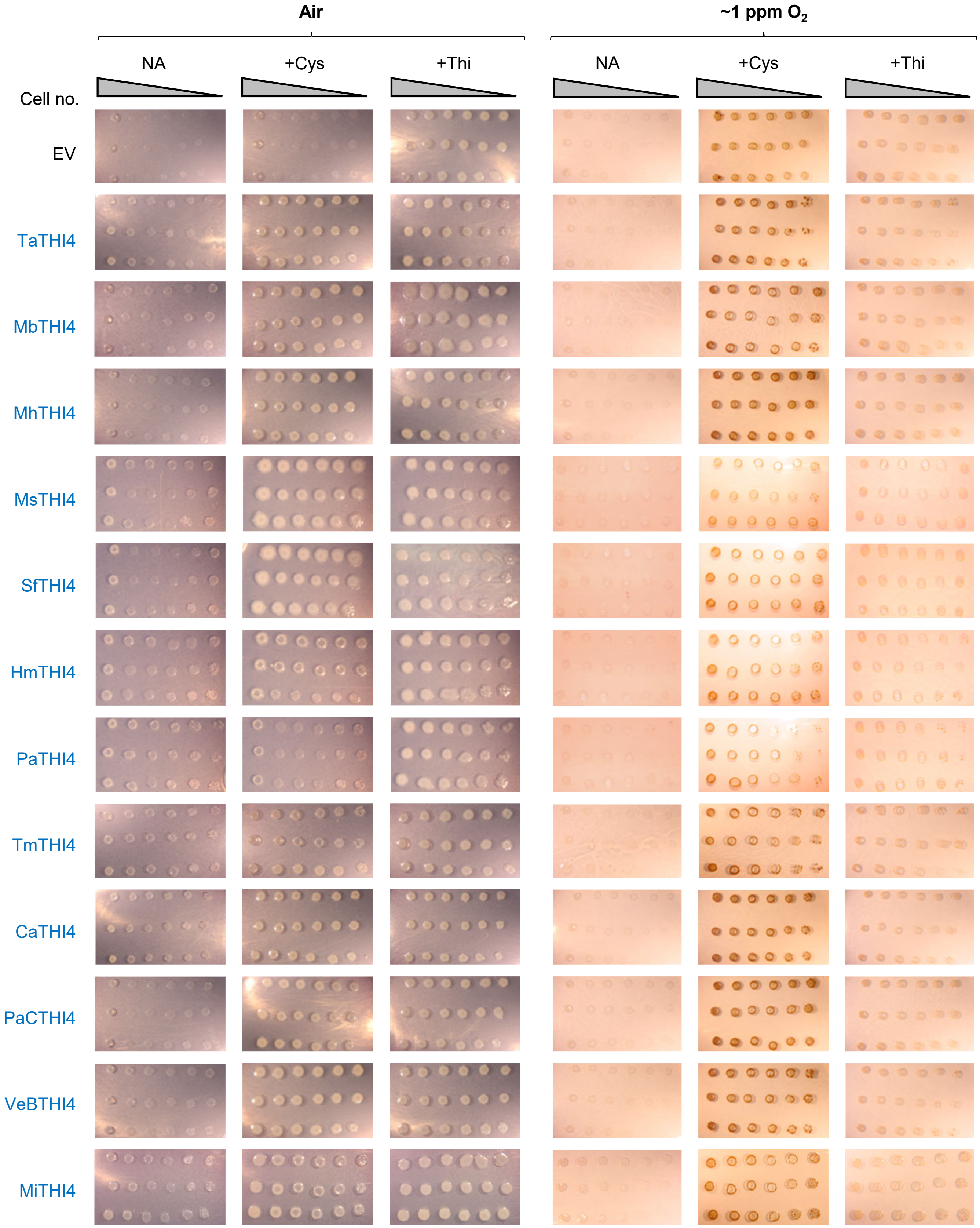

Air
~1 ppm $O_2$
NA
+Cys
+Thi
NA
+Cys
+Thi
Cell no.
EV
TaTHI4
MbTHI4
MhTHI4
MsTHI4
SfTHI4
HmTHI4
PaTHI4
TmTHI4
CaTHI4
PaCTHI4
VeBTHI4
MiTHI4

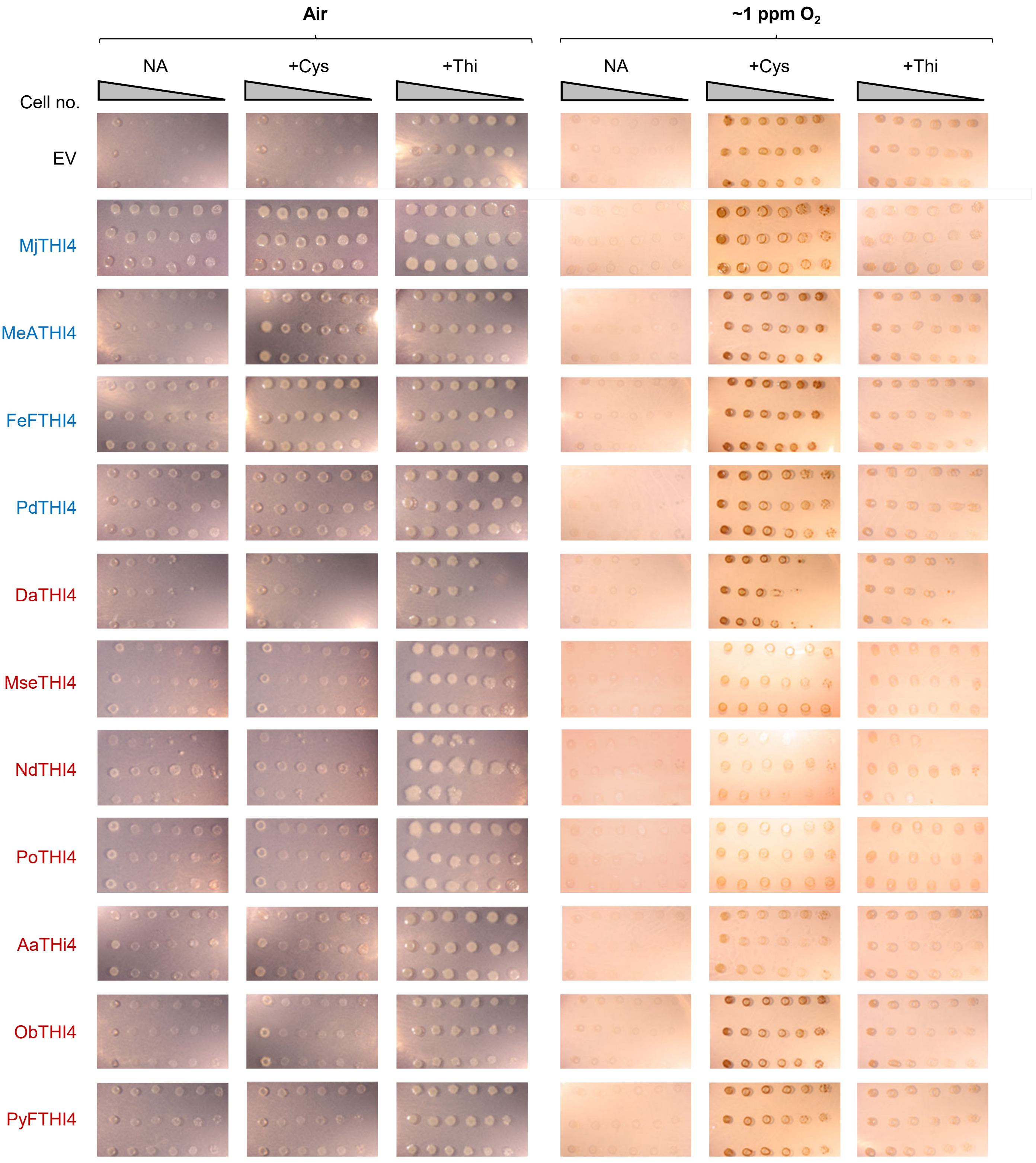


**Supplementary Figure 3. Functional complementation tests of non-Cys THI4s.**

Tests of functional complementation of an E. coli ΔthiG strain by all 23 soluble non-Cys THI4s or the empty vector (EV). Organism abbreviations are as in Table 1. Overnight cultures of three independent clones per construct were 10-fold serially diluted and spotted on plates of MOPS minimal medium containing 0.2% glycerol and 0.02% arabinose with no additions (NA) or plus 1 mM Cys or 100 nM thiamin. Cells were cultured in air or ~1 ppm O2. The medium used for culture in ~1 ppm O2 contained 40 mM nitrate. Images were captured after incubation at 37°C for 7 d. The high background in the ~1 ppm O2 +Cys treatment is staining of the inoculum cells. Organisms whose THI4s showed clear complementing activity in air, particularly with Cys supplementation, are blue; organisms whose THI4s did not show such activity are red. Note that complementing activity was scored from direct visual inspection of plates, not from the above images, which do not fully capture growth in every case.

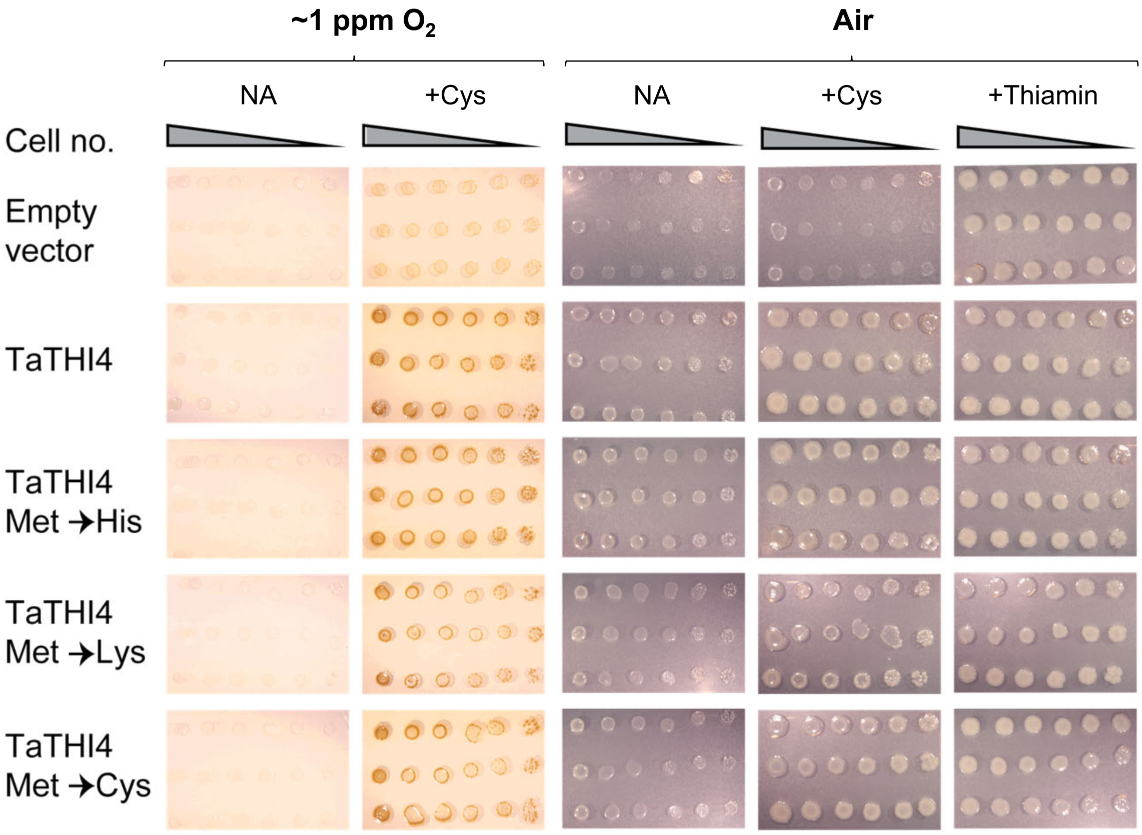


**Supplementary Figure 4. Complementation activity of TaTHI4 mutants**

An *E. coli* Δ*thiG* strain was transformed with empty vector or vector harboring wild type TaTHI4 or TaTHI4 with the indicated mutations of Met158. Overnight cultures of three independent clones per construct were 10-fold serially diluted and spotted on plates of MOPS minimal medium containing 0.2% (w/v) glycerol and 0.02% (w/v) arabinose with no addition (NA) or with1 mM Cys or 100 nM thiamin. The medium used for culture in ~1 ppm $O_2$ contained 40 mM nitrate. Cells were cultured in air or under $N_2$ containing ~1 ppm $O_2$. Images were captured after incubation at 37°C for 7 d.

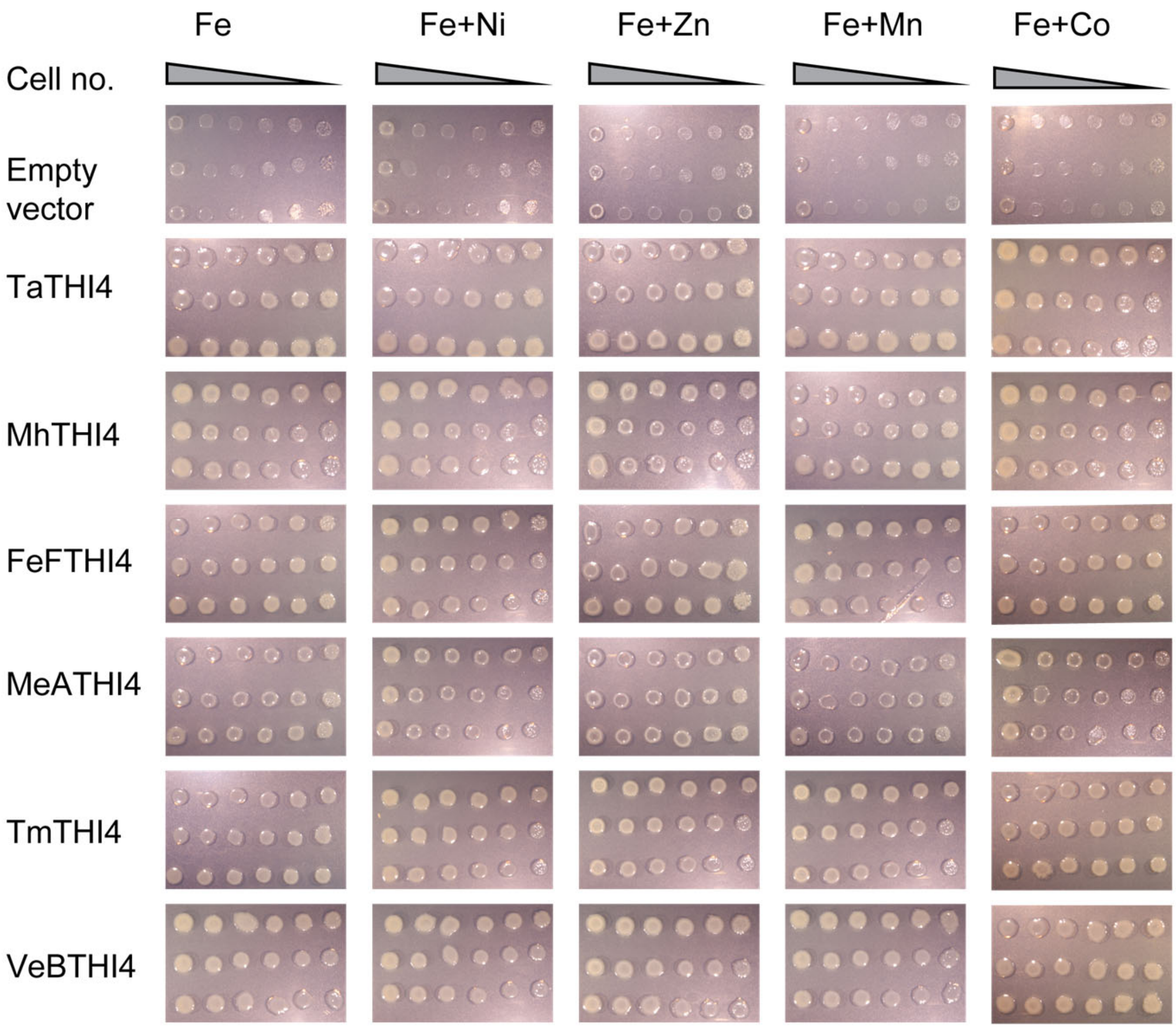


**Supplementary Figure 5 Effect of metal supplementation on complementing activity of THI4s**

An *E. coli* Δ*thiG* strain was transformed with empty vector or vector harboring the indicated THI4 sequence. Overnight cultures of three independent clones per construct were 10-fold serially diluted and spotted on plates of MOPS minimal medium containing 0.2% (w/v) glycerol, 0.02% (w/v) arabinose, 1 mM Cys, and 100 µM of the indicated metal. All media also contained the standard concentration of ferrous iron (100 µM). Cultures were incubated in air at 37°C for 7 d.

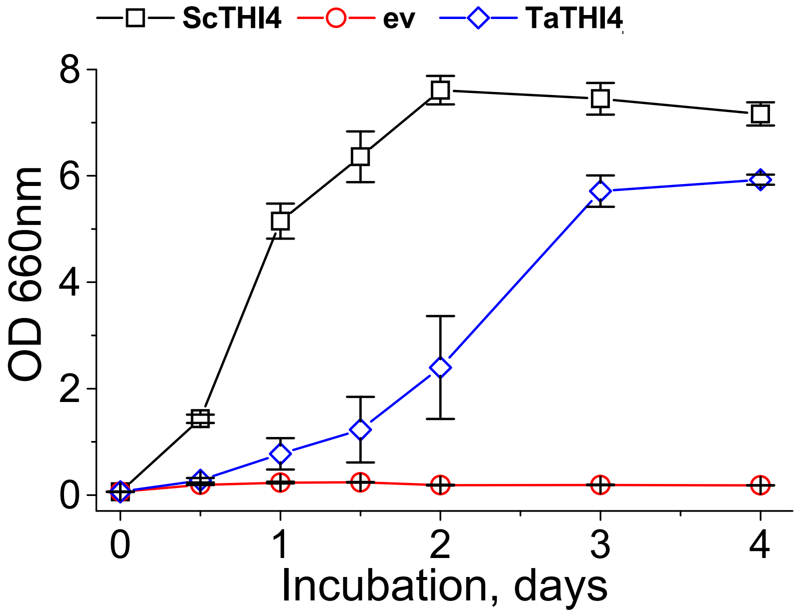


**Supplementary Figure 6  Complementation of a yeast Δ*THI4* strain by TaTHI4**

Cells of the Δ*THI4* strain transformed with empty vector (ev) or vector harboring TaTHI4 or yeast THI4 (ScTHI4) as positive control were cultured in thiamin-free SC minus histidine medium. Data are means of three or four independent clones ± S.E.
